# Versatile Computer Aided Design of Freeform DNA Nanostructures and Assemblies

**DOI:** 10.1101/2023.03.30.535006

**Authors:** Wolfgang G. Pfeifer, Chao-Min Huang, Michael G. Poirier, Gaurav Arya, Carlos E. Castro

**Affiliations:** Department of Mechanical and Aerospace Engineering, The Ohio State University, Columbus, OH 43210; Department of Physics, The Ohio State University, Columbus, OH 43210, USA; Department of Mechanical Engineering and Materials Science, Duke University, Durham, NC 27708; Interdisciplinary Biophysics Graduate Program, The Ohio State University, Columbus, OH 43210, USA; Department of Chemistry and Biochemistry, The Ohio State University, Columbus, OH 43210

## Abstract

Recent advances in structural DNA nanotechnology have been facilitated by design tools that continue to push the limits of structural complexity while simplifying an often-tedious design process. We recently introduced the software MagicDNA, which enables design of complex 3D DNA assemblies with many components; however, the design of structures with freeform features like vertices or curvature still required iterative design guided by simulation feedback and user intuition. Here, we present an updated design tool, MagicDNA 2.0, that automates the design of freeform 3D geometries, leveraging design models informed by coarse-grained molecular dynamics simulations. Our GUI-based, stepwise design approach integrates a high level of automation with versatile control over assembly and sub-component design parameters. We experimentally validated this approach by fabricating a range of DNA origami assemblies with complex freeform geometries, including a 3D Nozzle, G-clef, and Hilbert and Trifolium curves, confirming excellent agreement between design input, simulation, and structure formation.

## Introduction

Since its inception in the 1980s (*1*), structural DNA nanotechnology has found applications across a vast array of fields, including biosensing, nanoelectronics, gene and drug delivery, computing, optics, and plasmonics (*2-6*). The unique and exact molecular programmability inherent in the antiparallel and complementary base-pairing of double-stranded DNA enables the realization of nanoscale devices of high precision and geometric complexity. Additionally, the ability to integrate single- and double-stranded (ds) DNA with rigid bundles of dsDNA helices enables tailoring of both the dynamic and mechanical properties of these devices (*7-9*). This ability to precisely design structures with tunable stiffness and dynamics makes DNA nanotechnology highly suited for translating macroscopic mechanisms and machine- and materials-design concepts to the nanoscale. However, realizing advanced design concepts, such as compliant mechanisms and architected materials that contain intricate features like bends and vertices in closed- or open-loop 3D (“freeform”) geometries remains challenging, often requiring significant design iterations even for experts. Here, we introduce a new design algorithm and tool to automate the design of freeform structures, and we validate them both with current computational tools and experimental fabrication. Our results establish a powerful computer aided design (CAD) approach that will allow lay users to create complex 3D structures and assemblies without needing to learn the underlying molecular design concepts.

Scaffolded DNA origami is one approach particularly well-suited for the design of complex 3D DNA nanostructures (*10, 11*) In this approach, many short oligonucleotides (∼20–60 nucleotide [nt] long) called “staples” bind to a long (typically ∼7,000–8,000 nt long) single stranded DNA (ssDNA) termed “scaffold” to drive folding of the scaffold into a compact defined shape. The staples drive folding by binding to and bridging multiple distant contiguous sites of the scaffold to form dsDNA helices connected by migrationally immobile Holliday junctions that, if appropriately positioned, yield bundles of parallel dsDNA helices that can be arranged into a huge variety of 2D and 3D geometries.

As the DNA origami technique matured, CAD tools have become integral to facilitating a rational design process. A recent review article by Dey et al. (*12*) categorized available software tools for designing DNA origami structures into three generations. The first-generation design tools like caDNAno (*13*) and Tiamat (*14*) implemented graphical user interfaces (GUIs) to allow users to manually specify the routing and base-pairing relationship among DNA strands to generate staple strand sequence lists for folding. Second-generation design tools leveraged commercial CAD software to specify input geometries and developed algorithms to automate the underlying strand routings. These tools largely circumvent user inputs other than the original geometry, making structure design simpler for non-experts, but limiting the ability to tune local mechanical and dynamic properties (*15-21*). Lastly, third-generation software tools combine features from the first and second-generation tools to improve versatility for both expert and non-experts (*22, 23*). Within this realm, we recently introduced the tool MagicDNA (*24*) which combines the advantage of GUI and inherited routing algorithm to design complex DNA nanostructures.

Yet one limitation of all these CAD tools is that they inherently build up structures from straight segments of dsDNA helices or their bundles. Features like vertices and bends are achieved by coupling helices of different lengths together either to form bundles with angled edges (*i*.*e*., gradients) that can be connected to form a vertex or bundles that accumulate continuous bending stresses across their length to form curved features (*25, 26*). The integration of CAD tools with simulation tools (*27-32*) has been critical to enabling accurate design of these complex geometric features. In particular, the recently developed CAD tool MagicDNA facilitates integration with the oxDNA coarse-grained simulations (*30, 31, 33, 34*) to acquire 3D conformation feedback for design iteration. MagicDNA’s combination of graphical interfaces for 3D design manipulation and design parameter input, automated routing algorithms, and coupling to simulation to facilitate iterative design enables realization of complex multi-component assemblies with user control over local mechanical and dynamic properties. Additionally, the newly developed tool DNAxiS leverages simulation to design shapes with curvature, however currently limited to structures with revolved symmetry, either axisymmetric or periodic circular symmetry (*35*). However, even with these advanced design tools, achieving true freeform geometric designs is still challenging and often requires many iterations to tune the desired geometry.

To achieve true freeform design capability, we introduce here a simulation-guided algorithm for automated vertex and curvature design that we experimentally validate and implement in a new GUI in the MagicDNA package. Our algorithm takes sketched freeform spline curves as user input and converts these mathematical splines into physical DNA bundles. We introduce an analytical algorithm called *extrude* to automate the design of a vertex of defined vertex angle formed by the connection between two neighboring wedge-shaped bundles. This algorithm is informed by a series of oxDNA simulations of vertex joint designs that model the relationship between bending angle and vertex design parameters *(i*.*e*., edge gradients and bundle cross-section geometry). We also introduce an approach called *sweep* to design continuous, curved shapes in 3D from a series of subtly bending segments following similar simulation-based analytical models. These two algorithms are coupled to other useful features of MagicDNA such as 3D multi-component assembly, scaffold and staple routing algorithms, multi-scaffold design, and coarse-grained simulation feedback, providing a powerful platform for rapid design of freeform DNA architectures. To illustrate the versatility of our approach, we fabricated a range of 3D curved DNA origami structures designed using our platform and found excellent agreement between experiments and design predictions. Our results demonstrate outstanding control over the 3D geometry of DNA origamis through computer aided design and leverages the design versatility of integrated computer aided design and engineering through MagicDNA (*24*), allowing for rapid realization of complex DNA nanostructures, even by researchers from other fields.

## Results

### Overall approach: From freeform splines to DNA bundles

To realize freeform DNA origami structures, we developed a GUI where users can manually define their design using a series of points (Fig. 1a, b). These “control” points are connected either by straight discrete segments with the input points defining vertices (*extrude*), or by smooth splines, which are then broken into smaller segments with small relative angles to closely approximate the continuous curvature (*sweep*). Next, the mathematical straight-segment or smooth spline is converted into a DNA nanostructure of physical dimensions taking the dsDNA length and bundle cross-section into account (Fig. 1c). The cross-section can consist of any even number of duplexes in a square or honeycomb lattice. This conversion between conceptual lines to tangible DNA bundles involves the calculation of edge gradients for bending angles, local orientations for bundles, or non-linear duplex lengths within a bundle for continuous geometries (Fig. 1d, details discussed in extrude and sweep sections). Once DNA bundles are created with approximate positions and orientations in the assembly model (Fig. 1e), the connectivity matrix based on distances between connection sites serves as a high-level bridge between the user-defined bundle layout and the scaffold routing algorithm that connects all the DNA bundles via scaffold routing (Fig. 1f). The staple strand routing with frequent, periodic crossovers is mostly adapted from caDNAno (*13*), as in the original implementation of MagicDNA (*24*), to locally define the shape of the individual DNA bundles (Fig. 1g), with a few adaptions for continuous geometries and higher-order assembly through overhangs. Finally, oxDNA simulations (Fig. 1h) provide rapid feedback on the 3D conformation of the structure for fine-tuning the design before fabrication (Fig. 1i).

**Fig. 1.**
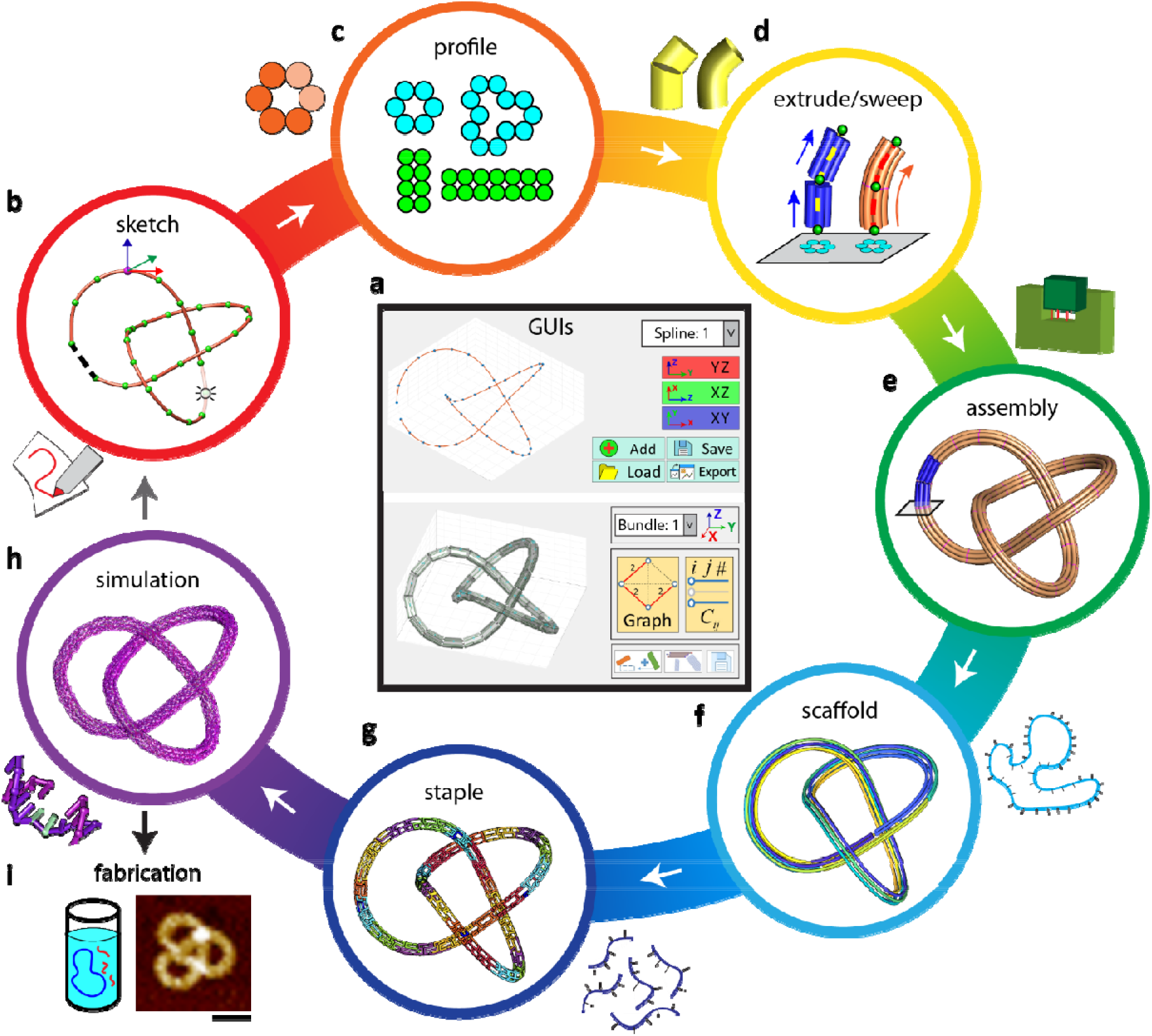
Flow chart of the key steps involved in the design of freeform DNA origami structures. (**a**) GUI of the software, showing the spline and bundle panel. (**b-h**) Steps involved in freeform design. (**i**) Experimental fabrication of the design for validation. The trefoil knot used here is purely for illustrative purposes: to show 3D spline curves and continuous geometries. In reality, this structure exhibited low experimental yields, likely due to kinetic traps arising from its unique topology. Scale bar = 50 nm.

### Extrude method for generating piecewise curved structures with vertices

The extrude tool allows users to connect individual bundle components with desired angles by automating the design of vertices (*i*.*e*., automatically specifying edge gradients) according to the defined input parameters (vertex angle and bundle cross-section). This involves the calculation of (1) bundle orientations (for projecting cross-section profiles along the helical axis direction) and (2) edge gradients for the two bending directions. The 3D orientation of each bundle is specified using two orthogonal unit vectors: one vector is normal to the cross-section profile (*i*.*e*., the helical axis) and the other points along the cross-section describing its rotation about the normal vector. Using a straight-line representation (connecting the control points of splines in a chain, Fig. 2a left), the algorithm takes the orientation of the first bundle as the reference frame and keeps propagating the orientations of the subsequent bundle relative to the prior bundle. Once all bundle orientations are identified, the user can define the cross-section for individual bundle components.

**Fig. 2.**
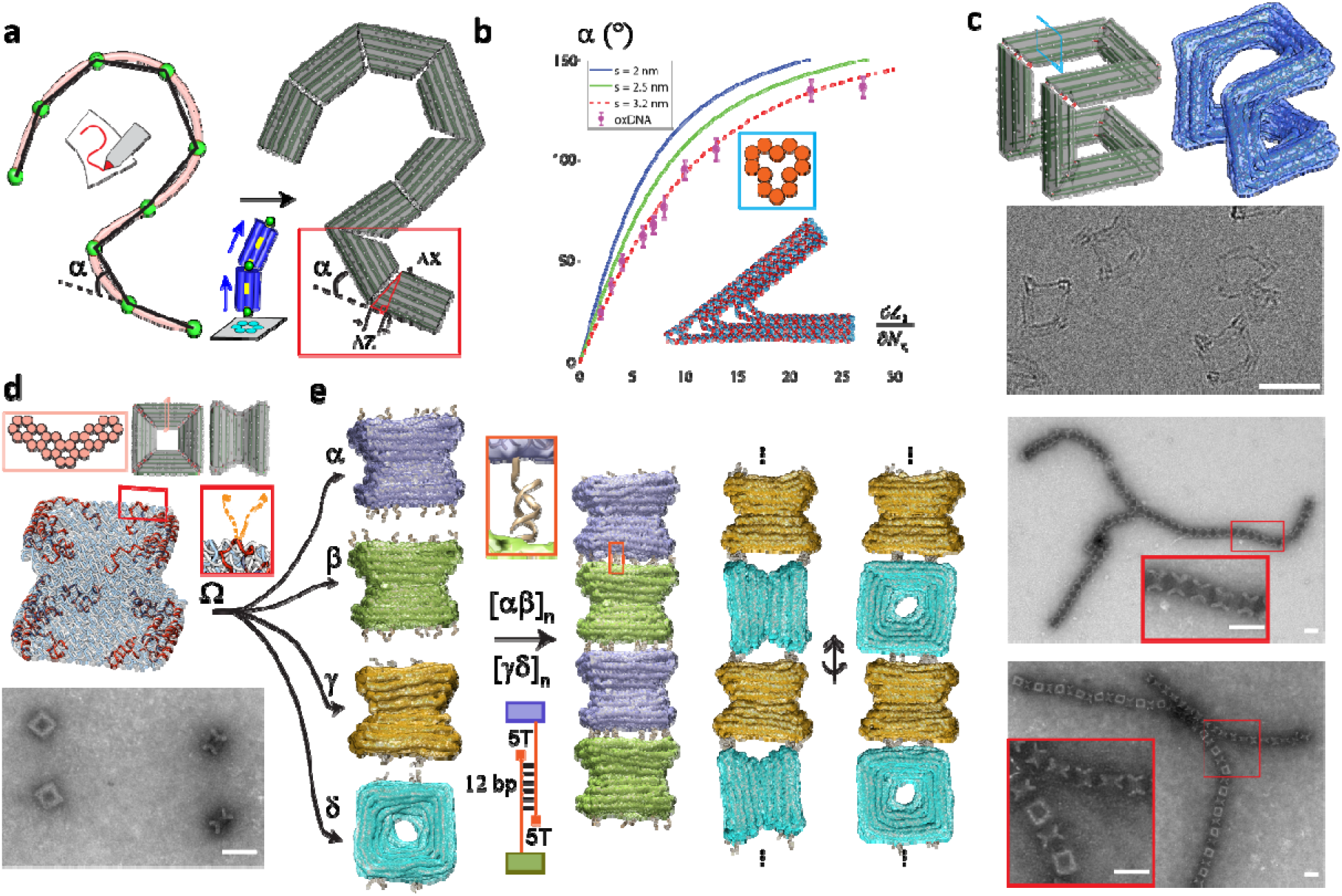
Extrude tool for creating piecewise curved structures from straight bundles which can be connected using overhangs to form larger arrays. (**a**) Conversion of a spline into a series of connected DNA origami bundle components with edge-gradients to approximate the target shape. (**b**) Single-joint oxDNA simulation to determine bending angles based on the input edge gradient (*i*.*e*., dZ/dX, dZ/dY) and cross-section. (**c**) Bundle model (left), oxDNA simulation average rendered with the outer surface (right), and cryo-EM image of the Hilbert structure (bottom). (**d**) From top to bottom: Cross-section, bundle model, helical routing model, and TEM image of the Nozzle structure. Inset shows overhang positions (orange). (**e**) oxDNA simulation averages with surface rendering of versions α, β, γ, and δ, showing the different overhang positions (left). The inset shows the formed, stable duplex of two complementary overhangs of the multimer [αβ]_n_. The sketch depicts our utilized overhang design, *i*.*e*., using a 5T spacer between the DNA origami interface and the sequence used in base-pairing to form the higher ordered structure. Results of the simulation of multimeric Nozzle filaments [αβ]_n_ and [γδ]_n_ and their respective, experimental validation. Inset show zoomed in structures. Scale bars = 50 nm.

The bending angle α at a vertex is related to the edge gradients of the two adjoining bundles on either side of the vertex, where the edge gradient is defined as the ratio of the difference in duplex lengths between neighboring layers of dsDNA helices to the layer, *i*.*e*., the center-to-center distance between duplexes in successive layers (Fig. 2b). Although one could utilize the geometry of DNA duplexes to derive an analytical relationship between the symmetric edge gradients and α, previous work has shown that the layer width is larger than the nominal 2 nm diameter of the DNA helix; the spacing between helices near vertices (*i*.*e*., edges of bundles) is likely to be even larger due to fraying effects (*27, 36, 37*). Thus, to evaluate the relationship between α and the edge-gradient parameters, we performed oxDNA simulations of a single DNA origami joint with four commonly used cross-sections (Fig. 2b, and Fig. S1). Our results show that larger cross-sections allow for more precise control of the bending angle. We also compared the simulation results to a geometric model of the vertex angle that depends on the difference in length between successive layers of dsDNA helices and the inter-helical spacing (Fig. S1). We found that an effective helical spacing of 3.2 nm near vertices best captures the vertex geometry (Fig. 2b). While prior work has found an effective inter-helical spacing of 2.1-2.4 nm (*38*) in DNA origami bundles, we attribute our slightly larger spacing to base pair fraying at the bundle edges commonly observed in regions with lower cross-over densities (Fig. S1). Therefore, we implemented this geometric vertex model with the 3.2 nm spacing to automate the vertex design process in our algorithm.

We implemented this vertex design model into the MagicDNA software tool and integrated it with the bundle location, orientation, and scaling calculations to automate the design of “extruded” 3D geometries consisting of straight segments with user defined length and cross-section connected by vertices. To illustrate the robustness of our automated extrude design approach, we designed and fabricated two structures: a Hilbert-curve structure with a 13-helix cross-section, which contains eight well-defined vertices forming a first order Hilbert-curve in 3D (Fig. 2c, top); and a Nozzle structure, which contains four well-defined vertices adjoining four bundles with a V-shaped cross-section to form a 3D nozzle geometry (Fig. 2d, top). Implementing common DNA origami protocols (*12, 39*), we realized high yields of properly folded structures, (Figs. 2c and 2d). TEM imaging revealed a homogeneous set of structures for both the Hilbert-curve and Nozzle designs. As the Hilbert structure has a more open geometry that could collapse upon surface deposition, we used cryogenic electron microscopy (cryo-EM) to reveal multiple orientations of this structure and confirm its 3D geometry. For the Nozzle structure, we observed various orientations in negative-stain TEM clearly illustrating different design features and validating successful folding.

We next used the Nozzle structure to demonstrate and validate an overhang design tool in MagicDNA, where ssDNA overhangs can be positioned at precise locations on the surface of the structure for connecting them into higher-order assemblies. We folded four versions (α, β, γ, and δ) of the Nozzle structure with unique overhangs. Versions α and β were designed with corresponding patterns of mutually complementary overhangs on the ends of the structure, while γ was designed with an overhang pattern on its ends that matches an overhang pattern on the side of δ. We then performed an ABAB type multimerization by mixing these structures. We realized two varieties of 1D filaments: one where both subunits are oriented similarly (Fig. 2e, [αβ]_n_) with the Nozzle axis aligned along the length of the filament, and another where the Nozzles alternate between aligned and perpendicular orientations with respect to the filament axis (Fig. 2e, [γδ]_n_). TEM imaging revealed proper multimerization of both filaments (Fig. 2e, right panel) and agarose gel-electrophoresis shows that multimerization works over a broad range of MgCl_2_ concentrations (Fig. S13). Gel electrophoresis analysis and additional TEM images are provided in Figs. S3 - 4, 6-12, and 14-17 for all extrude designs explored in this work.

### Sweep method for design of continuously curved structures

Although one could adopt the extrude method and keep adding control points in splines to form increasingly shorter bundles with more subtle bending angles for close to continuous curved geometries. A less tedious alternative is to exploit the smoothness of the splines to automatically sub-divide splines into DNA bundles that closely track the curved geometry. We refer to this approach as the sweep method (Fig. 3a). In this method, the algorithm treats the spline as a parametric curve, in terms of variable *s*, and takes the first derivative of the positions **p**(*s*) ≡ [*x*(*s*), *y*(*s*), *z*(*s*)] on the spline to obtain the unit tangent vector **ê**_*k*_(*s*) describing the normal vector of the cross-section profile in a continuous manner. Then, the algorithm calculates the second derivative of **p**(*s*) to obtain the unit normal vector **ê**_*i*_(*s*), a reference vector lying on the cross-section plane, and the binormal vector **ê**_*j*_(*s*) from the vector product **ê**_*k*_ × **ê**_*i*_ (Fig. 3a). Since the spline describes the centroid of the cross-section, and the 3D locations of each dsDNA duplex along the spline curve can be calculated by vector addition of the spline position **p** and linear combinations of **ê**_*i*_ and **ê**_*j*_. In this manner, the conformation of all duplexes in the target structure can be described using a single variable *s*, allowing creation of a continuous 3D model of the structure satisfying the inputs of the spline path and the desired cross-section profile. Through user-defined slicing (magenta slices in Fig. 3a right), the entire 3D model can be discretized into multiple DNA bundles of desired duplex lengths.

**Fig. 3.**
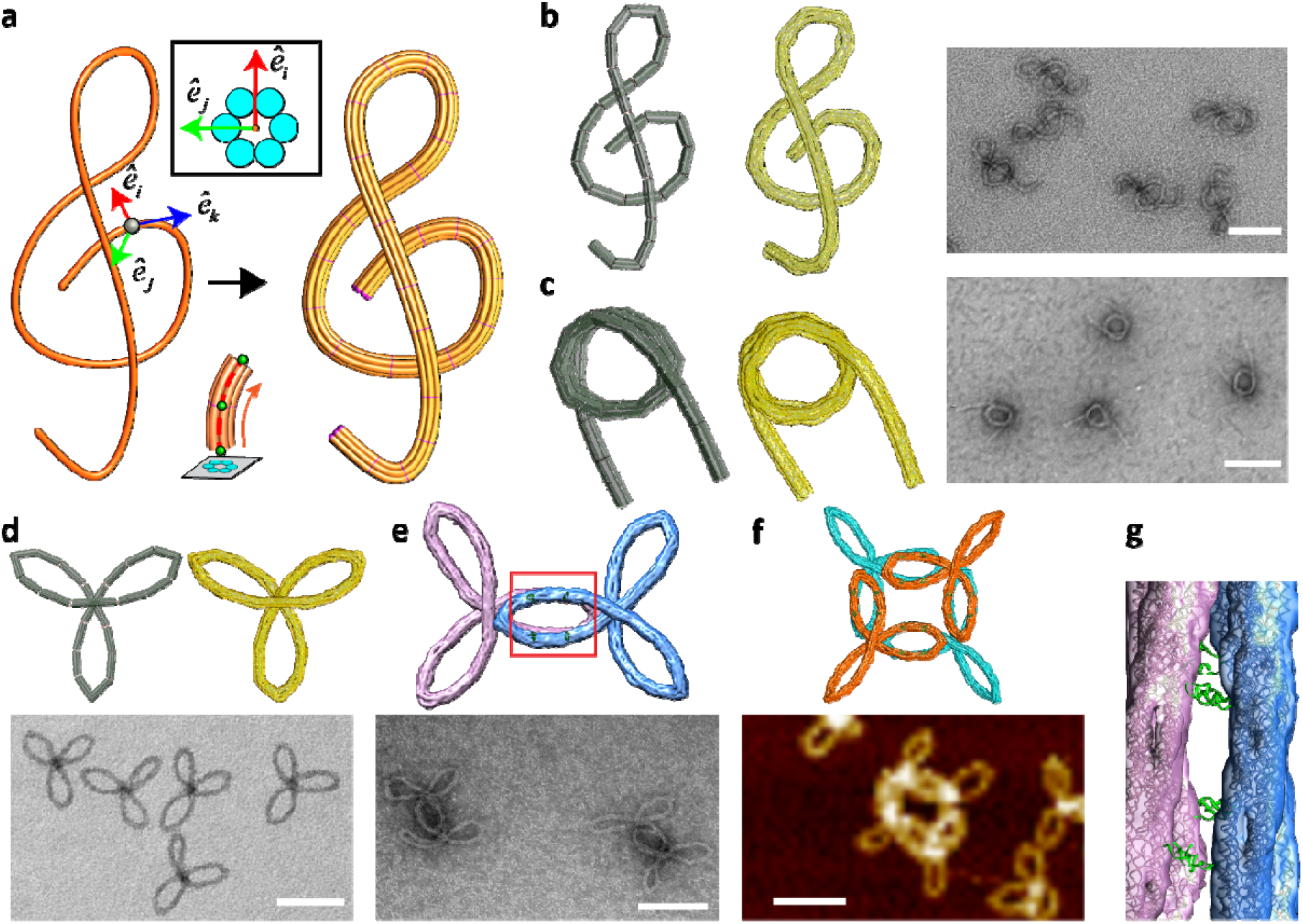
Sweep tool for designs with continuous freeform curvature. (**a**) Conversion of a mathematical spline into a bundle model (**b**) G-Clef structure (from left to right): Bundle model, oxDNA simulation, TEM image. (**c**) Nucleosome-like spring structure (from left to right): Bundle model, oxDNA simulation, TEM image. (**d**) Monomeric version of the Trifolium structure (bundle model and oxDNA simulation on top; TEM image on bottom). (**e**) Dimeric version of the Trifolium structure (oxDNA simulation on top; TEM image on bottom). (**f)** Closed-ring tetrameric version of the Trifolium structure (oxDNA simulation on top; AFM image on bottom). (**g**) Zoom-in on the red square from the overlapping arms in panel **e** showing the hybridization of overhangs on different bundles. Scale bars = 100 nm.

To validate the sweep algorithm, we designed and fabricated three distinct freeform structures, namely a G-Clef, a Nucleosome-like spring, and a Trifolium structure, each with a 6-helix bundle cross-section. Beginning with the G-Clef structure, Fig. 3a shows the conversion of the initial sketched spline into a continuous 3D bundle model, which was then discretized into 25 shorter components. The lengths of dsDNA duplexes in each segment were calculated by integrating the parametric curves for duplex axes between consecutive slice points (indicated by red lines in Fig. 3). The interfaces between consecutive bundles were held tight and parallel *via* multiple scaffold connections (typically zero bases long). We also implemented specific scaffold double connections between bundles 1 and 19, 6 and 21, and 11 and 18 (Fig. S18) to constrain the bundles into the G-Clef shape. OxDNA simulations revealed that the structure closely approximates the freeform G-Clef design (Fig. 3b, middle), and TEM images of the fabricated structures confirmed successful folding into the predicted structure (Fig. 3b, right). The Nucleosome-like spring was designed to form a 3D curve with 1.5 turns, forming a ∼60 nm nominal diameter core with ∼60-80 nm straight extensions on both sides (Fig. 3c). Simulations and images (Fig. 3c, middle) again revealed that the design accurately captures the desired 3D freeform geometry. Lastly, we designed the Trifolium design and showed successful folding of the structures according to design predictions (Fig. 3d). As in the case of the Nozzle structure, we used the Trifolium structure to demonstrate integration of sweep tool with the overhang tool for higher-order assembly. To this end, we designed dimeric and tetrameric assemblies of the Trifolium structures using the overhang tool and confirmed successful fabrication of both assemblies using simulations and imaging (Fig. 3e–3g). Gel electrophoresis analysis and additional TEM images are provided in Figs. S19–20, 22–23, and 31–36 for all sweep designs.

### Integrating with MagicDNA features to expand design space

The extrude and sweep approaches allow for the automated design of complex 3D geometries. Implementing this freeform design automation in MagicDNA allows users to leverage other features of this software to expand on freeform design. Here, we expand on freeform designs by leveraging two specific features of MagicDNA: 1) control over components-level cross-section, and 2) versatile scaffold routing algorithm including multi-scaffold design. We used the extrude approach to design a Crown structure resembling the symbol for the Queen chess piece (Fig. 4a). We carried the line model design (Fig. 4a, left) through the MagicDNA design workflow to assign distinct cross-sections to individual components, including a 6 helix-bundle honeycomb cross-section to the spikes of the crown, an 8 helix-bundle square lattice cross-section to the base, and a 2-helix cross-section to the struts that connect the base to the spikes (Fig. 4a, middle). The versatile scaffold routing algorithm and assembly operations of the MagicDNA framework allowed us to place scaffold connections to the base and the outer frame on either side of the struts to stabilize the overall geometry. OxDNA simulations (Fig. 4a, right) and TEM images (Fig. 4b) revealed that the design accurately captured the target geometry and that the structure folded efficiently.

**Fig. 4.**
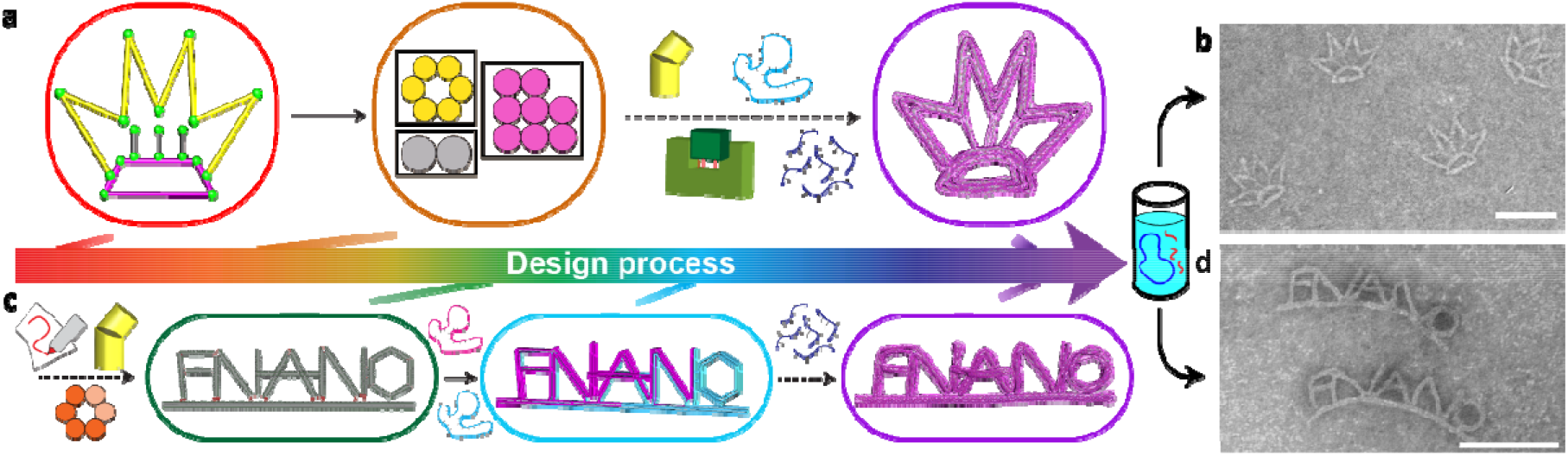
Combining freeform design with variable cross-section and multi-scaffold features of MagicDNA. (**a**) The Crown design workflow starts with the line model comprising multiple splines totaling 15 bundle components. Three different cross-sections were assigned to the yellow (crown spikes, 6 helix bundle), pink (base, 8 helix bundle), and gray (struts, 2-helix bundles) components. Components were assembled followed by automated scaffold and staple routing, and the final design was simulated in oxDNA. (**b**) TEM images illustrate well-folded Crown structures. (**c**) FNANO design workflow follows similar steps, with the bundle model and multi-scaffold routing highlighted, culminating in the final design simulated in oxDNA. (**d**) TEM images reveal well-folded FNANO script structures. Scale bars = 100 nm.

These advances in design capability are synergistic with recent advances in fabrication methods, namely the ability to incorporate multiple orthogonal sequence scaffolds to fold larger structures in a single-pot folding reaction (*40*). We previously implemented multi-scaffold design into MagicDNA. To demonstrate integration of multi-scaffold design with freeform automation, we designed an FNANO-script structure comprised of five letters that are assembled and connected to a stiff support bundle (Fig. 4c). All letters have an 8-helix-bundle cross-section, and the support bundle has a 6-helix-bundle cross-section. The multi-scaffold routing algorithm in MagicDNA automatically divided the design into two scaffolds with the structure formed from a total of 14,778 base pairs. Successful realization of the FNANO structure is shown in Fig. 4d. Gel electrophoresis analysis and additional TEM images are provided in Figs. S25–26 and 28–29 for structures with different cross-sections and multiple scaffolds.

Finally, we highlight the capability of the automated freeform design implemented in MagicDNA with an assorted gallery of sophisticated DNA origami nanostructures presented in Fig. 5 including numbers, lowercase, uppercase, and Greek letters, parametric curves, chess pieces, and other 3D freeform designs, demonstrating the versatility of our design approach and software tool. All designs were simulated in oxDNA, and the molecular structure of the average configuration is overlaid with a semi-transparent surface view that envelops the maximum conformational fluctuations, giving an indication of the local flexibility and the overall structural stability. As expected, structures with open topology, small cross-sections (*e*.*g*., 6-helix-bundles), and lengthy components display larger fluctuations around the mean configuration *(e*.*g*., the letter c) (Fig. S43).

**Fig. 5.**
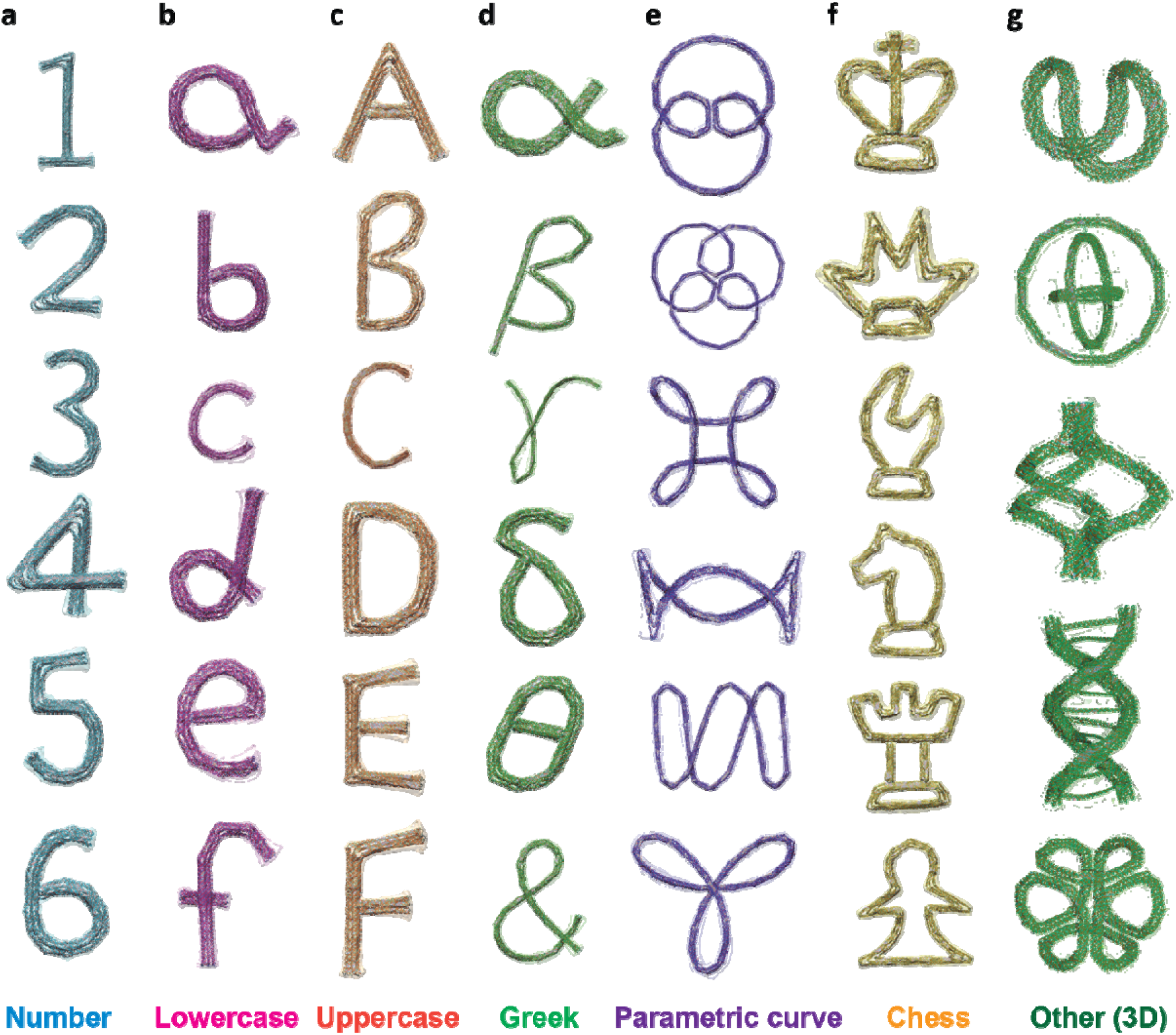
Design gallery of freeform structures created using our software. Conformations of additional freeform designs obtained using oxDNA simulations. The designs include: (**a**) numbers, (**b**) lowercase letters, (**c**) uppercase letters, (**d**) Greek letters, and the ampersand symbol, (**e**) parametric curves, (**f**) six chess pieces, and (**g**) other 3D structures. Each structure is shown by its mean conformation computed from the simulation with a bounding surface representing the standard deviation of fluctuations the structure exhibits in the simulation.

## Discussion

### Design philosophy

We present a versatile design framework that interweaves GUIs (Fig. 1a) with algorithmic automations to enable the design of arbitrary freeform DNA origami structures and assemblies based on user-defined design parameters. The design process starts with the user sketching the geometry of the envisioned structure as freeform dimensionless splines by taking advantage of a GUI using which one can define and manipulate the shape of the splines and visualize them in real time. This real-time sketching allows users to rely on gradual modifications to achieve the target design. Realizing the 3D form of the structural design requires additional inputs such as cross-section profiles and edge gradients. Typically, the structures are comprised of many dsDNA bundles, which would make manual input of bundle design parameters tedious and error prone. To address this problem, we implemented two new algorithms: the *extrude* and *sweep* methods. Both algorithms rely on a series of oxDNA simulations that allowed us to derive a geometric model for relating bundle design parameters to bending angle, which was implemented as an algorithm to automatically convert the spline model to the assembly model (*i*.*e*., geometry comprised of connected bundles) in MagicDNA. Having the full sketch and assembly design process implemented in one software (unlike second-generation design tools that import geometry models from exterior CAD software) allows for flexible iteration between the sketch and the assembly step to rapidly define and tune the geometry, while still leveraging the algorithms implemented in MagicDNA for scaffold and staple strand routing to iterate and complete designs. These designs can be evaluated computationally by coarse-grained molecular dynamics simulations such as oxDNA with automatically generated simulation input files to achieve an iterative robust design framework before moving forward to experimental fabrication.

### Versatile design framework and hierarchical assembly

We sacrificed full automation of the design process to allow users to customize structures for intended applications. For example, the same geometry can be realized with different bundle cross-sections to modulate the structure stiffness, as demonstrated with the DNA crown structure (Fig. 4a). Furthermore, a semi-automatic software allows the user to examine the design status at several stages, modify details from default settings, and seamlessly move forward and backward through the design steps, all of which is facilitated by rapid feedback in GUIs to visualize results of design choices. Furthermore, the new design paradigm introduced here can fully leverage the growing library of scaffold sources (*40-43*), removing restrictions in spatial dimensions and allowing users to realize even more diverse structures. For example, we used the multi-scaffold feature of MagicDNA to achieve large structures, such as the “FNANO” script (Fig. 4c), which was designed from two scaffolds with orthogonal sequences (*42*) to honor this annual conference that has played a seminal role in fostering the field of DNA nanotechnology.

### Expanded design scope with freeform features

The design algorithms and associated software and GUIs developed here significantly expand the capability of the MagicDNA design framework for designing freeform, curved features. Our results demonstrate a vast array of freeform designs with excellent agreement between oxDNA simulations and experiments. Apart from validating our design strategy, these results also illustrate the importance of coarse-grained simulations to computationally support the immense design space available for freeform structures with customizable features, allowing users to control both geometry and properties. In particular stiffness can be increased by using larger cross-sections (which might require a multi-scaffold design), or, for cases with overlapping components, introducing inter-bundle connections. For example, the bottom structure in the “Other 3D” series (Fig. 5) where a continuous freeform 6HB routes through the six faces of a cube (*44*) includes six inter-bundle connections along its edges. One could also exploit the versatility of the semi-automatic framework to design dynamic devices with multiple components, connected by single-stranded scaffold segments, to engineer pre-defined motions, as shown in the gyroscope structure (Fig. 5g, second from top) where two rotational degrees of freedoms exist between the inner and outer rings.

The wide range of DNA origami designs presented illustrate the realization of advanced freeform features within the general categories of static, compliant, and dynamic DNA nanostructures. The freeform design algorithms integrate seamlessly with other features in MagicDNA such hierarchical assembly and multi-scaffold designs to further broaden the design space for DNA self-assembly and provide a foundation for the realization of advanced materials, including assemblies of compliant mechanisms and meta-materials.

## Materials and Methods

### Materials

Oligonucleotides for folding of the DNA origami structures were purchased at 10 nmol synthesis scale from Eurofins Genomics (https://eurofinsgenomics.com) or at 25 nmol synthesis scale from Integrated DNA Technologies Inc. (www.IDTDNA.com). DNA oligonucleotides were purchased with salt-free purification at 100 µM concentration in RNAse free water and were used without further purification. The single-stranded scaffold (p8064) was prepared as described previously (*45*), the single-stranded scaffold (CS03) was purchased from tilibit (https://www.tilibit.com/).

### DNA Origami folding and purification

Folding of freeform DNA origami structures was performed by mixing 10 nM scaffold DNA with 100 nM corresponding staple strands in TEMg buffer (Tris: 5 mM, EDTA: 1 mM, MgCl_2_: 10 mM, pH 8). Samples were folded in a BioRad PCR cycler by first heating them up to 65 °C for 15 minutes and then cooling them down to 20 °C over the course of 14 hours. The Nozzle-structure was folded over 2.5 days. The detailed protocols for thermal annealing and all staple strand sequences are provided in the Supplementary Material. Purification of DNA origami structures was performed by gel extraction (Freeze ‘N Squeeze, BioRad) or PEG precipitation (*46*). Hierarchical assembly of the Trifolium structure was performed by first folding monomers with different sets of overhangs, purifying them individually, mixing them in an equimolar ratio and incubating them at temperatures between 40 and 55 °C for 20 hours. Unless stated otherwise, the Magnesium concentration for these hierarchical assemblies was adjusted to 20 mM. Hierarchical assembly of the Nozzle multimers was performed by folding structures with different sets of overhangs, purifying them individually, mixing them in an equimolar ratio, and incubating them at 40 °C for 20 hours.

### AFM imaging

Gel purified structures (around 1 nM) were used for AFM imaging by adsorbing 6 µl of sample onto freshly cleaved mica (V1, Ted Pella). After three minutes of incubation, the mica was rinsed carefully with milliQ-H_2_O and dried with a gentle flow of air. Samples were subsequently imaged in ScanAsyst Mode using a Bruker BioScope Resolve microscope equipped with a Nanoscope V controller. ScanAsyst Air probes (Bruker) with a nominal spring constant of 0.4 N/m were used for scanning. Height information was recorded in the retrace channel.

### Negative stain transmission electron microscopy

Purified DNA origami structures (1–10 nM) were adsorbed onto freshly glow discharged copper grids (Electron Microscopy Sciences, Hatfield, PA) and incubated for 4 minutes. Excess sample solution was subsequently wicked off with filter paper (Whatman #4) and the grid stained with two 6 µl drops of 1% aqueous Uranyl acetate (SPI Supplies) solution. Samples were dried for at least 30 minutes prior to imaging. Imaging was performed on a FEI Tecnai G2 Spirit operated at 80 kV acceleration.

### Cryo-electron microscopy

The Hilbert structure used for Cryo-EM analysis was subjected to two rounds of PEG precipitation to ensure sufficient purity. Furthermore, the second round was used to increase the DNA origami concentration to 500 nM. Glow discharged grids (Ted Pella) were used with a Thermo Scientific Vitrobot at 22 °C, 0 s drain time, 3 s blot time, 0 blot force at 100% humidity. Imaging was done on a Thermo Scientific Glacios cryo-TEM, equipped with a Falcon III direct electron detector and 200 kV x-FEG.

### Coarse-grained MD simulations

For performing oxDNA2 simulations, the relevant topology and initial configuration files were generated using the refined version of MagicDNA developed here. The initial configuration was relaxed in a manner similar to our previous study. Briefly, this involved substituting the DNA back-bone potential with linear springs, gradually increasing the force constants of these springs, and then applying mutual traps between paired scaffold and staple bases over a period of 100,000 timesteps. Next, the backbone potential was restored and a further 1 million timesteps of simulations were carried out, still retaining the mutual traps to finalize the relaxation process. During both these relaxation steps, we used a small time step of 3.03 fs. Finally, we removed the mutual traps and continued the simulation for an additional period of 20 million time steps of size 15.15 fs for calibrating the edge gradients at bundle vertex, and 10 million time steps of size 3.03 fs for the other simulations providing feedback on structure conformation for each design. We used a John thermostat (with diffusion coefficient and Newtonian step settings of 2.5 and 103) to maintain a constant temperature of 30 °C. A monovalent salt concentration at 0.5 M was chosen to mimic standard Mg-induced folding conditions. GPU acceleration was used whenever available. The trajectory files were analyzed using functions built into MagicDNA, including visualization of configurations and calculation of root-mean-square deviation (RMSD) and root-mean-squared fluctuations (RMSF). The average configurations were exported to the UCSF Chimera software and rendered to obtain high-quality images. We typically report the mean configuration of each structure and a surface map enveloping its conformational fluctuations.

## Acknowledgments

We acknowledge resources from the Campus Microscopy and Imaging Facility (CMIF) at The Ohio State University for negative stain TEM imaging. Cryo-electron microscopy was performed at the Center of Electron Microscopy and Analysis (CEMAS) at The Ohio State University. We are thankful to Prof. G. Agarwal (The Ohio State University) for providing access to the Bruker BioScope Resolve AFM. We gratefully acknowledge Hendrik Dietz for providing the CS03 scaffold.

## Author contributions

WGP performed all experiments. CMH wrote the MatLab code and performed oxDNA simulations. WGP and CMH wrote the initial draft of the manuscript and prepared the figures. All authors discussed results. All authors reviewed and edited the final manuscript.

