## Supplementary Material for "Versatile Computer Aided Design of Freeform DNA Nanostructures and Assemblies"

**This PDF file includes:**

Supplementary Text Folding Protocols

Figure S1: Simulation of bending angles as function of edge gradients, using oxDNA.
Figure S2: Design details of the Hilbert structure.
Figure S3: Magnesium screening during the folding of the Hilbert structure.
Figure S4: TEM micrograph the Hilbert structure.
Figure S5: Design details of the Nozzle structure.
Figure S6: Magnesium screening during the folding of the Nozzle structure.
Figure S7: TEM micrograph of the Nozzle structure (no overhangs).
Figure S8: Gel electrophoretic analysis of Nozzle variants with different overhangs.
Figure S9: TEM micrograph of the Nozzle structure (overhang species α).
Figure S10: TEM micrograph of the Nozzle structure (overhang species β).
Figure S11: TEM micrograph of the Nozzle structure (overhang species γ).
Figure S12: TEM micrograph of the Nozzle structure (overhang species δ).
Figure S13: Gel electrophoretic analysis of the multimerization of Nozzle structures with overhangs.
Figure S14: TEM micrograph of multimeric Nozzle structure (overhang species α & β), incubated in the presence of 10 mM $MgCl_2$.
Figure S15: TEM micrograph of multimeric Nozzle structure (overhang species α & β), incubated in the presence of 20 mM $MgCl_2$.
Figure S16: TEM micrograph of multimeric Nozzle structure (overhang species γ & δ), incubated in the presence of 10 mM $MgCl_2$.
Figure S17: TEM micrograph of multimeric Nozzle structure (overhang species γ & δ), incubated in the presence of 20 mM $MgCl_2$.
Figure S18: Design details of the G-clef structure.
Figure S19: Magnesium screening during the folding of the G-clef structure.
Figure S20: AFM image of the G-clef structure.
Figure S21: Design details of the nucleosome-like spring structure.
Figure S22: Magnesium screening during the folding of the nucleosome-like spring structure.
Figure S23: TEM micrograph of the nucleosome-like spring structure.
Figure S24: Design details of the crown structure.
Figure S25: Magnesium screening during the folding of the crown structure.

Figure S26: TEM micrograph of crown structure.
Figure S27: Design details of the FNANO-script structure.
Figure S28: Magnesium screening during the folding of the FNANO-script structure.
Figure S29: TEM micrograph of the FNANO-script structure.
Figure S30: Design details of the Trifolium structure.
Figure S31: Magnesium screening during the folding of the trifolium structure.
Figure S32: AFM image of the Trifolium structure.
Figure S33: Multimerization of Trifolium structures with overhangs Species A(1+2) and overhangs Species B(1) or Species B(2).
Figure S34: AFM image of the Trifolium dimer-structures.
Figure S35: AFM image of higher order Trifolium structures.
Figure S36: AFM image of higher order Trifolium structures.
Figure S37: Other freeform examples in the number series (0 & 1).
Figure S38: Other freeform examples in the number series (2 & 3).
Figure S39: Other freeform examples in the number series (4 & 5).
Figure S40: Other freeform examples in the number series (6 & 7).
Figure S41: Other freeform examples in the number series (8 & 9).
Figure S42: Other freeform examples in the lowercase series (a & b).
Figure S43: Other freeform examples in the lowercase series (c & d).
Figure S44: Other freeform examples in the lowercase series (e & f).
Figure S45: Other freeform examples in the lowercase series (g & h).
Figure S46: Other freeform examples in the lowercase series (i & j).
Figure S47: Other freeform examples in the uppercase series (A & B).
Figure S48: Other freeform examples in the uppercase series (C & D).
Figure S49: Other freeform examples in the uppercase series (E & F).
Figure S50: Other freeform examples in the uppercase series (G & H).
Figure S51: Other freeform examples in the uppercase series (I & J).
Figure S52: Other freeform examples in the symbol series (alpha & beta)
Figure S53: Other freeform examples in the symbol series (gamma & theta).
Figure S54: Other freeform examples in the symbol series (ampersand & delta).
Figure S55: Other freeform examples in the symbol series (sigma & omega).
Figure S56: Other freeform examples in the symbol series (psi).
Figure S57: Other freeform examples in the parametric curve series (I).
Figure S58: Other freeform examples in the parametric curve series (II).
Figure S59: Other freeform examples in the parametric curve series (III).
Figure S60: Other freeform examples in the chess series (Queen & King).
Figure S61: Other freeform examples in the chess series (Bishop & Knight).
Figure S62: Other freeform examples in the chess series (Rook & Pawn).
Figure S63: Other freeform examples in the 3D series (Baseball-seam & Gyroscope).
Figure S64: Other freeform examples in the 3D series (Twist stair railing & Duplex DNA).
Figure S65: Other freeform example in the 3D series (3D continuous curve).

**Folding protocols**
Folding of freeform DNA origami structures was performed with the following protocol.

All structures (except the nozzle) were folded with the following14hour protocol.

| T [°C] | t [min] |
|---|---|
| 65 | 15 |
| 64-61 | 3 |
| 60 | 5 |
| 59-58 | 10 |
| 57 | 15 |
| 56 | 25 |
| 55 | 30 |
| 54 | 45 |
| 53-49 | 60 |
| 48-45 | 42 |
| 44 | 36 |
| 43-42 | 32 |
| 41-39 | 20 |
| 38 | 15 |
| 37 | 10 |
| 36-35 | 5 |
| 34-30 | 2 |

All variants of the nozzle structure were folded with the following, extended folding protocol.

| T [°C] | t [min] |
|---|---|
| 70 | 15 |
| 65-62 | 60 |
| 61-59 | 120 |
| 58-46 | 180 |
| 45-40 | 60 |
| 39-25 | 30 |

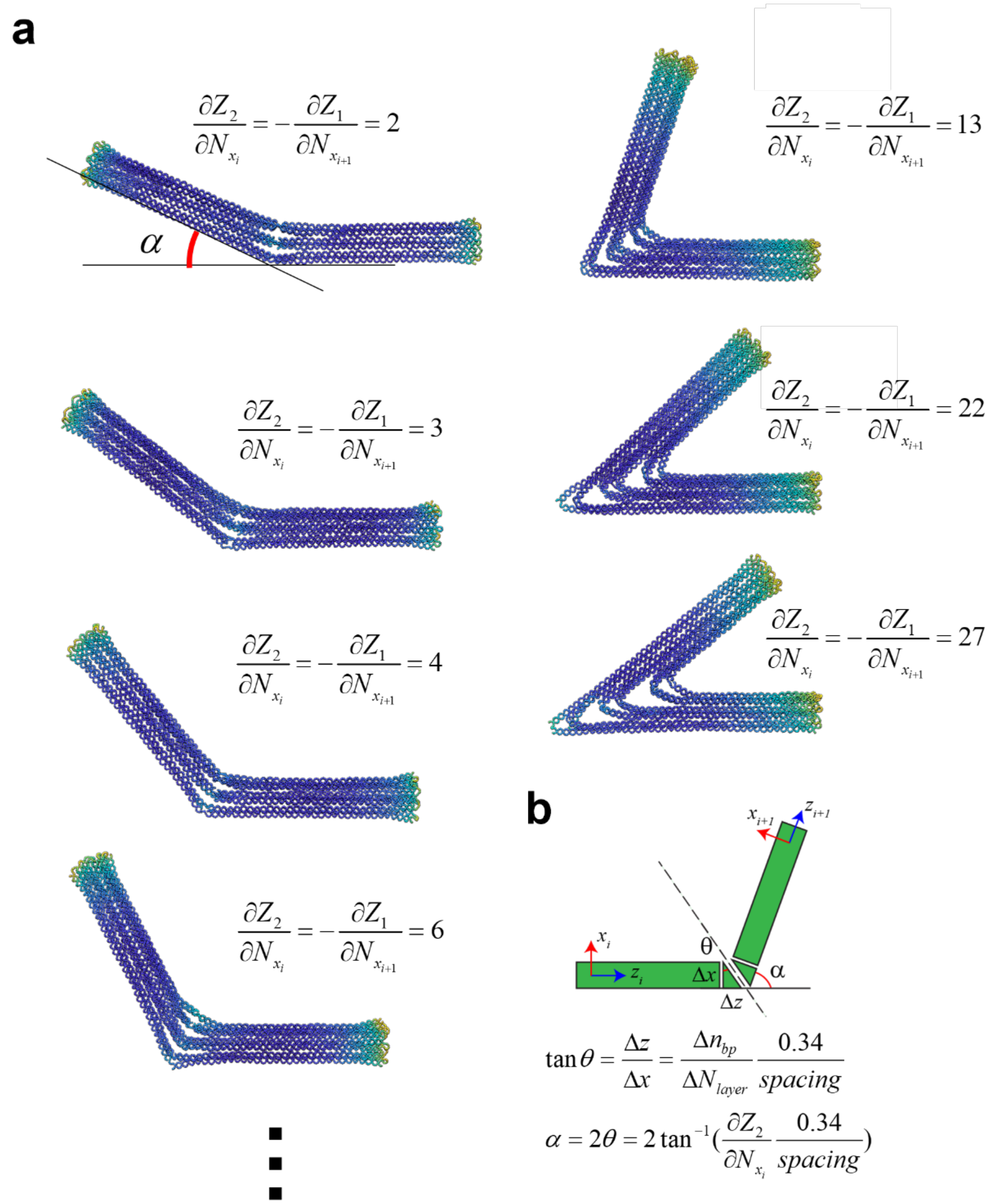


**Fig. S1.**
**Simulation of bending angles as function of edge gradients, using oxDNA.**

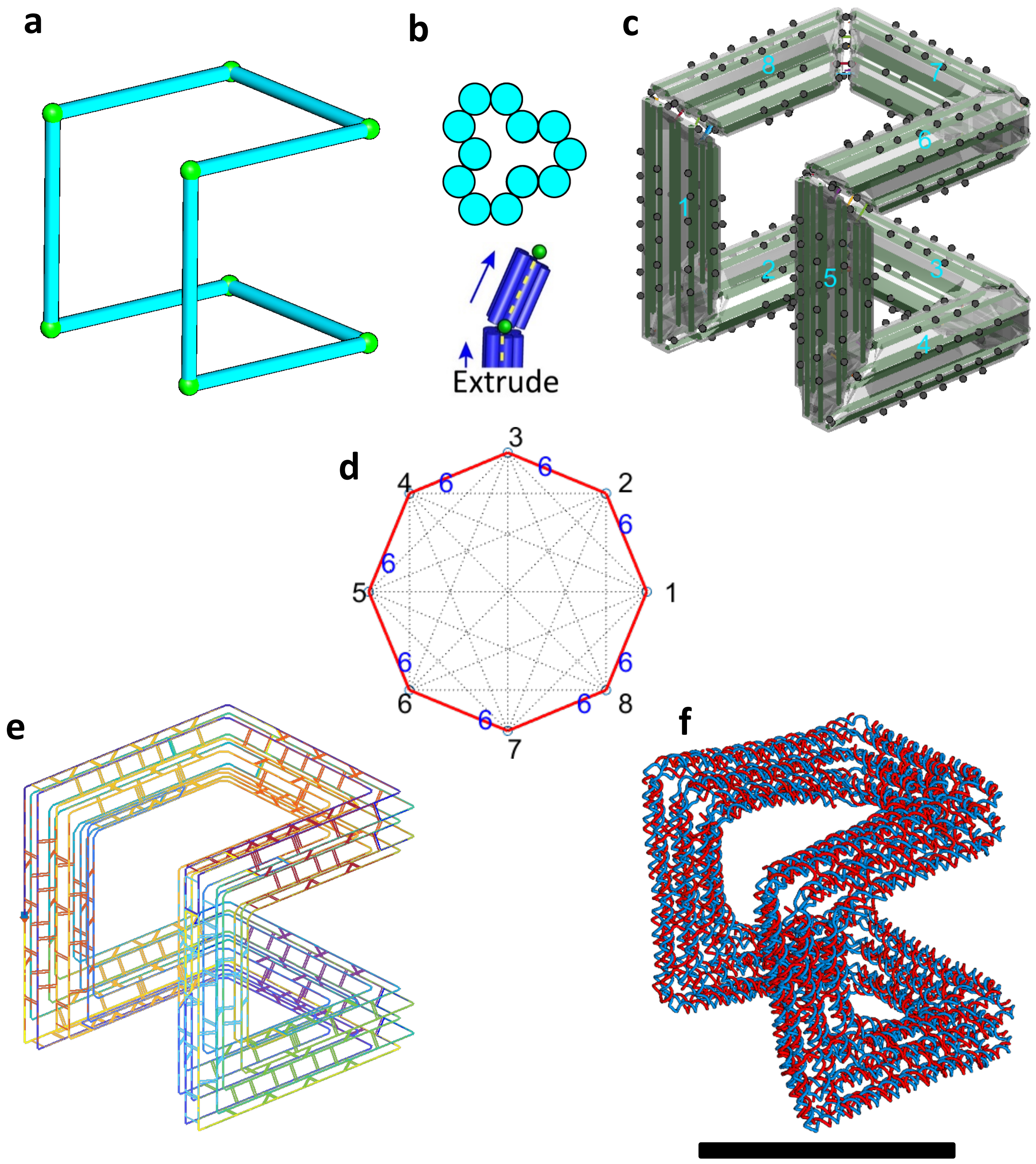


**Fig. S2.**

**Design details of the Hilbert structure.** (a) Line sketch of target structure. (b) Cross-section of each line. (c) Assembly model consisting of 8 bundles. (d) Connectivity matrix for assembly and routing algorithms. (e) Final scaffold and staple routings in the 3D representation. (f) oxDNA simulation result for the mean configuration. Scale bar = 50 nm.

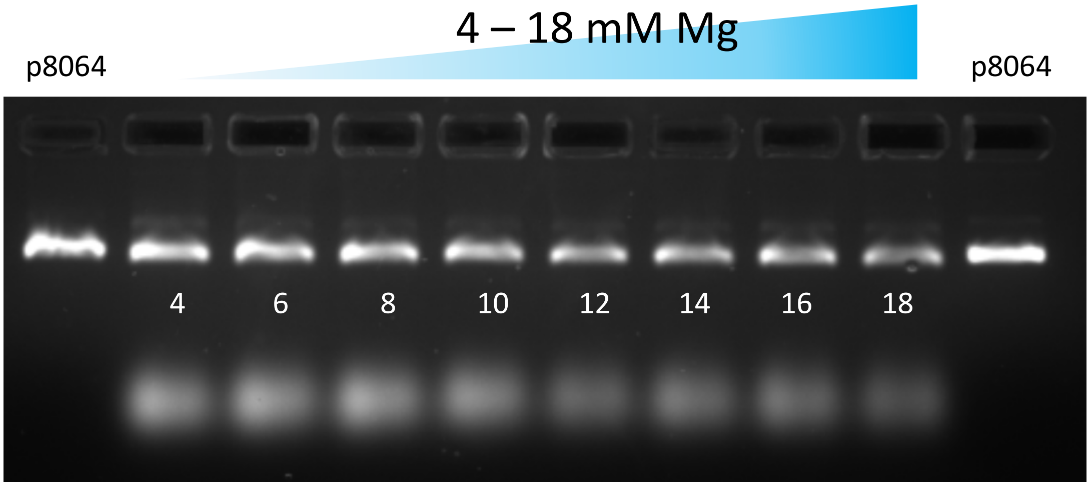


**Fig. S3.**

**Magnesium screening during the folding of the Hilbert structure.** Samples were run in a 1.5% agarose gel, using 45 mM Tris, 45 mM boric acid, 1 mM EDTA and 11 mM $MgCl_2$ as buffer. The gel was run at 90 V for 90 minutes and prestained with Ethidium bromide.

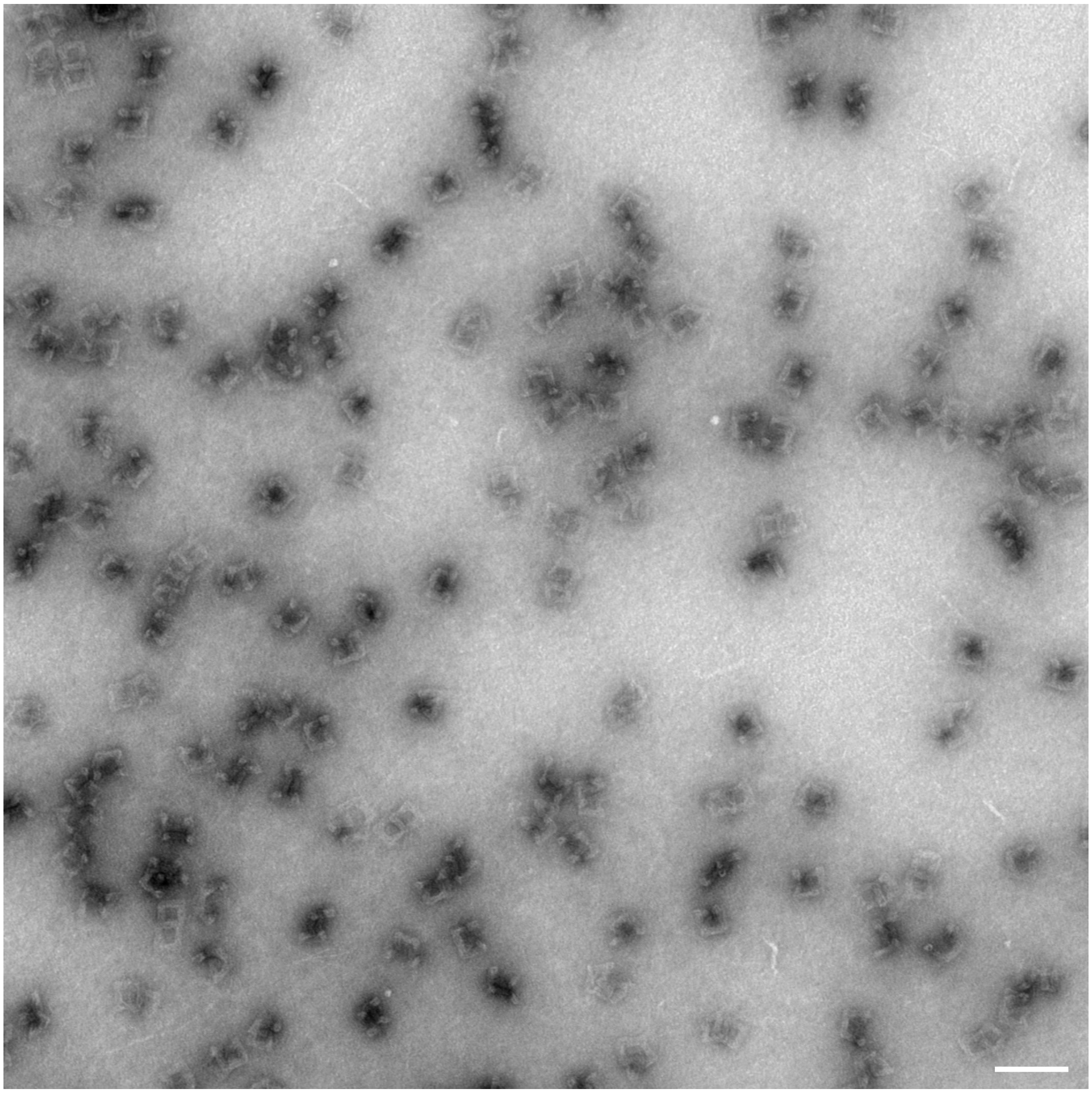

**Fig. S4.**

**TEM micrograph the Hilbert structure.** The sample was purified via the Freeze `N Squeeze kit and stained with 1% Uranyl acetate. Scale bar = 100 nm.

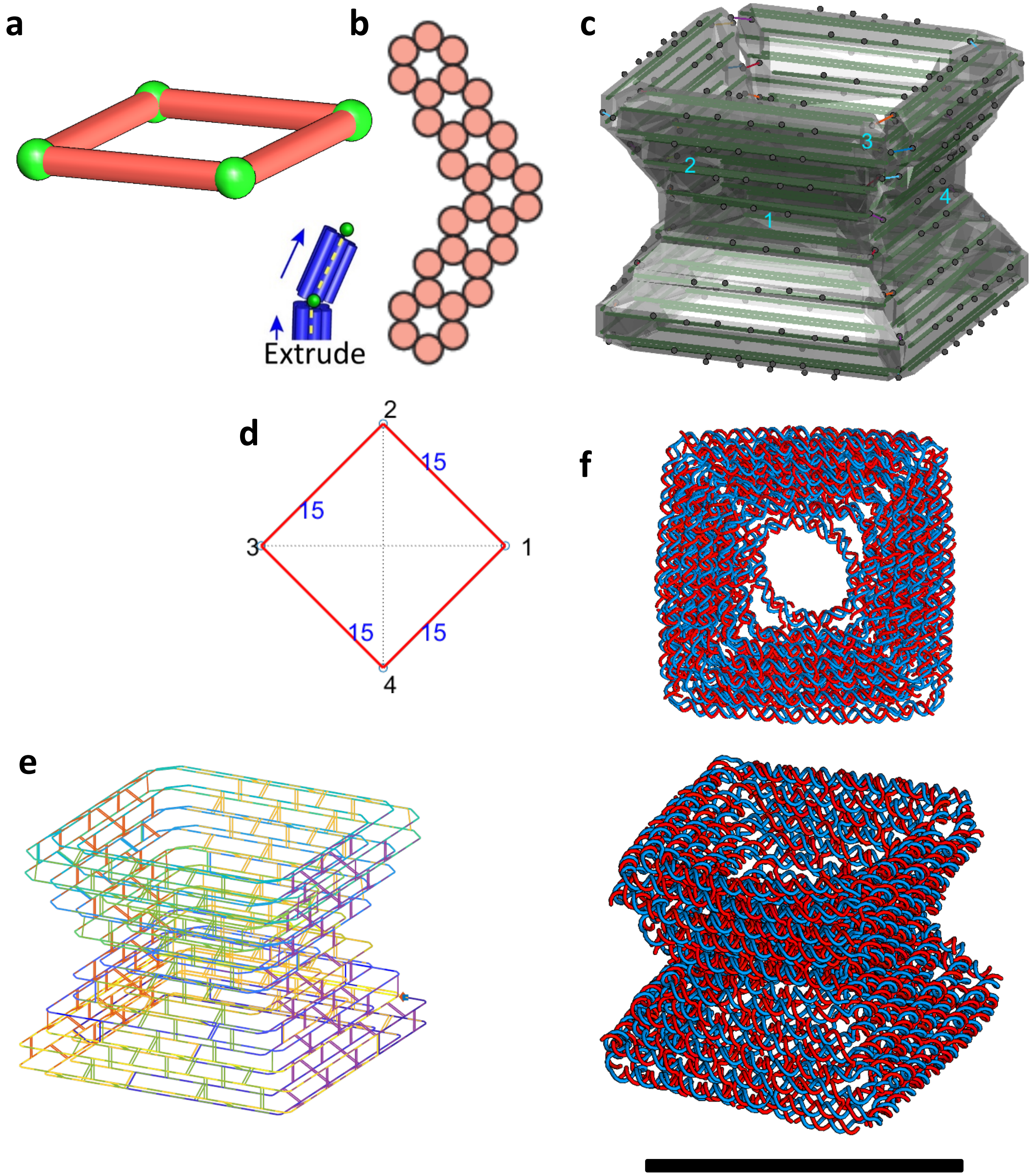


**Fig. S5.**

**Design details of the Nozzle structure.** (a) Line sketch of target structure. (b) Cross-section of each line. (c) Assembly model consisting of 4 bundles. (d) Connectivity matrix for assembly and routing algorithms. (e) Final scaffold and staple routings in the 3D representation. (f) oxDNA simulation result for the mean configuration. Scale bar = 50 nm.

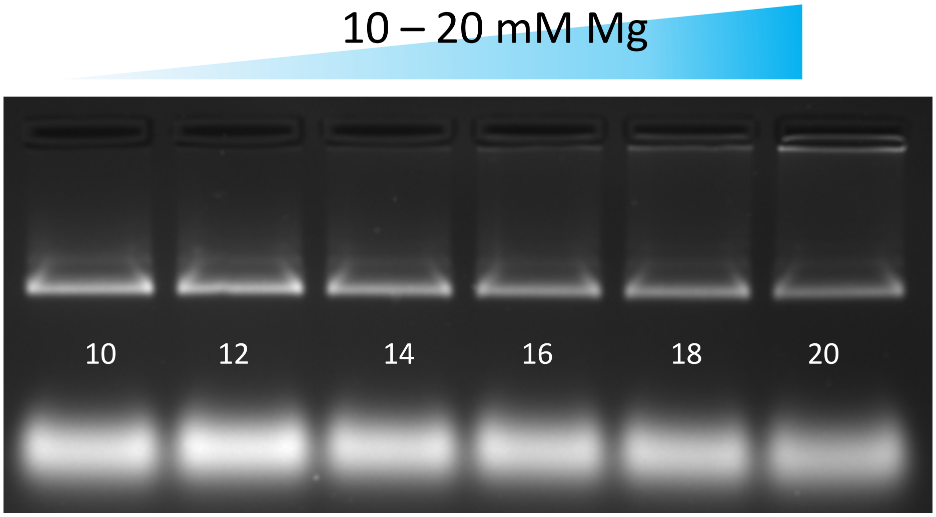


**Fig. S6.**

**Magnesium screening during the folding of the Nozzle structure.** Samples were run in a 1.5% agarose gel, using 45 mM Tris, 45 mM boric acid, 1 mM EDTA and 11 mM $MgCl_2$ as buffer. The gel was run at 90 V for 90 minutes and prestained with Ethidium bromide.

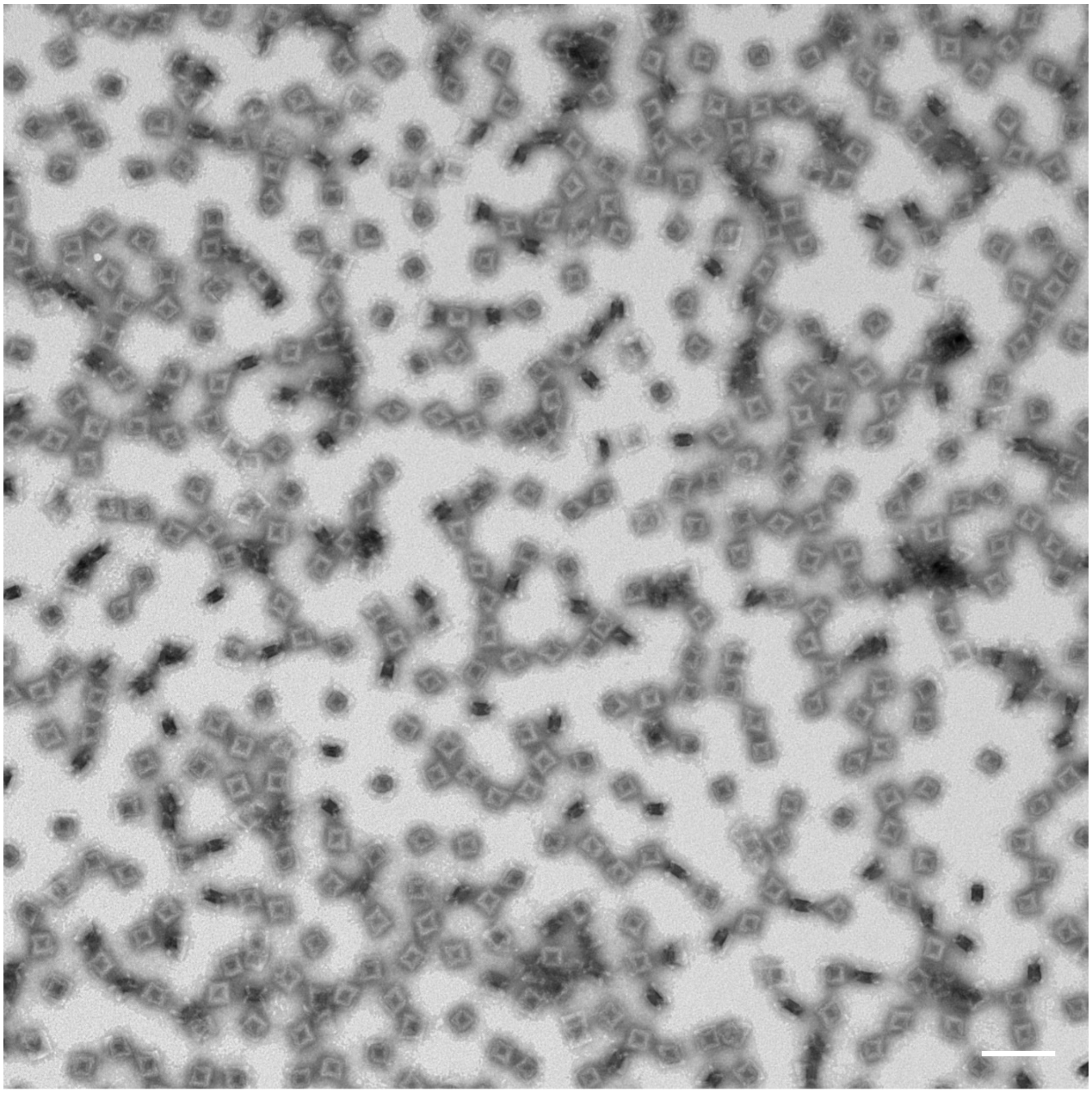

**Fig. S7.**
**TEM micrograph of the Nozzle structure (no overhangs).** The sample was purified via the Freeze `N Squeeze kit and stained with 1% Uranyl acetate. Scale bar = 100 nm.

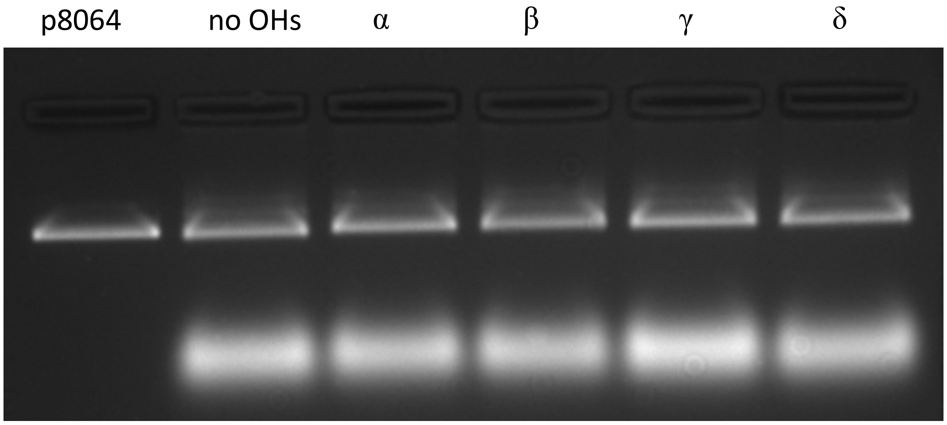


**Fig. S8.**

**Gel electrophoretic analysis of Nozzle variants with different overhangs**. Samples were run in a 1.5% agarose gel, using 45 mM Tris, 45 mM boric acid, 1 mM EDTA and 11 mM $MgCl_2$ as buffer. The gel was run at 90 V for 90 minutes and prestained with Ethidium bromide.

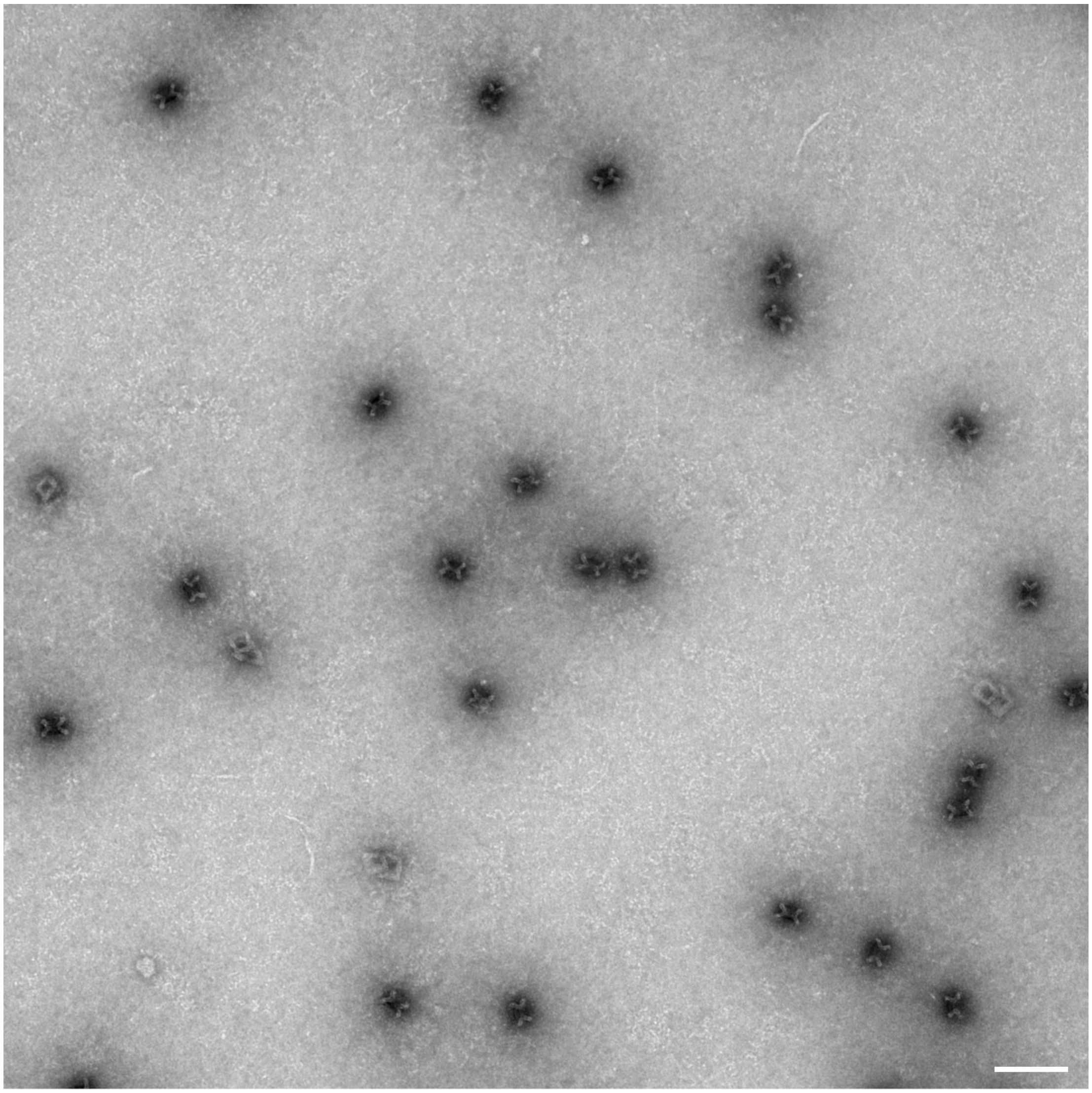

**Fig. S9.**
**TEM micrograph of the Nozzle structure (overhang species α).** The sample was purified via the Freeze `N Squeeze kit and stained with 1% Uranyl acetate. Scale bar = 100 nm.

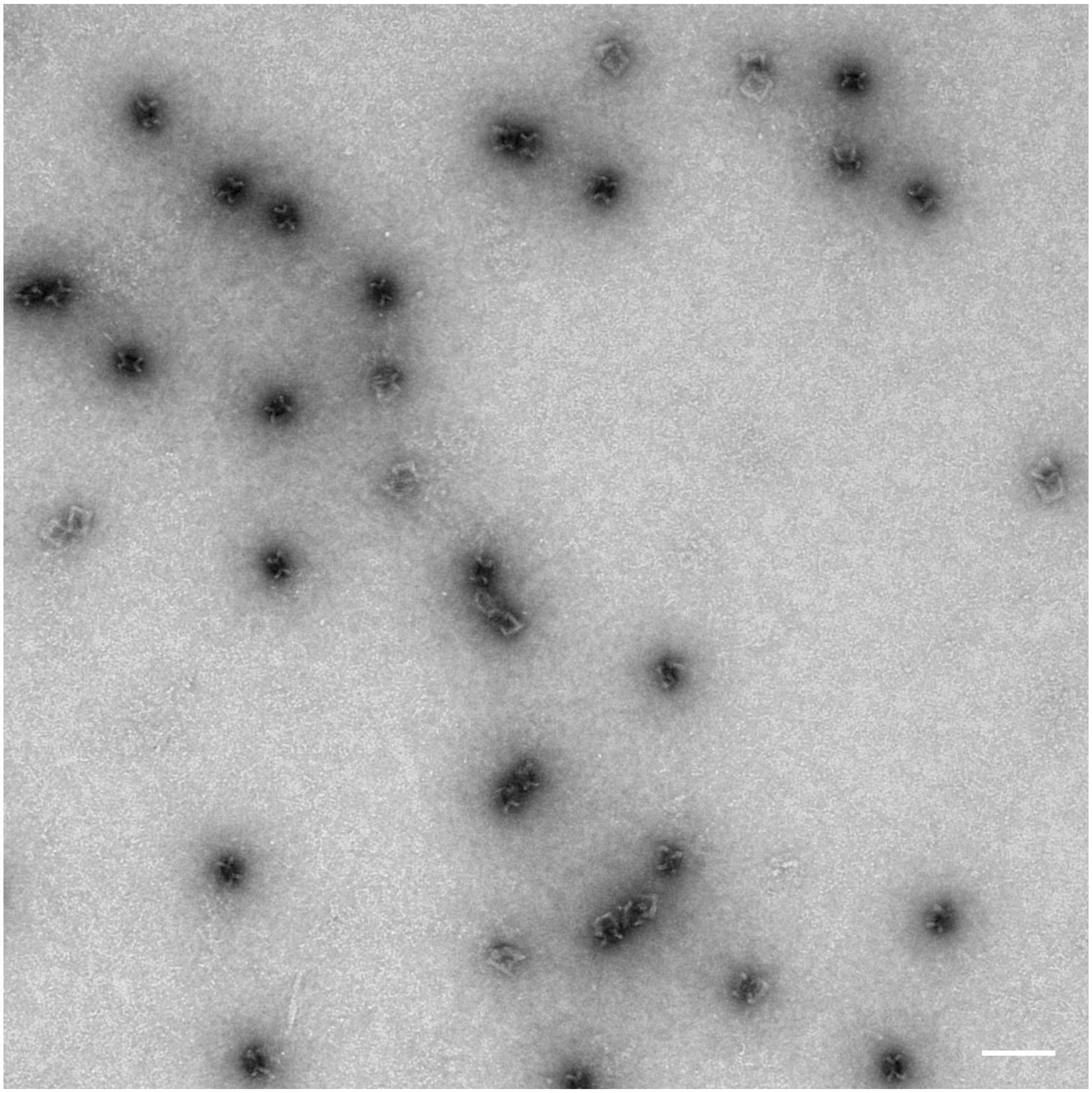

**Fig. S10.**
**TEM micrograph of the Nozzle structure (overhang species β).** The sample was purified via the Freeze `N Squeeze kit and stained with 1% Uranyl acetate. Scale bar = 100 nm.

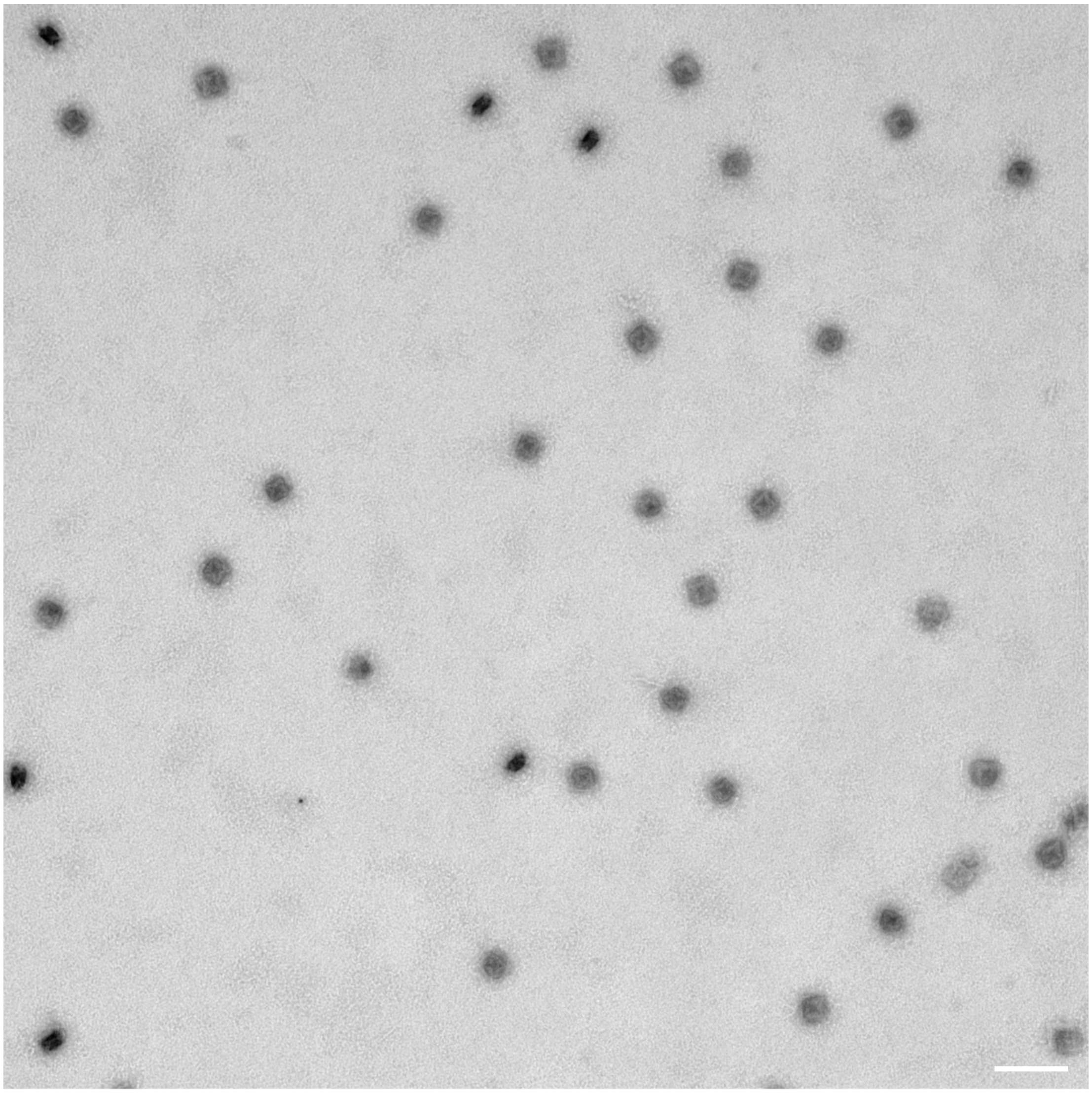

**Fig. S11.**

**TEM micrograph of the Nozzle structure (overhang species γ).** The sample was purified via the Freeze `N Squeeze kit and stained with 1% Uranyl acetate. Scale bar = 100 nm.

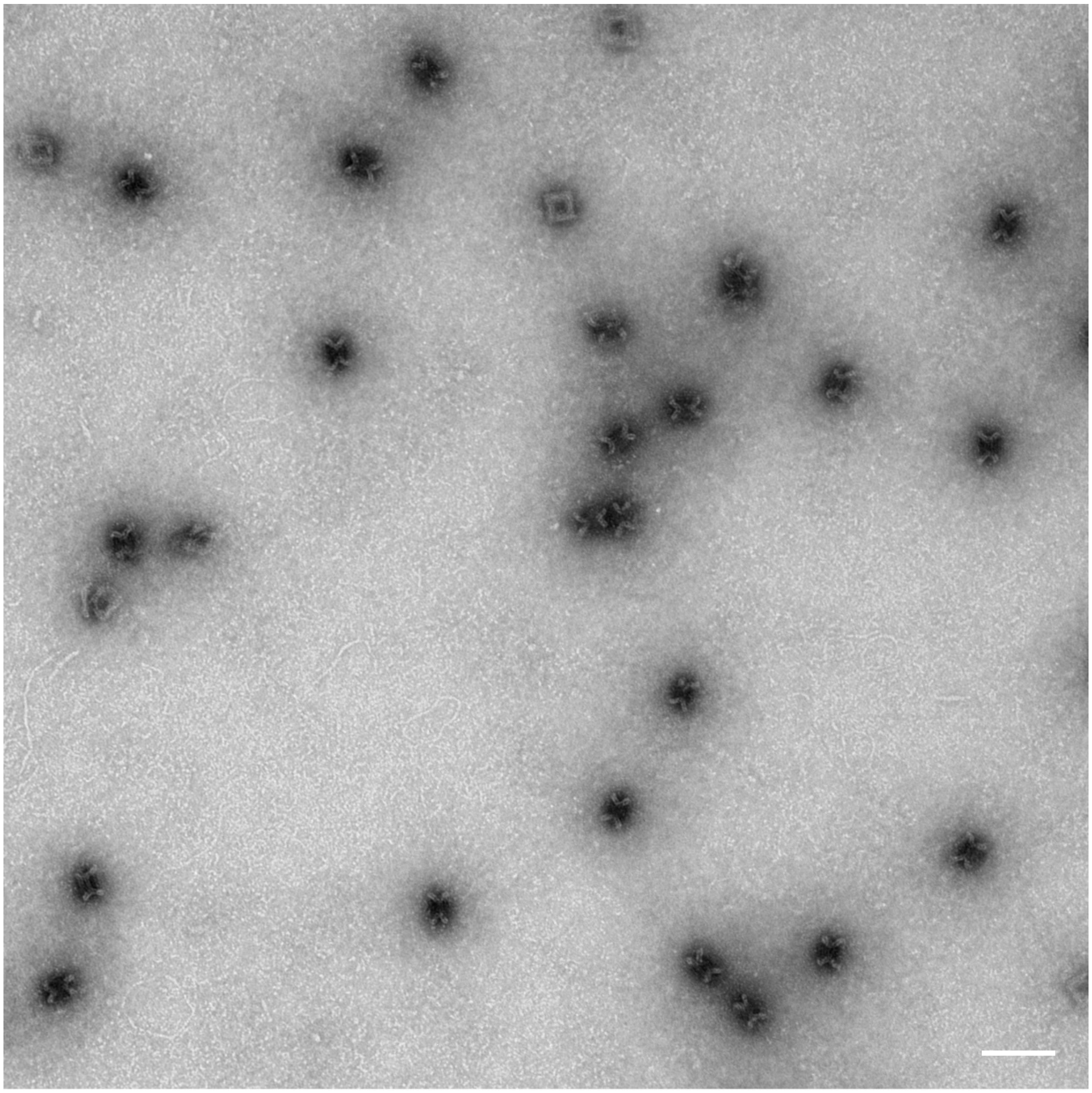

**Fig. S12.**
**TEM micrograph of the Nozzle structure (overhang species δ).** The sample was purified via the Freeze `N Squeeze kit and stained with 1% Uranyl acetate. Scale bar = 100 nm.

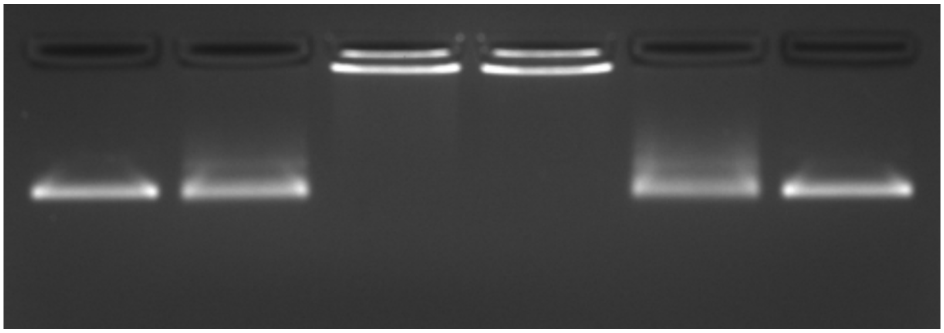


**Fig. S13.**

**Gel electrophoretic analysis of the multimerization of Nozzle structures with overhangs.** Species α and overhangs Species β. Multimerization was performed at different Magnesium concentrations for 20 h at 40°C. Samples were run in a 1.5% agarose gel, using 45 mM Tris, 45 mM boric acid, 1 mM EDTA and 11 mM $MgCl_2$ as buffer. The gel was run at 90 V for 90 minutes and prestained with Ethidium bromide.

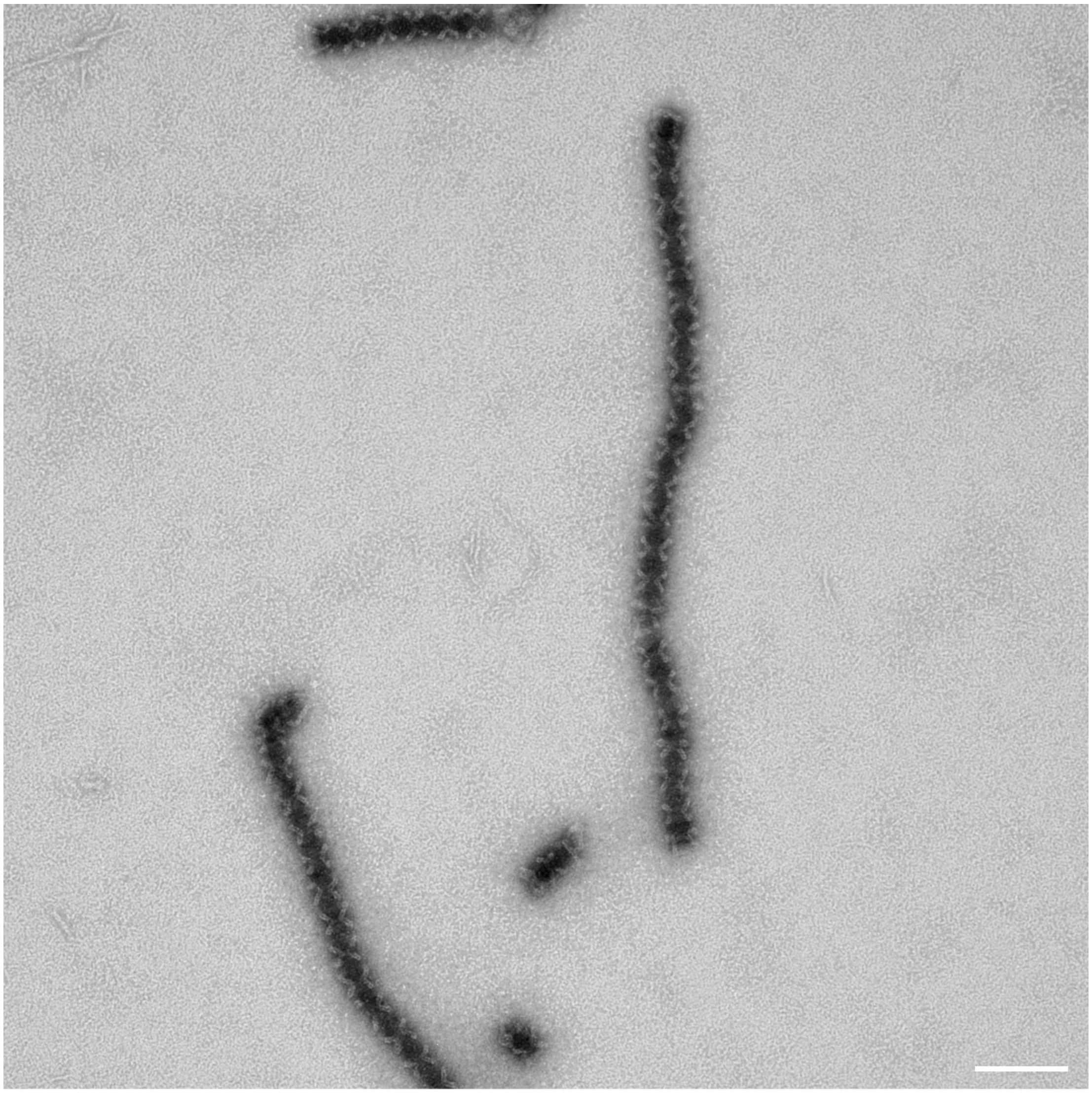

**Fig. S14.**

**TEM micrograph of multimeric Nozzle structure (overhang species α & β), incubated in the presence of 10 mM $MgCl_2$.** Individual species were purified via Freeze `N Squeeze prior to multimerization. Samples were stained with 1% Uranyl acetate. Scale bar = 100 nm.

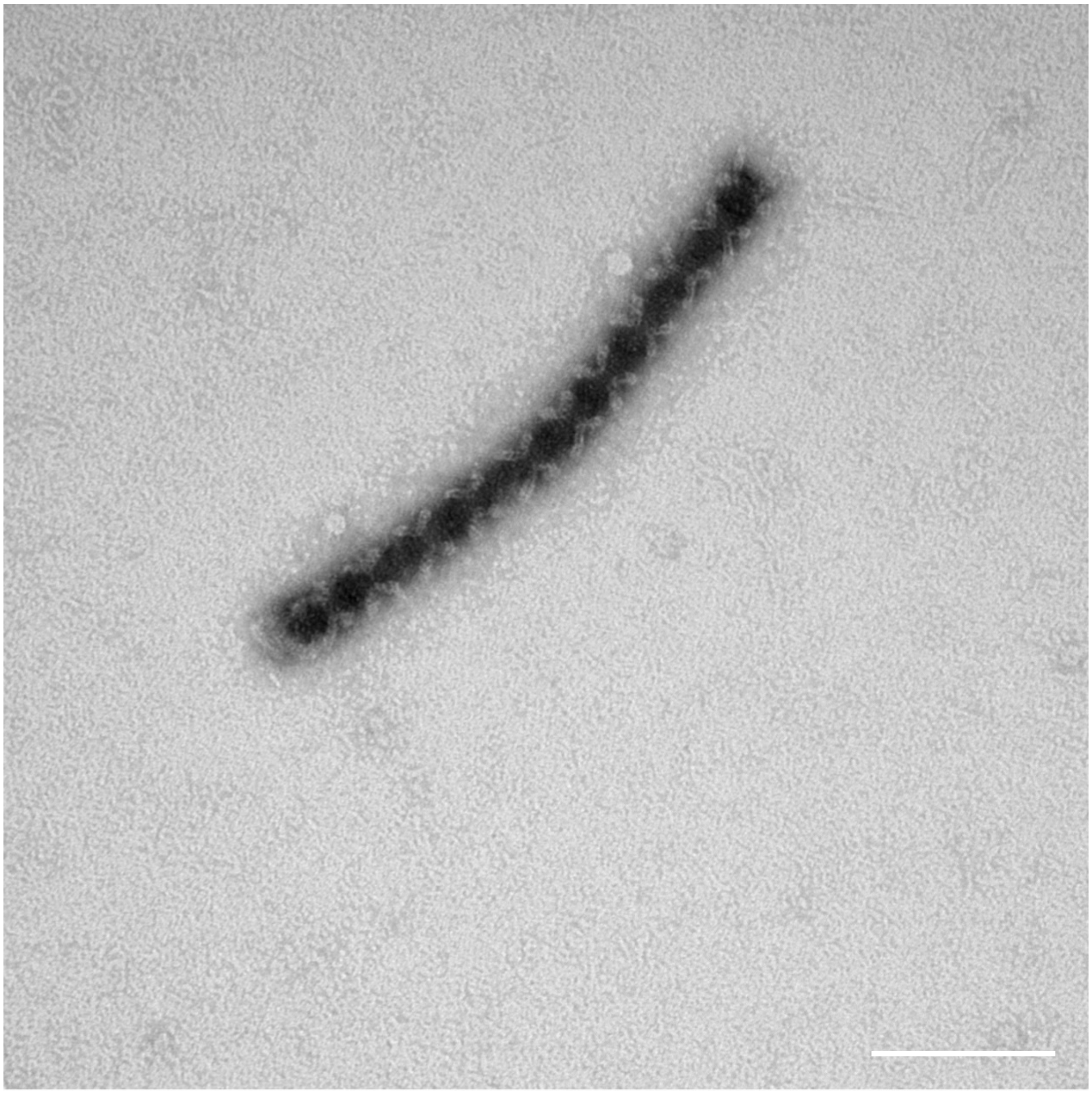

**Fig. S15.**

**TEM micrograph of multimeric Nozzle structure (overhang species α & β), incubated in the presence of 20 mM $MgCl_2$.** Individual species were purified via Freeze `N Squeeze prior to multimerization. Samples were stained with 1% Uranyl acetate. Scale bar = 100 nm.

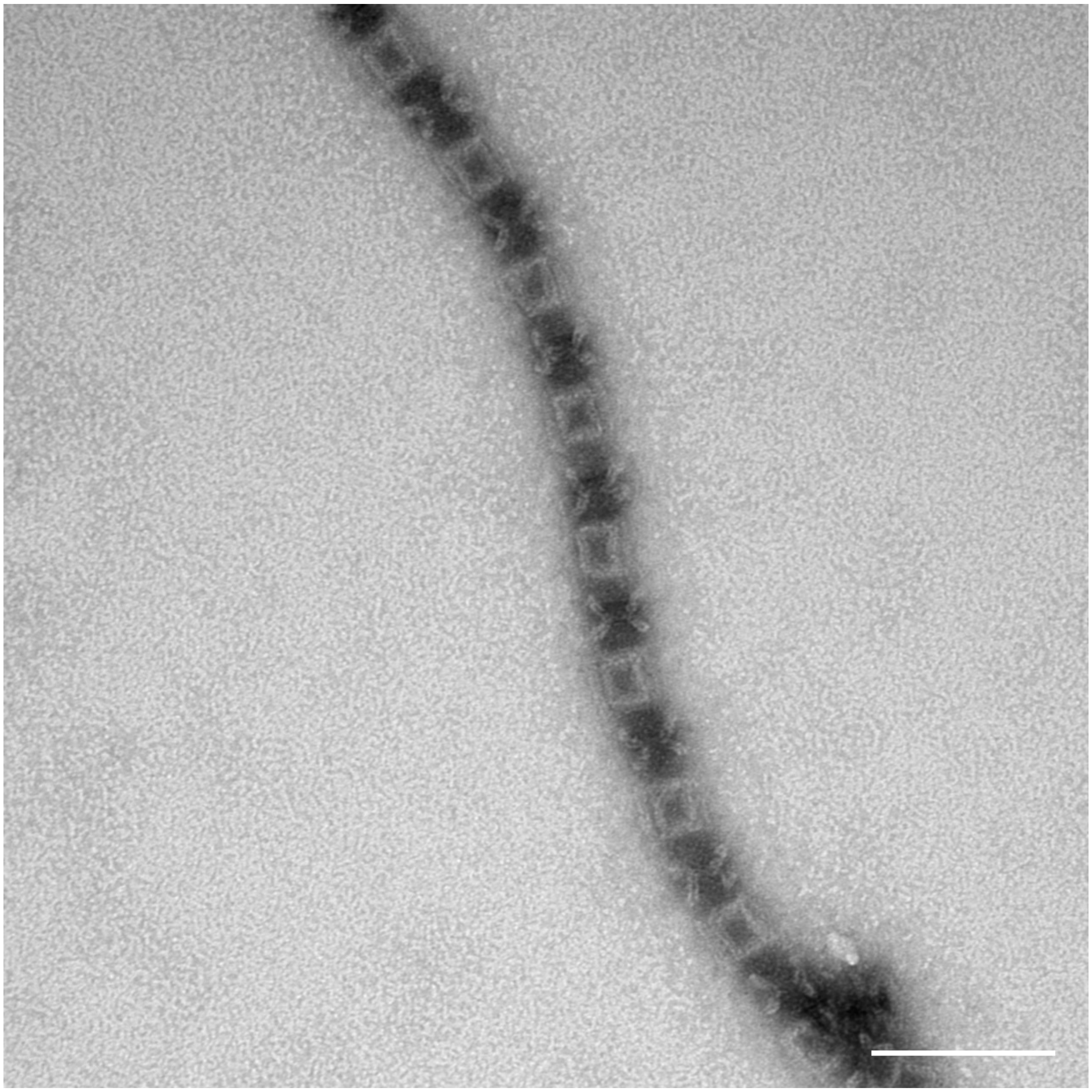

**Fig. S16.**

**TEM micrograph of multimeric Nozzle structure (overhang species γ & δ), incubated in the presence of 10 mM $MgCl_2$.** Individual species were purified via Freeze `N Squeeze prior to multimerization. Samples were stained with 1% Uranyl acetate. Scale bar = 100 nm.

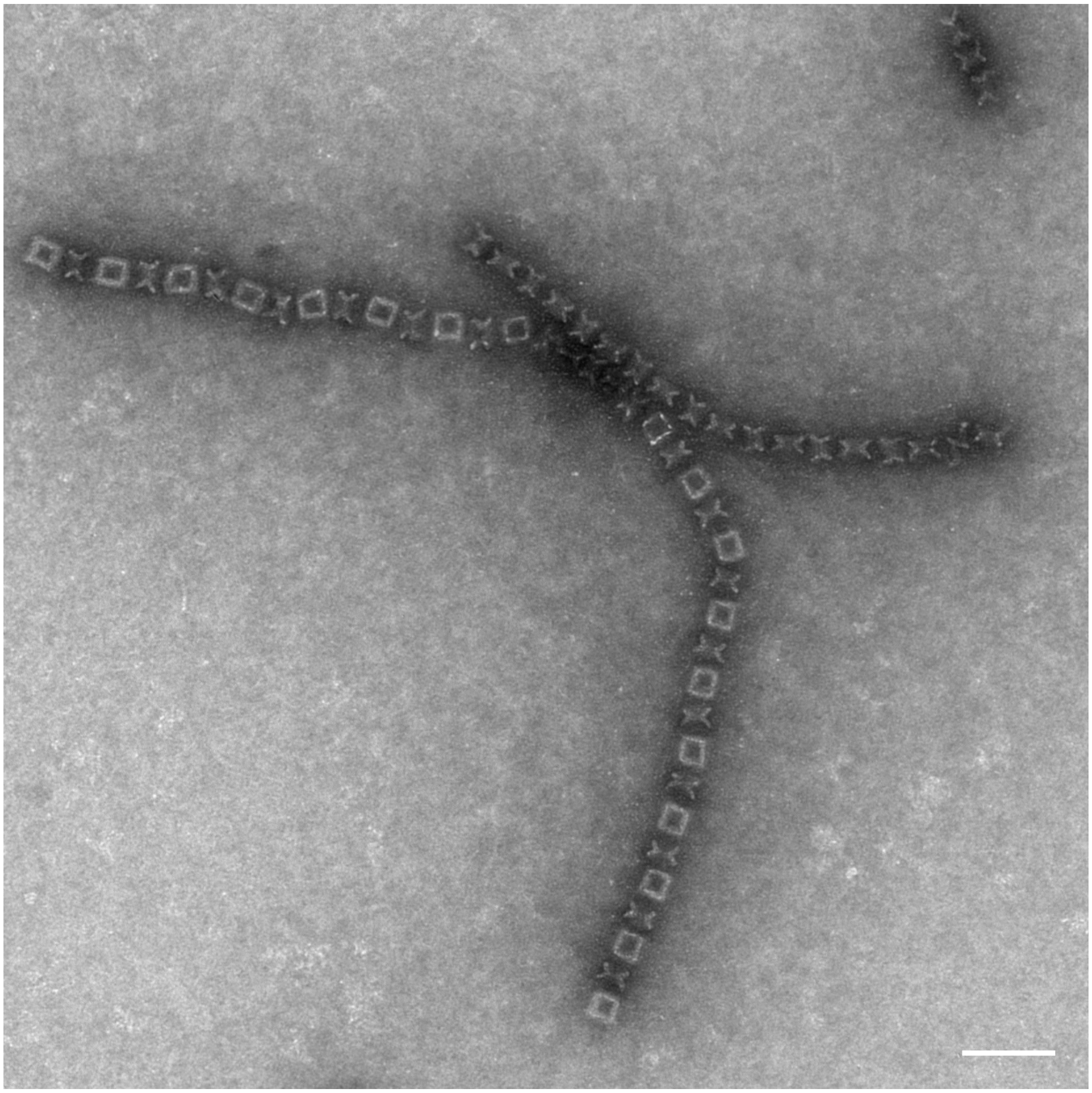

**Fig. S17.**

**TEM micrograph of multimeric Nozzle structure (overhang species γ & δ), incubated in the presence of 20 mM $MgCl_2$.** Individual species were purified via Freeze `N Squeeze prior to multimerization. Samples were stained with 1% Uranyl acetate. Scale bar = 100 nm.

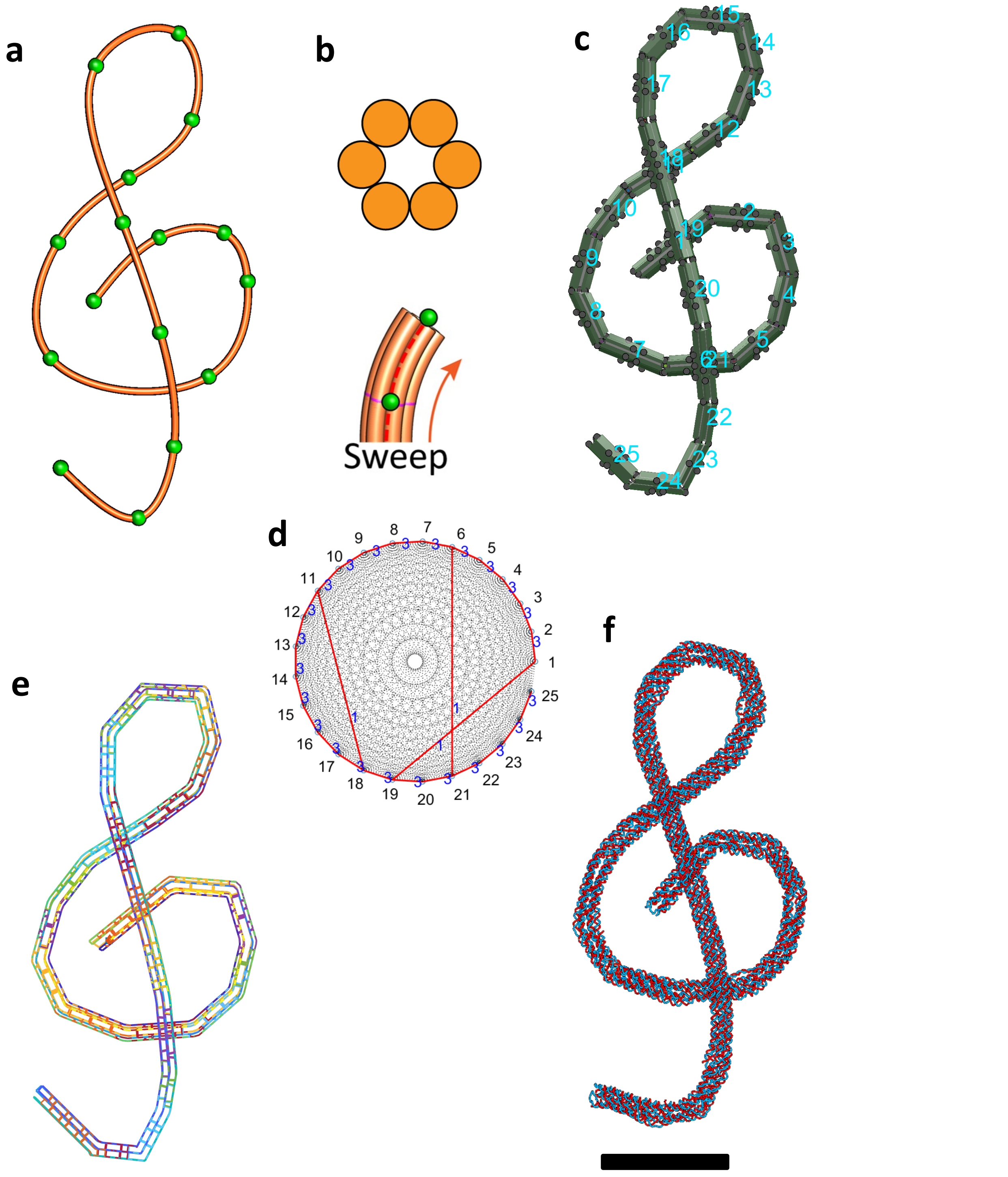


**Fig. S18.**

**Design details of the G-clef structure.** (a) Line sketch of target structure. (b) Cross-section of each line. (c) Assembly model consisting of 25 bundles. (d) Connectivity matrix for assembly and routing algorithms. (e) Final scaffold and staple routings in the 3D representation. (f) oxDNA simulation result for the mean configuration. Scale bar = 50 nm.

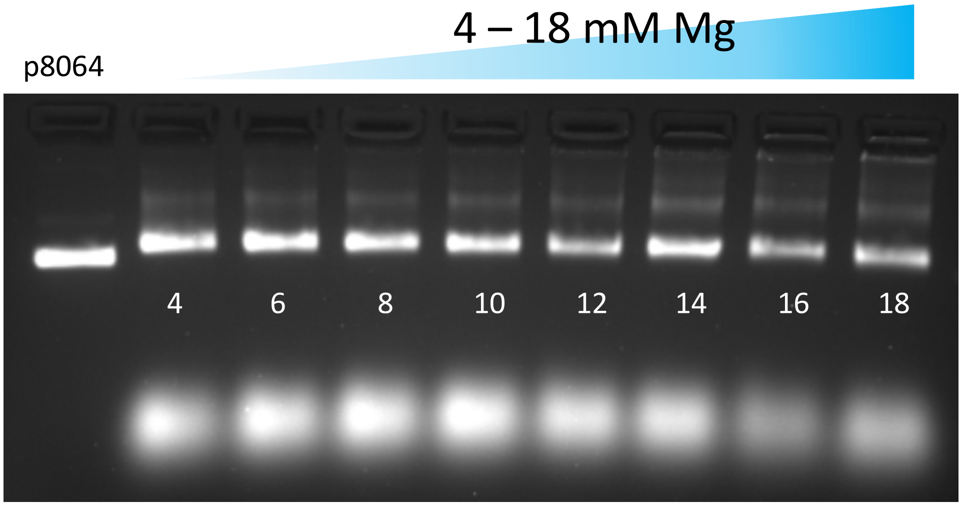


**Fig. S19.**

**Magnesium screening during the folding of the G-clef structure.** Samples were run in a 1.5% agarose gel, using 45 mM Tris, 45 mM boric acid, 1 mM EDTA and 11 mM $MgCl_2$ as buffer. The gel was run at 90 V for 90 minutes and prestained with Ethidium bromide.

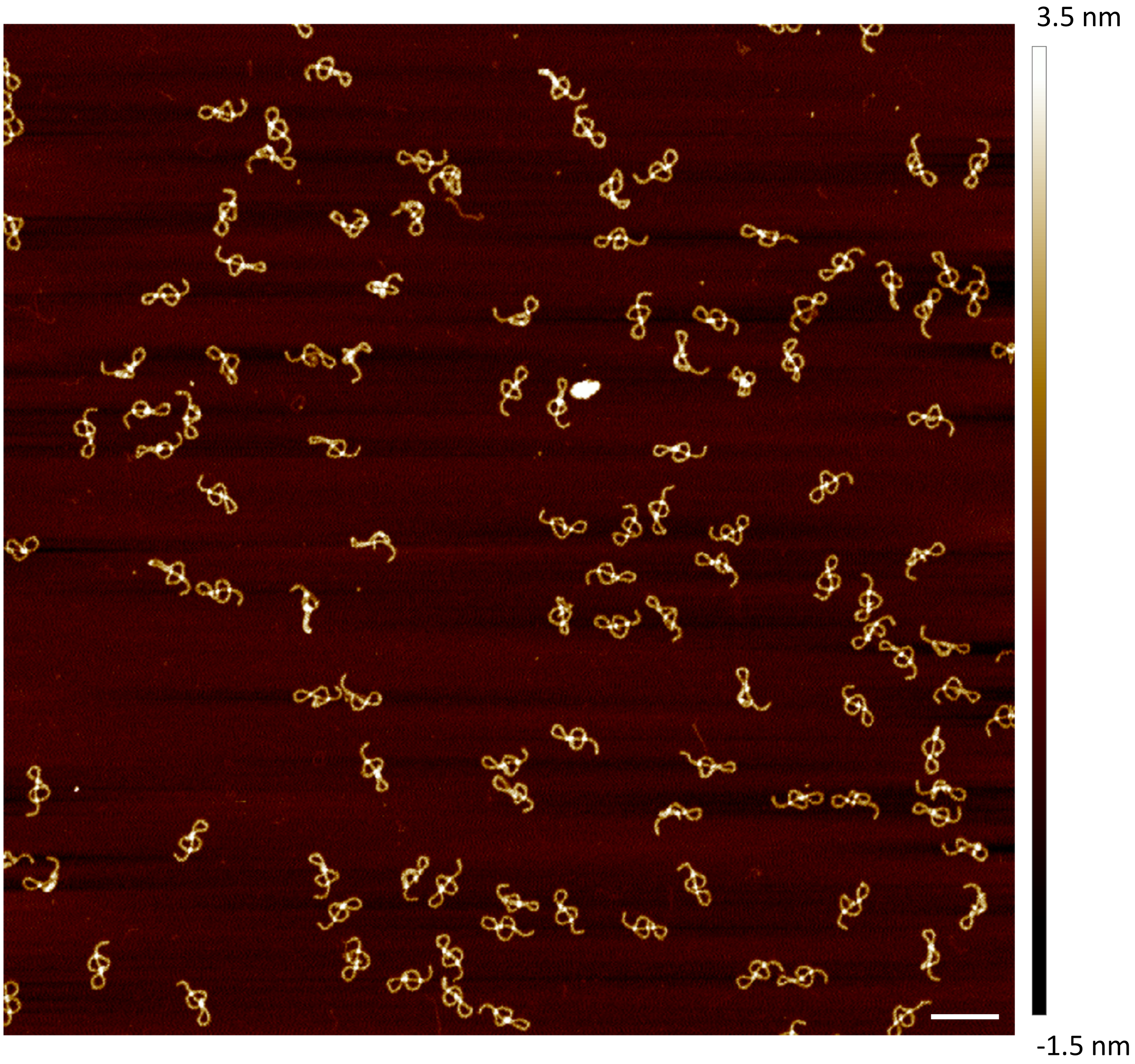


**Fig. S20.**

**AFM image of the G-clef structure.** Freeze ‘N Squeeze purified samples were incubated on freshly cleaved mica for 3 minutes, washed briefly with $H_2O$ and subsequently dried with a gentle flow of air. Samples were measured using ScanAsyst Air tips. Scale bar = 200 nm.

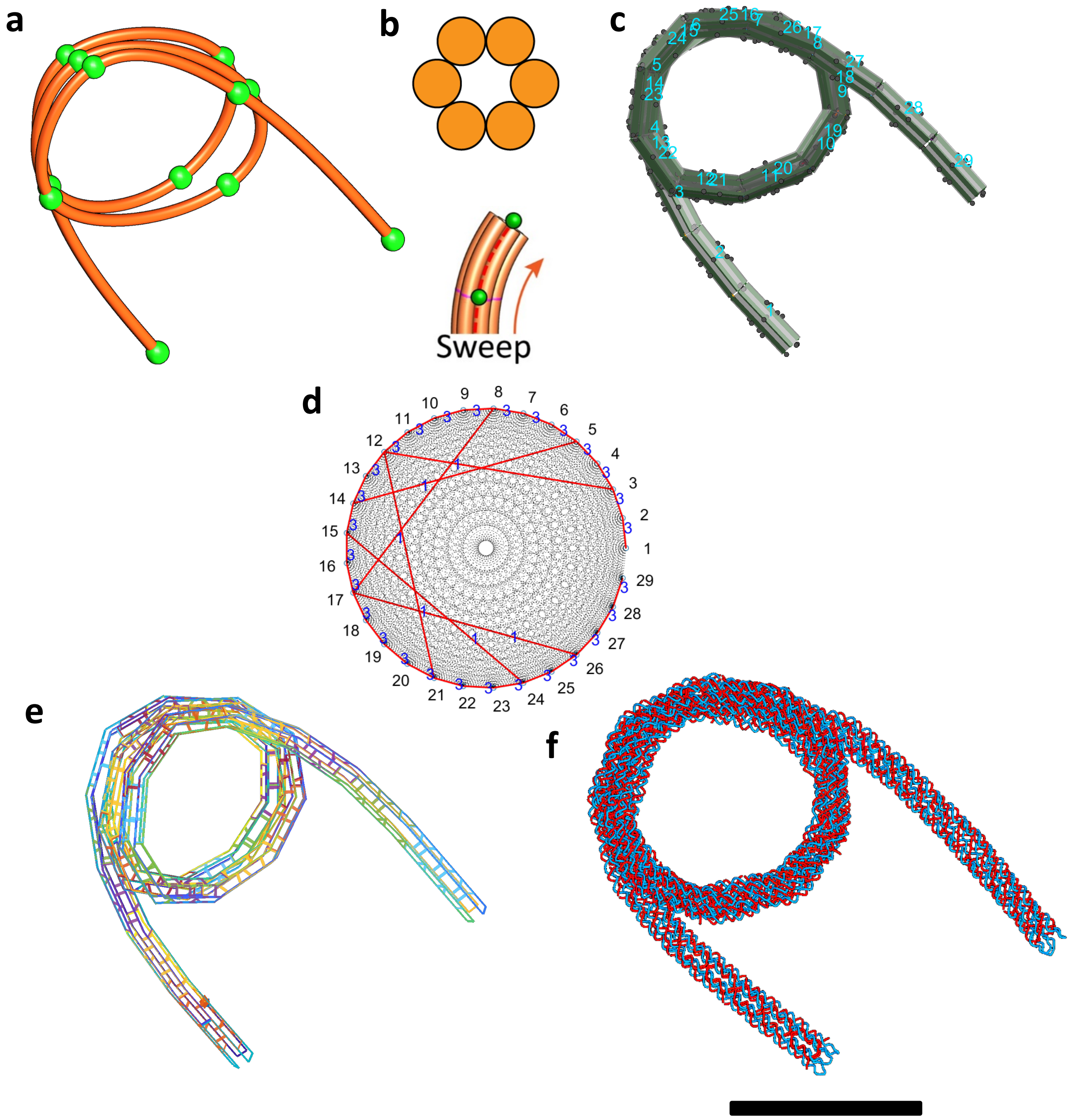


**Fig. S21.**

**Design details of the nucleosome-like spring structure.** (a) Line sketch of target structure. (b) Cross-section of each line. (c) Assembly model consisting of 29 bundles. (d) Connectivity matrix for assembly and routing algorithms. (e) Final scaffold and staple routings in the 3D representation. (f) oxDNA simulation result for the mean configuration. Scale bar = 50 nm.

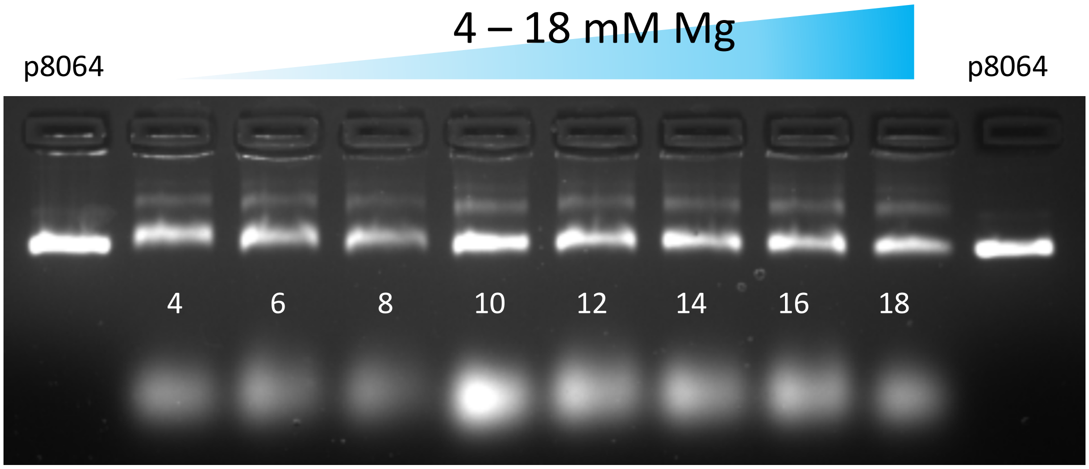


**Fig. S22.**

**Magnesium screening during the folding of the nucleosome-like spring structure.** Samples were run in a 1.5% agarose gel, using 45 mM Tris, 45 mM boric acid, 1 mM EDTA and 11 mM $MgCl_2$ as buffer. The gel was run at 90 V for 90 minutes and prestained with Ethidium bromide.

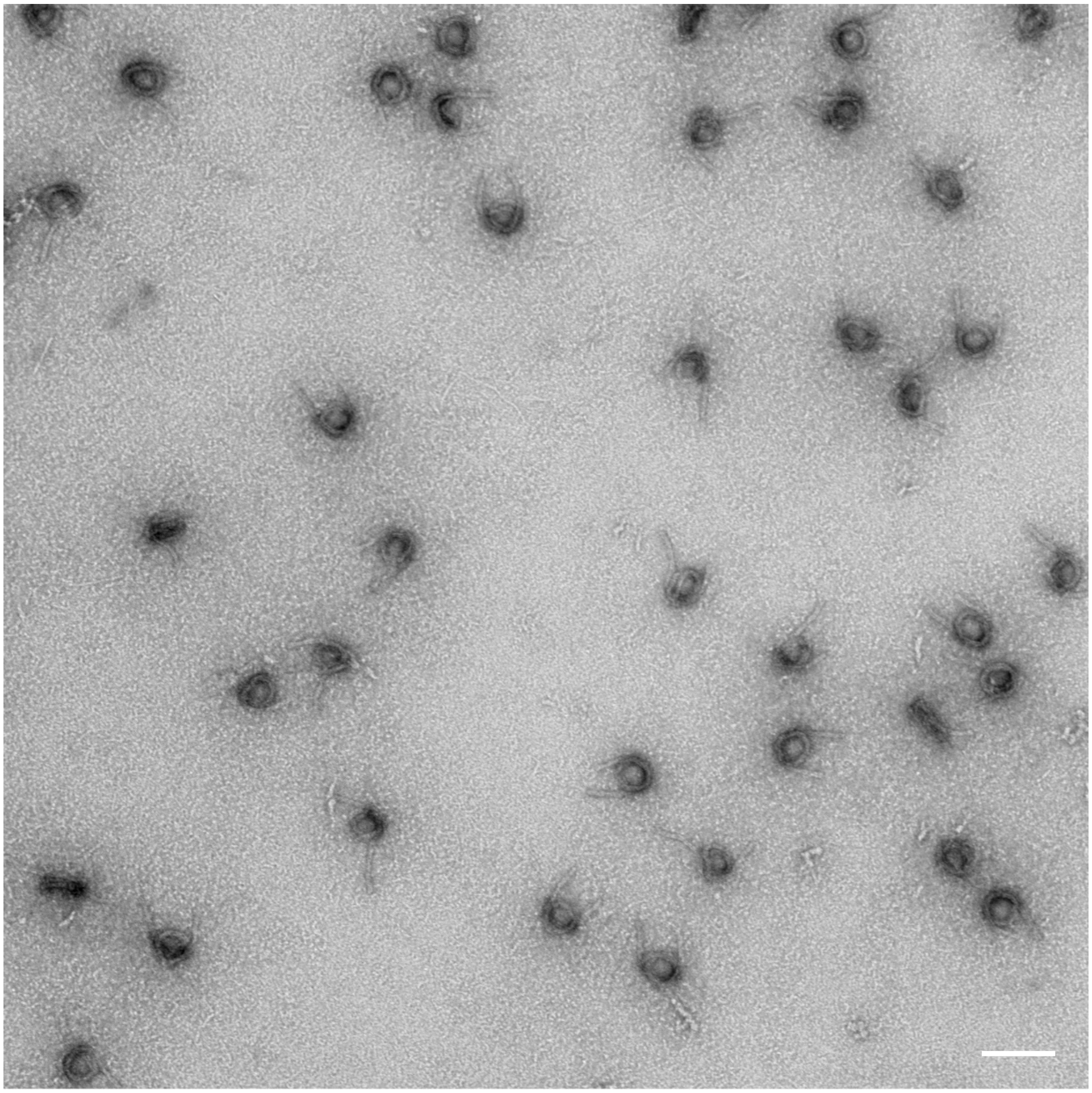

**Fig. S23.**
**TEM micrograph of the nucleosome-like spring structure.** The sample was purified via the Freeze `N Squeeze kit and stained with 1% Uranyl acetate. Scale bar = 100 nm.

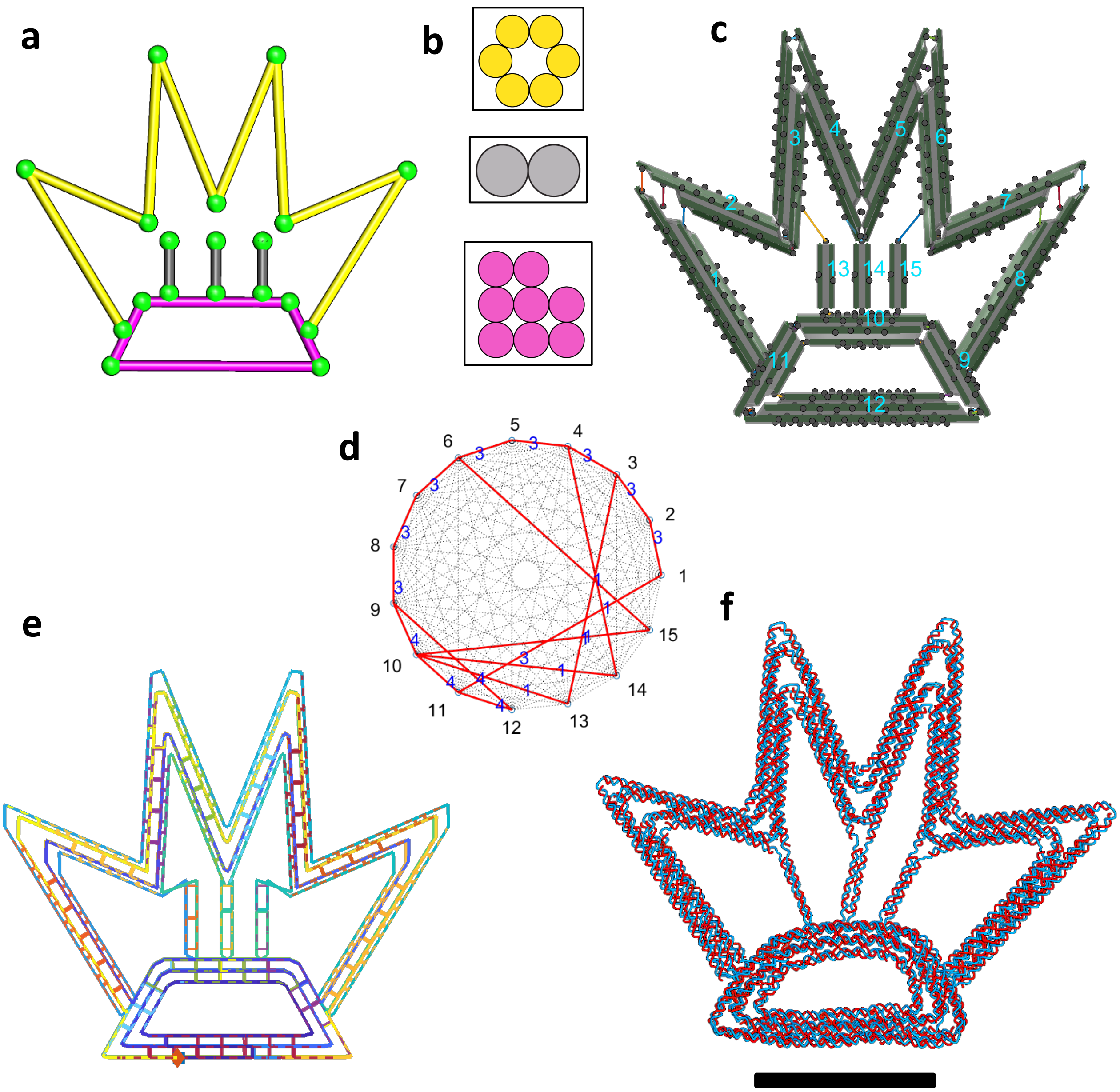


**Fig. S24.**

**Design details of the crown structure.** (a) Line sketch of target structure. (b) Cross-section of each line. The bundles in the top portion (yellow spline), in three trusses, and in the bottom base were assigned with 6HB, 2HB, and 8HB, respectively, (c) Assembly model consisting of 15 bundles. (d) Connectivity matrix for assembly and routing algorithms. (e) Final scaffold and staple routings in the 3D representation. (f) oxDNA simulation result for the mean configuration. Scale bar = 50 nm.

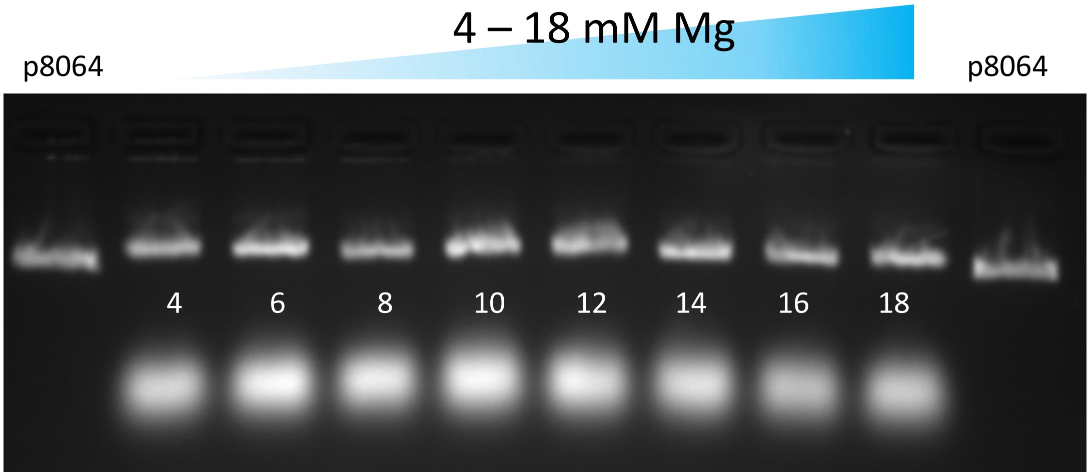


**Fig. S25.**

**Magnesium screening during the folding of the crown structure.** Samples were run in a 1.5% agarose gel, using 45 mM Tris, 45 mM boric acid, 1 mM EDTA and 11 mM $MgCl_2$ as buffer. The gel was run at 90 V for 90 minutes and prestained with Ethidium bromide.

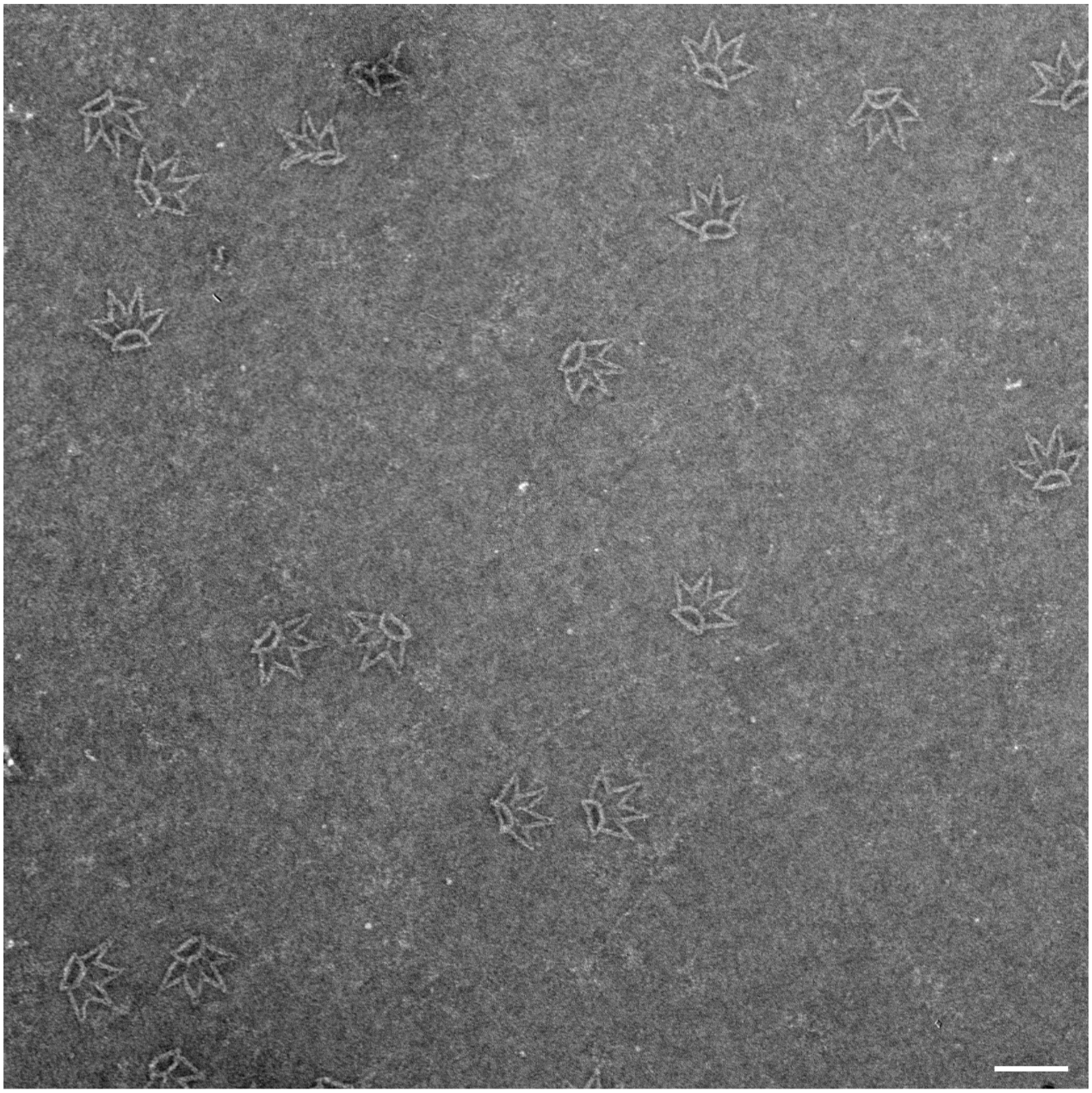

**Fig. S26.**
**TEM micrograph of crown structure.** The sample was purified via the Freeze `N Squeeze kit and stained with 1% Uranyl acetate. Scale bar = 100 nm.

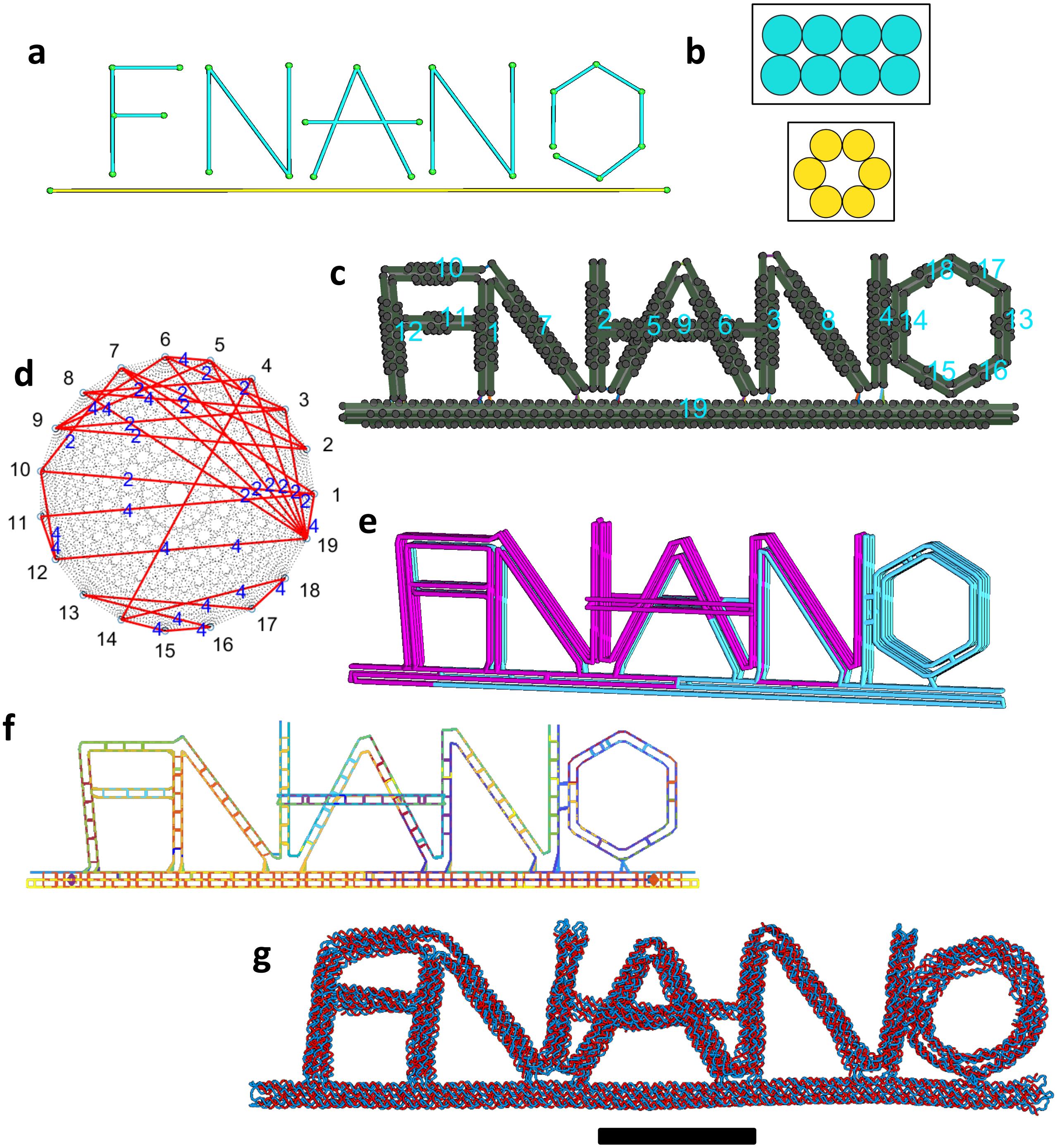


**Fig. S27.**

**Design details of the FNANO-script structure.** (a) Line sketch of target structure. (b) Cross-section of each line. The bundles in the top portion (cyan spline) and in the underline were assigned with 8HB and 6HB, respectively, (c) Assembly model consisting of 19 bundles. (d) Connectivity matrix for assembly and routing algorithms. (e) Two-scaffold routings (PCS3_L_7560 [ref] and p8064) using the multi-scaffold algorithm. (f) Final scaffold and staple routings in the 3D representation. (g) oxDNA simulation result for the mean configuration. Scale bar = 50 nm.

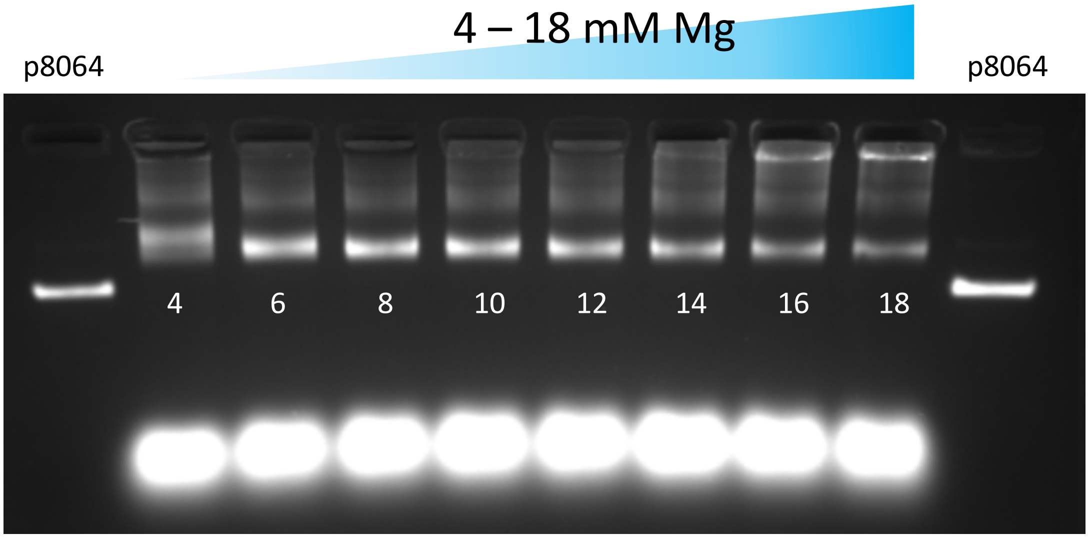


**Fig. S28.**

**Magnesium screening during the folding of the FNANO-script structure.** Samples were run in a 1.5% agarose gel, using 45 mM Tris, 45 mM boric acid, 1 mM EDTA and 11 mM $MgCl_2$ as buffer. The gel was run at 90 V for 90 minutes and prestained with Ethidium bromide.

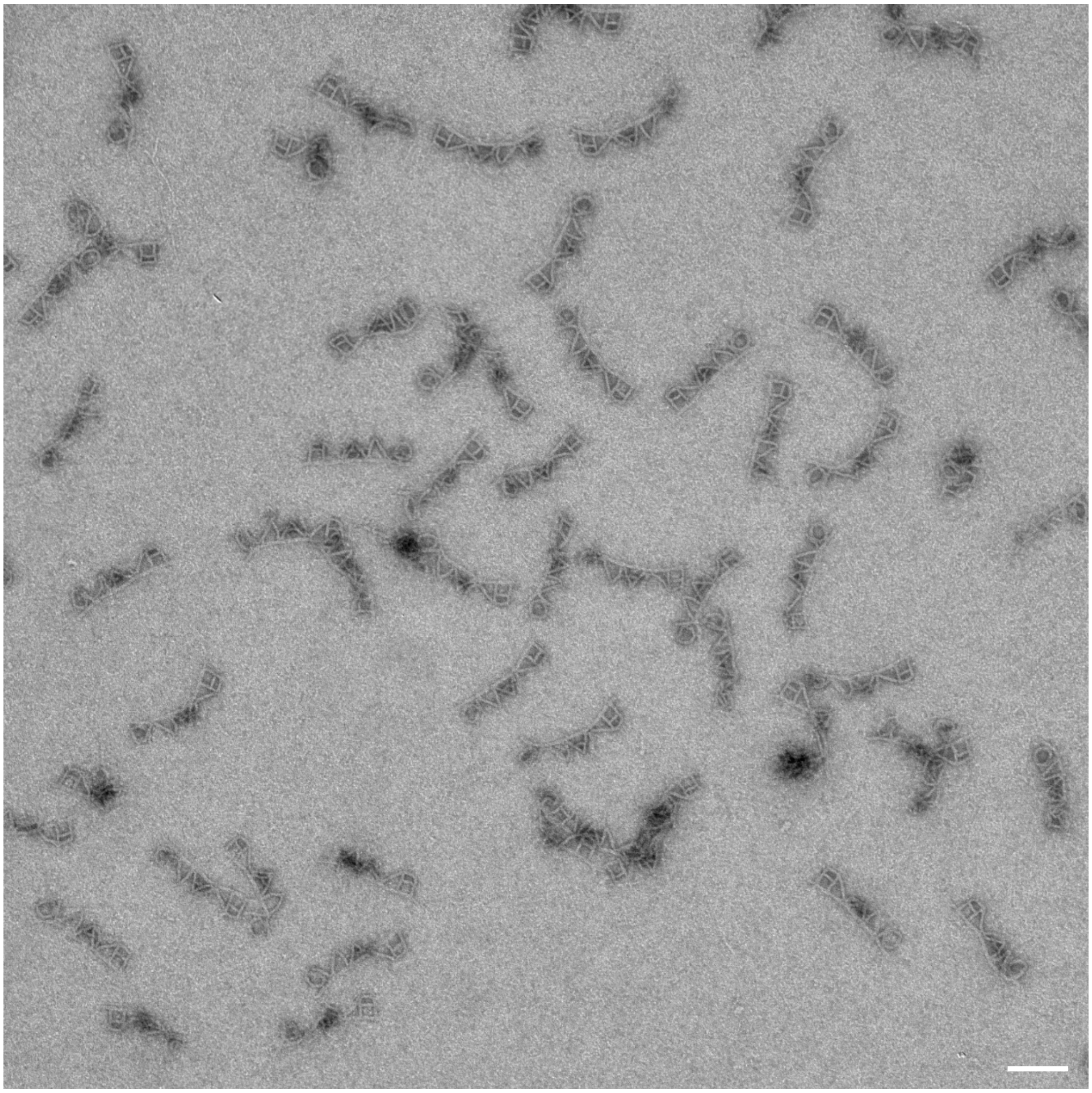

**Fig. S29.**
**TEM micrograph of the FNANO-script structure.** The sample was purified via the Freeze `N Squeeze kit and stained with 1% Uranyl acetate. Scale bar = 100 nm.

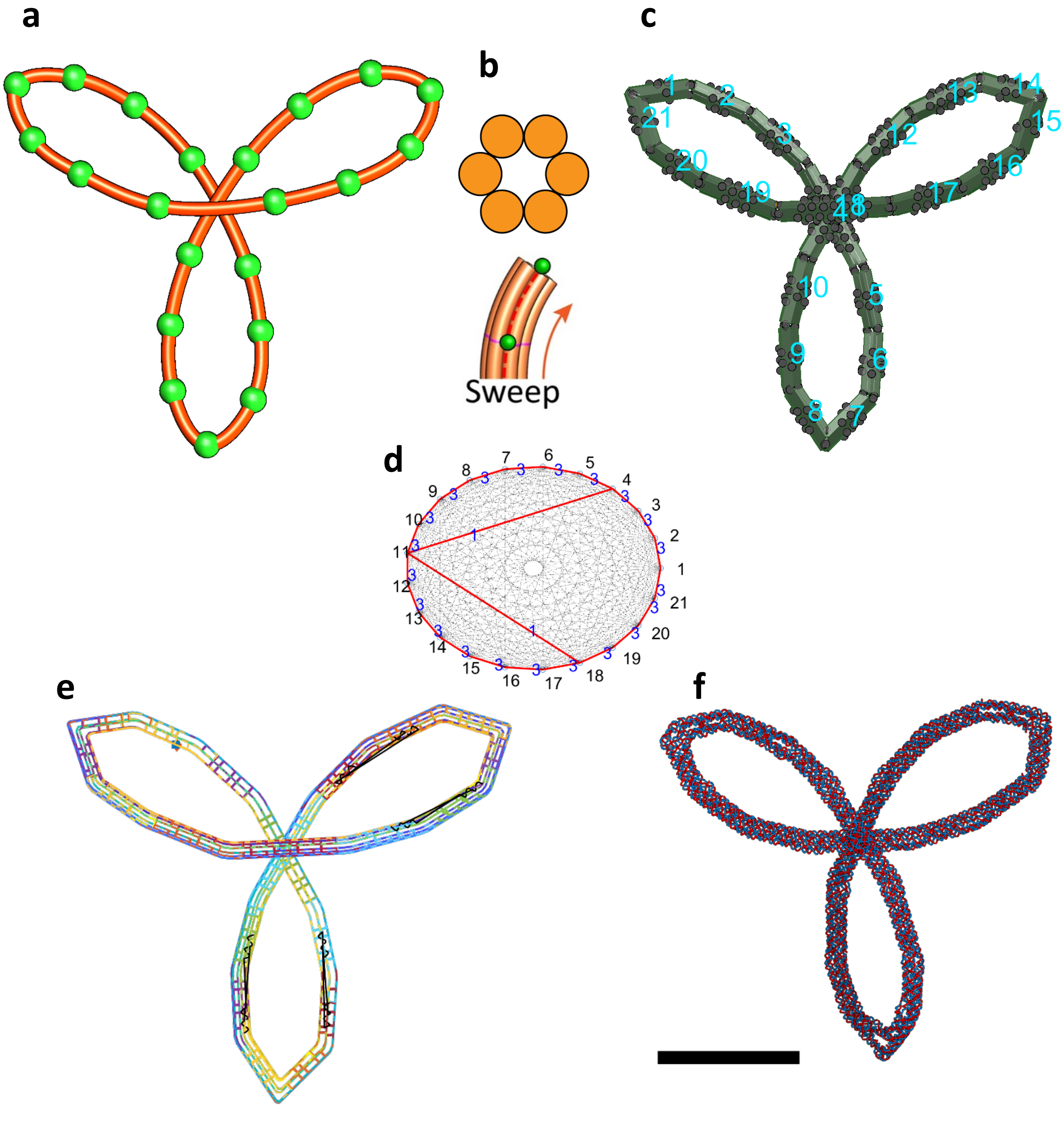


**Fig. S30.**

**Design details of the Trifolium structure.** (a) Line sketch of target structure. (b) Cross-section of each line. (c) Assembly model consisting of 21 bundles. (d) Connectivity matrix for assembly and routing algorithms. (e) Final scaffold and staple routings in the 3D representation. (f) oxDNA simulation result for the mean configuration. Scale bar = 50 nm.

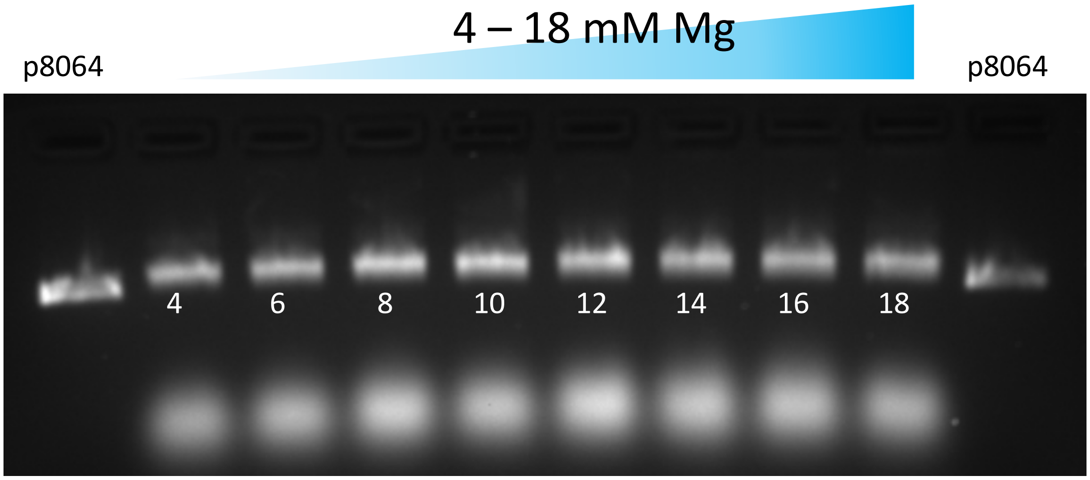


**Fig. S31.**

**Magnesium screening during the folding of the Trifolium structure.** Samples were run in a 1.5% agarose gel, using 45 mM Tris, 45 mM boric acid, 1 mM EDTA and 11 mM $MgCl_2$ as buffer. The gel was run at 90 V for 90 minutes and prestained with Ethidium bromide.

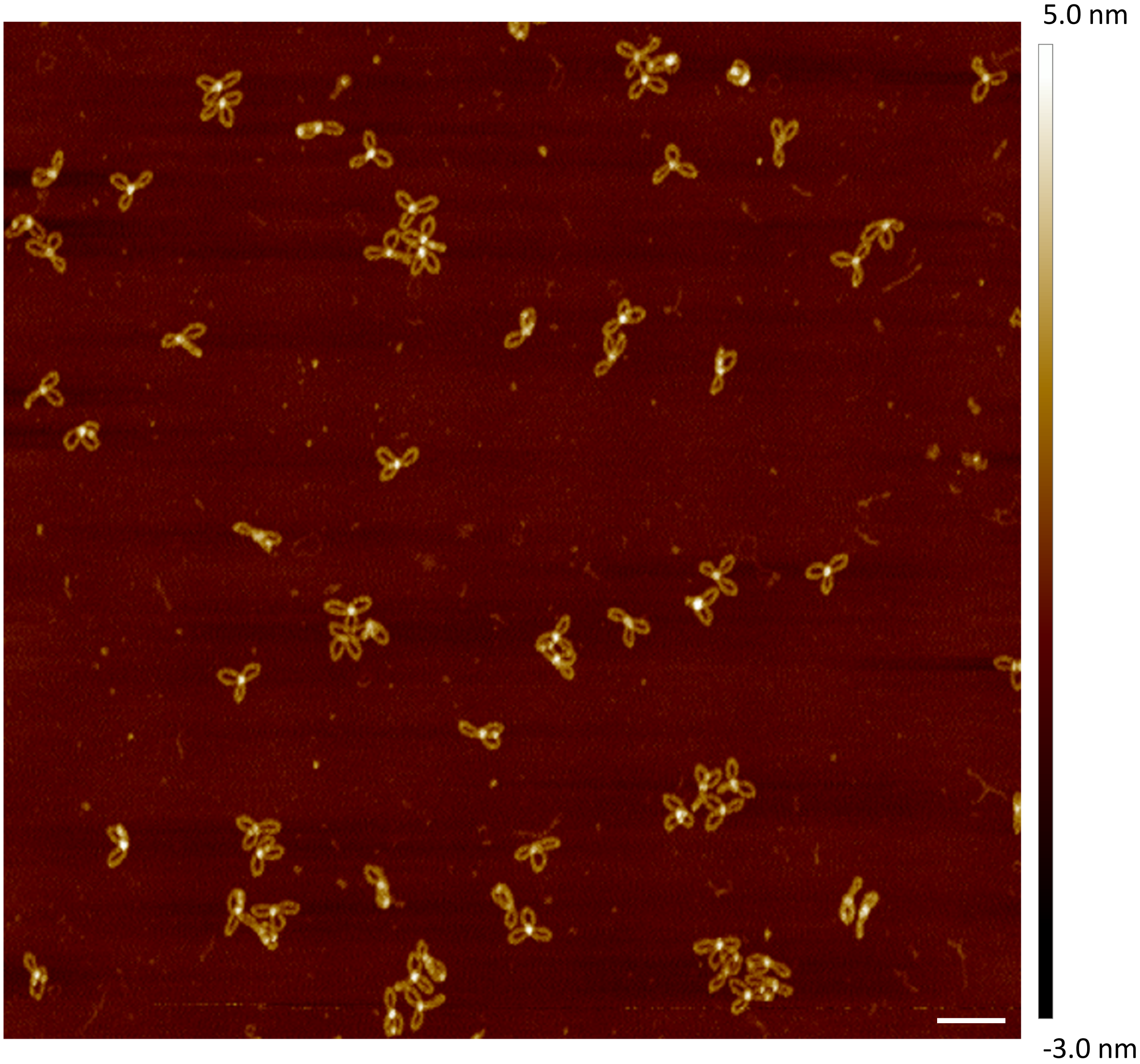


**Fig. S32.**

**AFM image of the Trifolium structure.** Freeze 'N Squeeze purified samples were incubated on freshly cleaved mica for 3 minutes, washed briefly with $H_2O$ and subsequently dried with a gentle flow of air. Samples were measured using ScanAsyst Air tips. Scale bar = 200 nm.

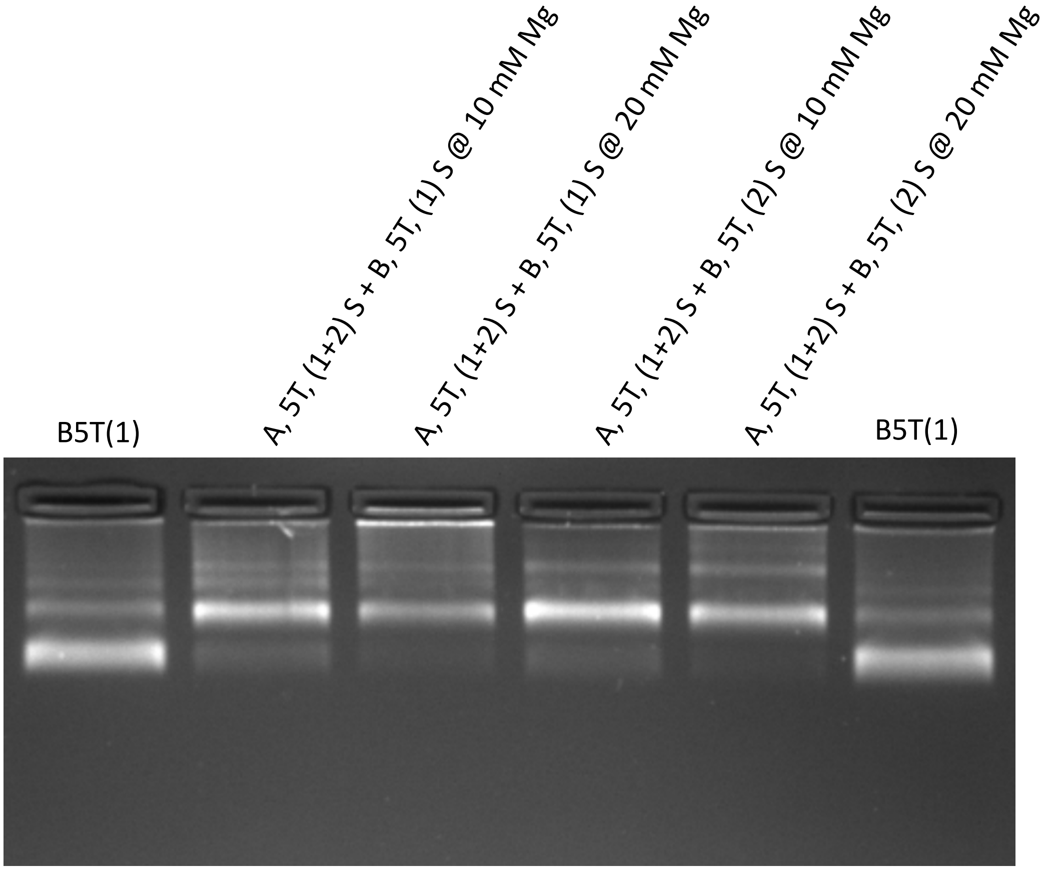


**Fig. S33.**

**Multimerization of Trifolium structures with overhangs Species A(1+2) and overhangs Species B(1) or Species B(2).** Multimerization was performed at different Magnesium concentrations for 20 h at 40°C. The monomeric structure with overhang set B(1) was used to indicate the running of monomeric species. Samples were run in a 1.5% agarose gel, using 45 mM Tris, 45 mM boric acid, 1 mM EDTA and 11 mM $MgCl_2$ as buffer. The gel was run at 90 V for 90 minutes and prestained with Ethidium bromide.

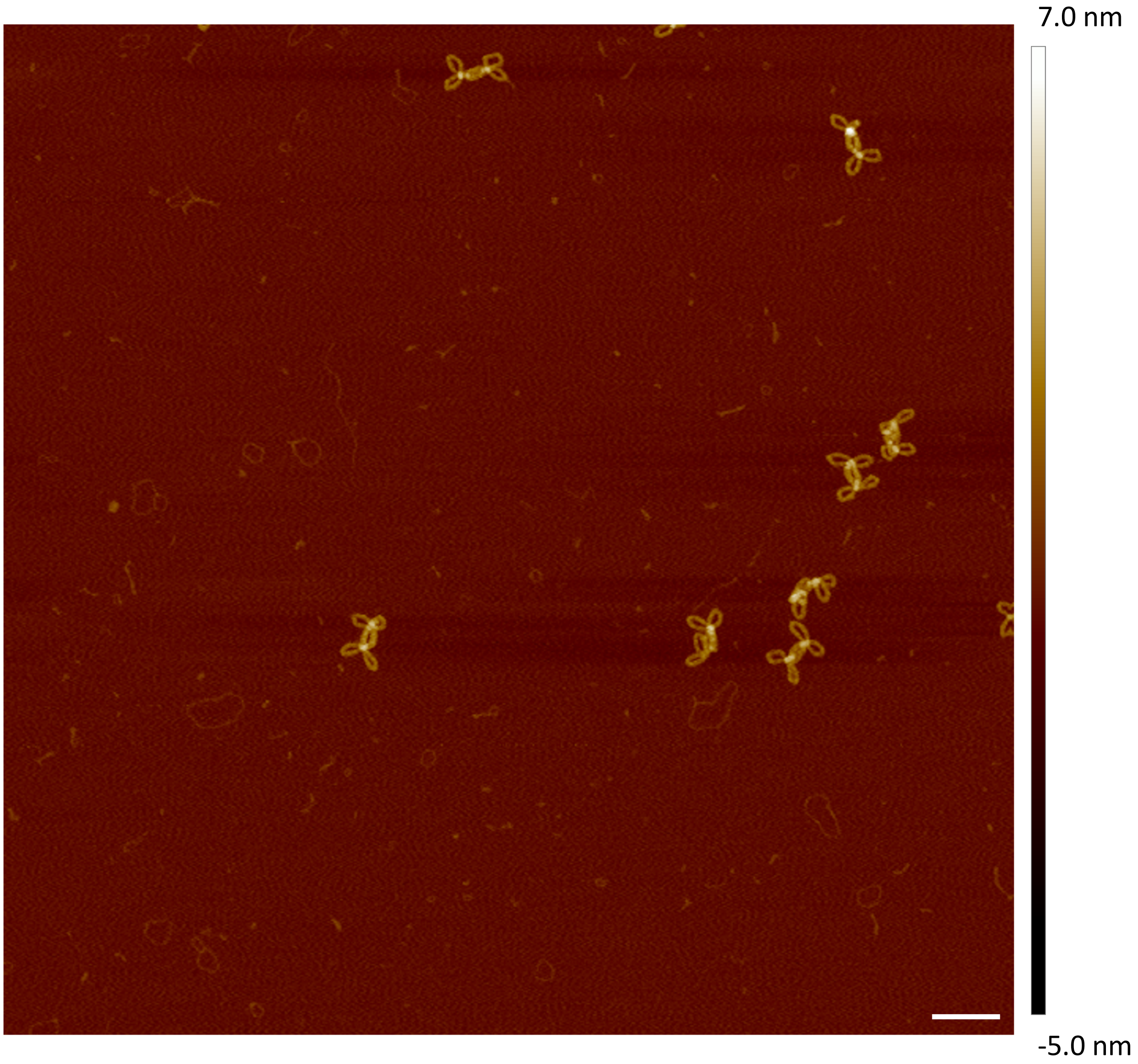


**Fig. S34.**

**AFM image of the Trifolium dimer-structures.** Freeze 'N Squeeze purified samples were incubated on freshly cleaved mica for 3 minutes, washed briefly with $H_2O$ and subsequently dried with a gentle flow of air. Samples were measured using ScanAsyst Air tips. Scale bar = 200 nm.

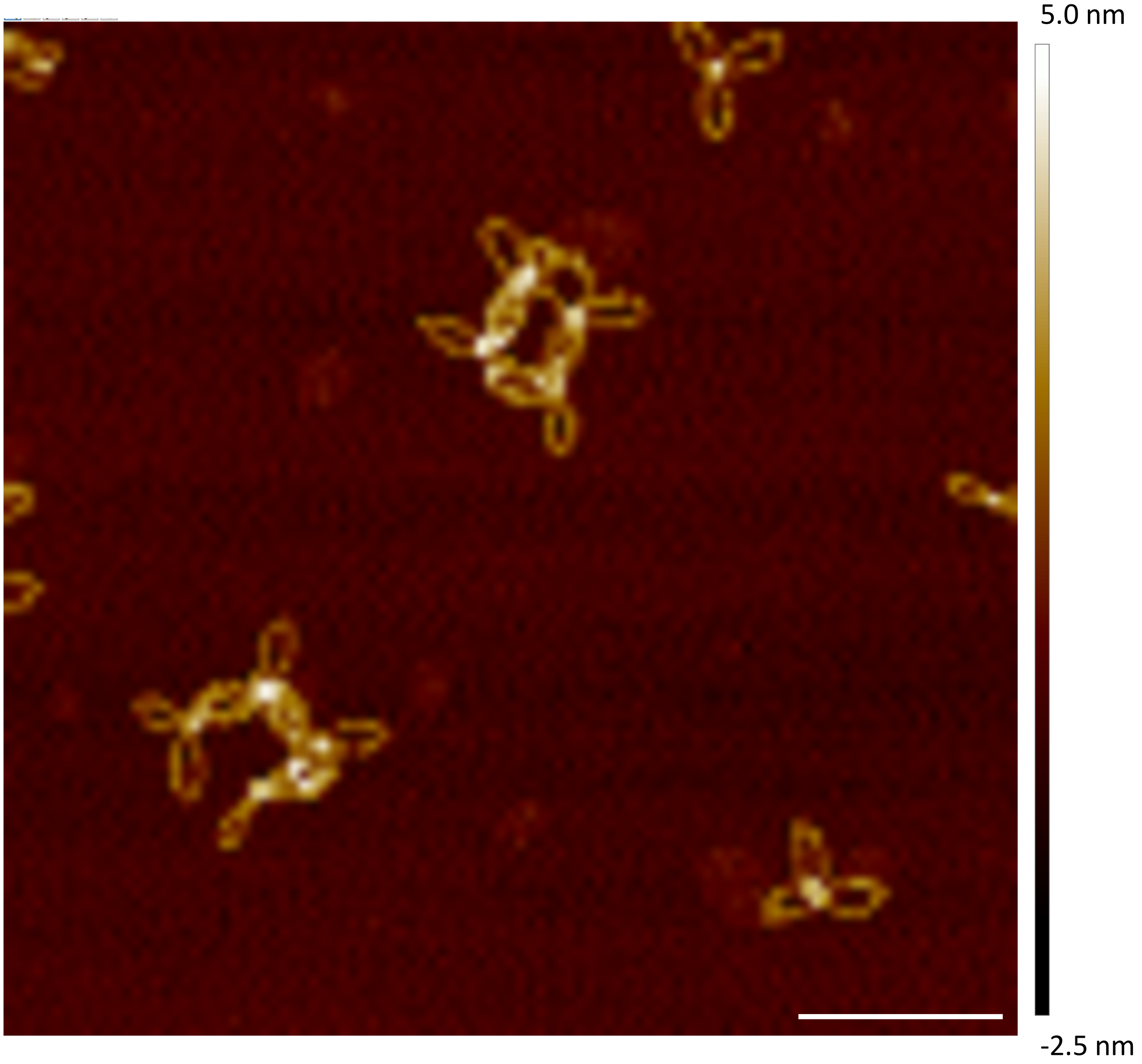


**Fig. S35.**

**AFM image of higher order Trifolium structures.** Samples were incubated on freshly cleaved mica for 3 minutes, washed briefly with $H_2O$ and subsequently dried with a gentle flow of air. Samples were measured using ScanAsyst Air tips. Scale bar = 200 nm.

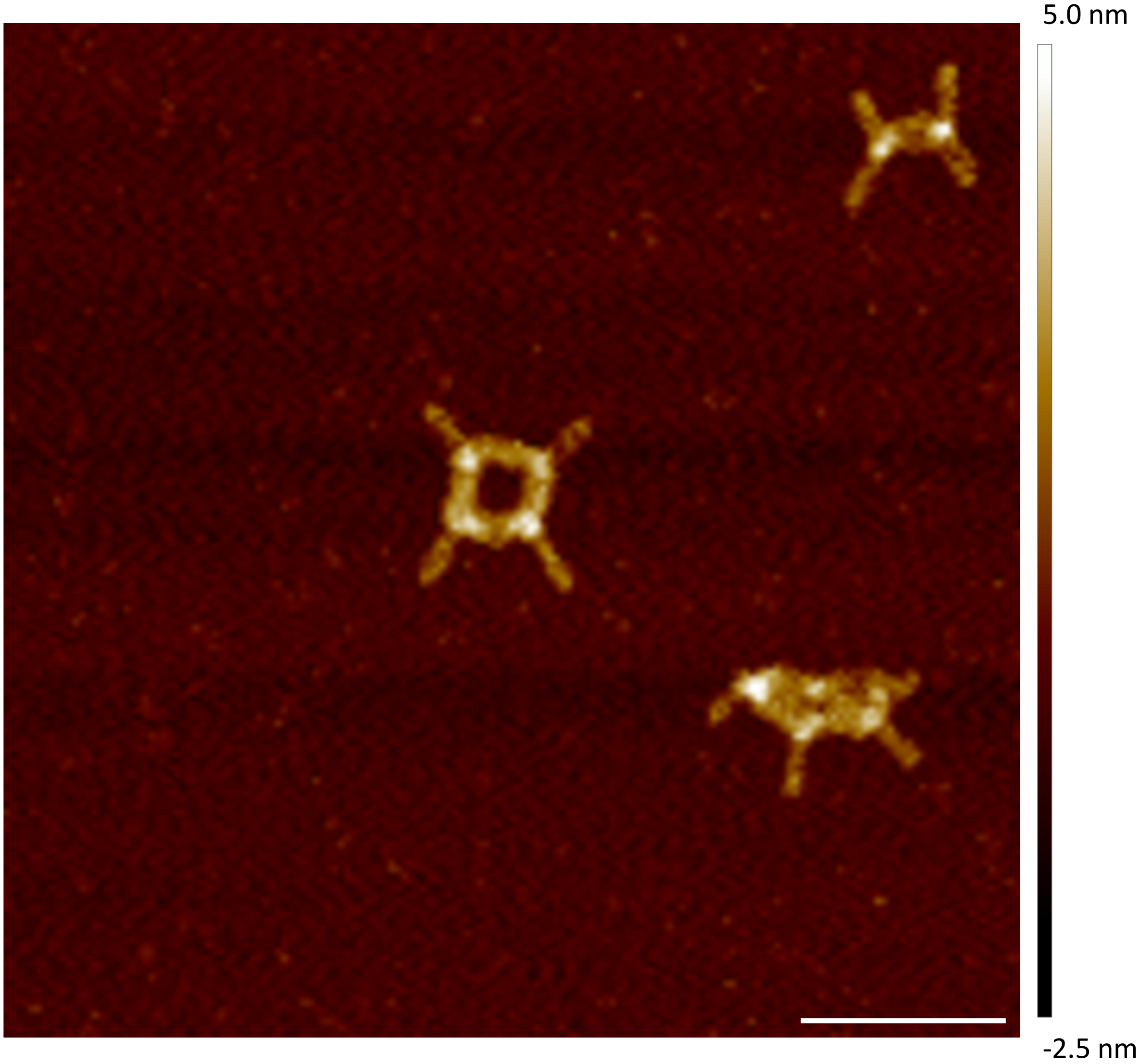


**Fig. S36.**

**AFM image of higher order Trifolium structures.** Samples were incubated on freshly cleaved mica for 3 minutes, washed briefly with $H_2O$ and subsequently dried with a gentle flow of air. Samples were measured using ScanAsyst Air tips. Scale bar = 200 nm.

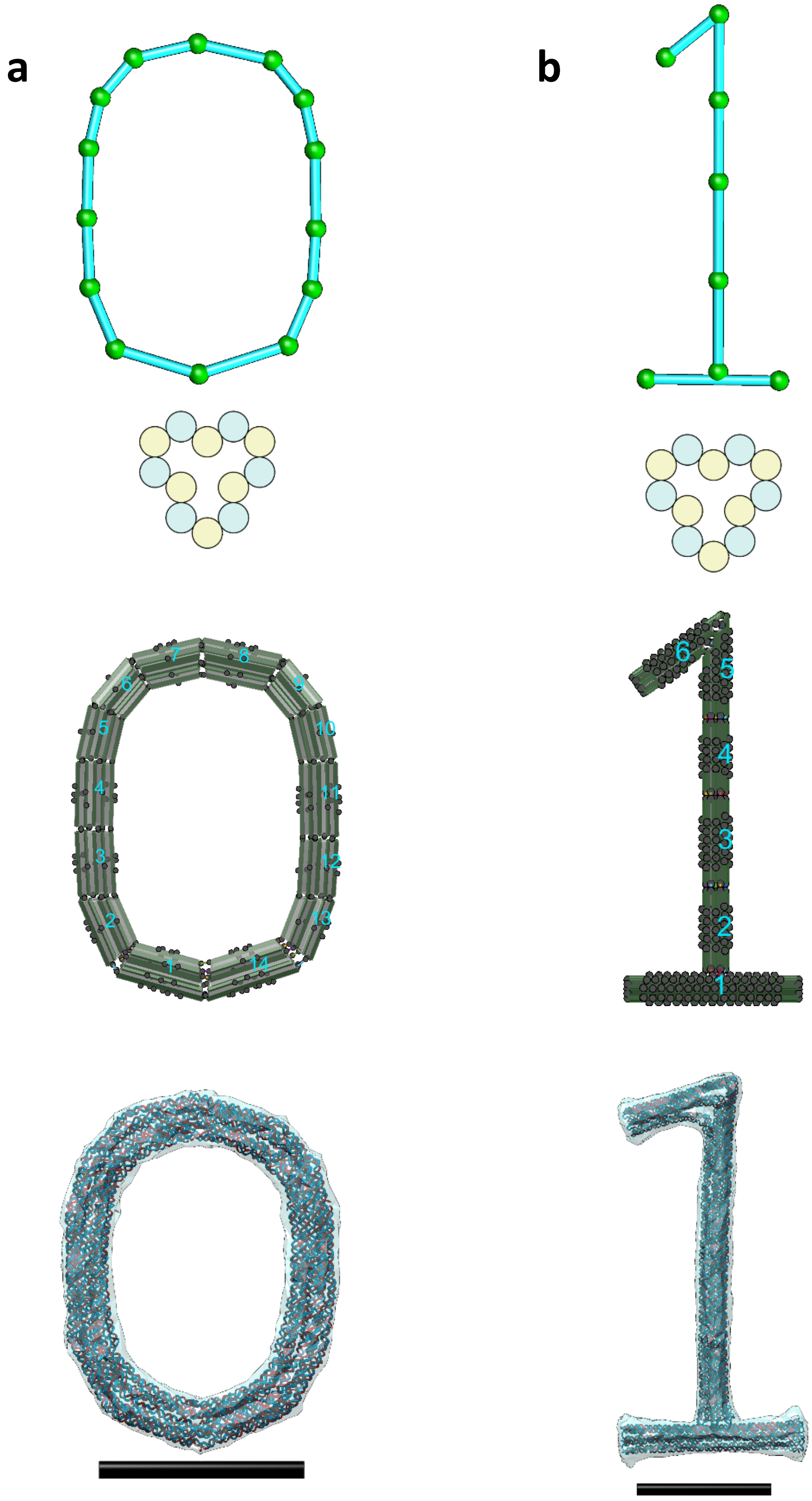


**Fig. S37.**

**Other freeform examples in the number series (0 & 1).** (a) A design of “0”. (b) A design of “1”. From Top to bottom, line sketch, cross-section, assembly model, and oxDNA simulation result. Scale bars = 50 nm.

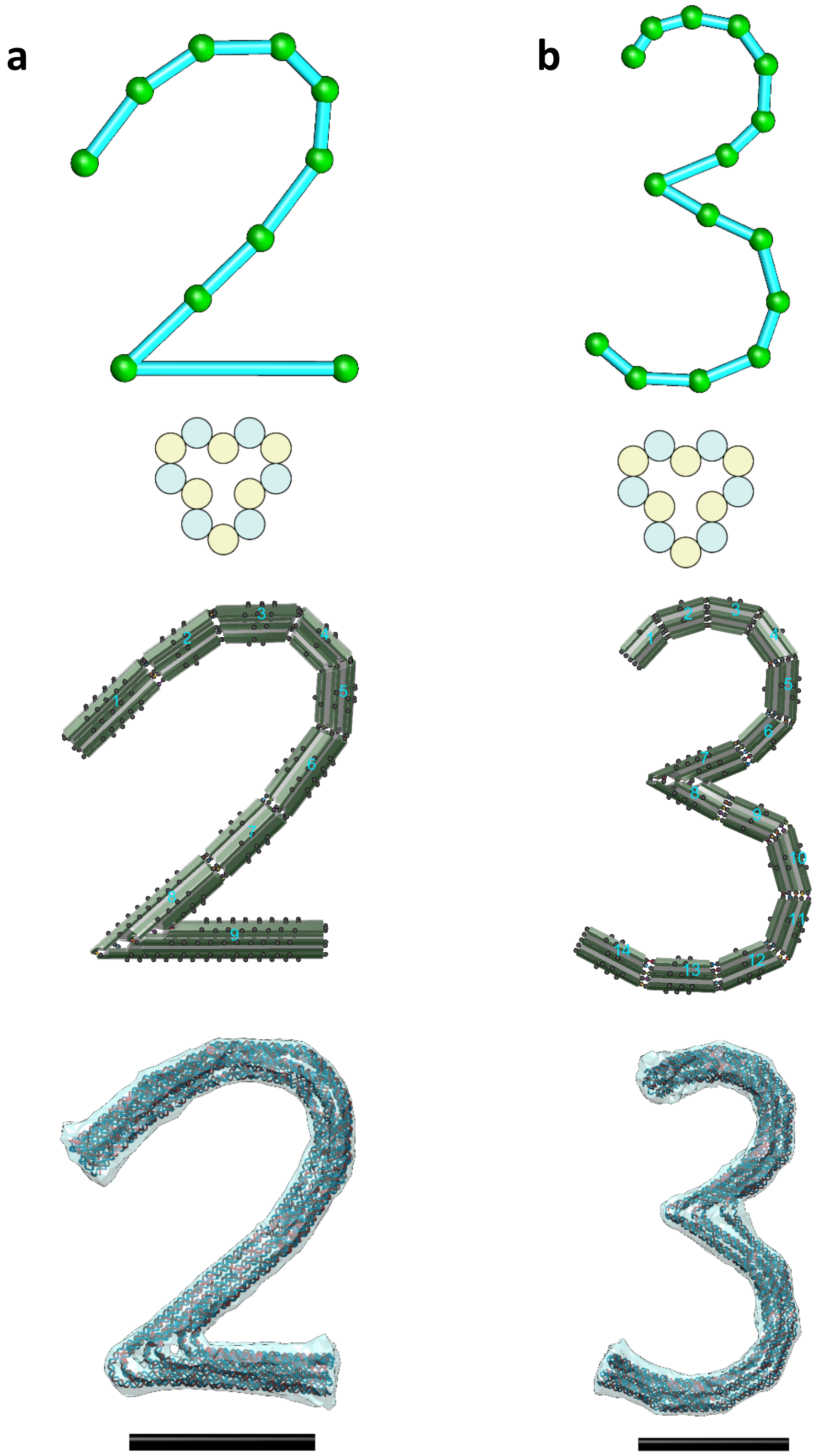


**Fig. S38.**

**Other freeform examples in the number series (2 & 3).** (a) A design of “2”. (b) A design of “3”. From Top to bottom, line sketch, cross-section, assembly model, and oxDNA simulation result. Scale bars = 50 nm.

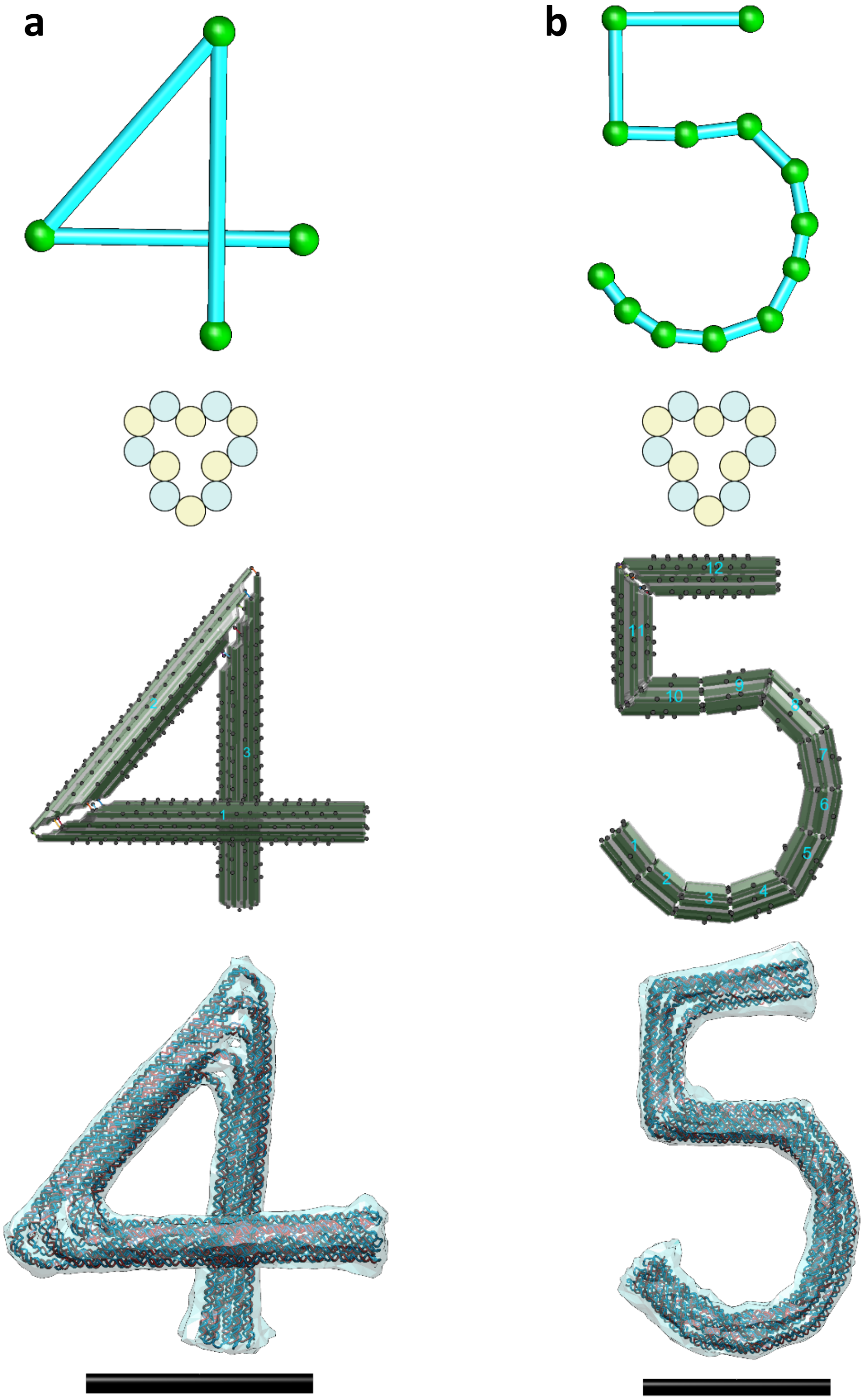


**Fig. S39.**
**Other freeform examples in the number series (4 & 5).** (a) A design of "4". (b) A design of "5". From Top to bottom, line sketch, cross-section, assembly model, and oxDNA simulation result. Scale bars = 50 nm.

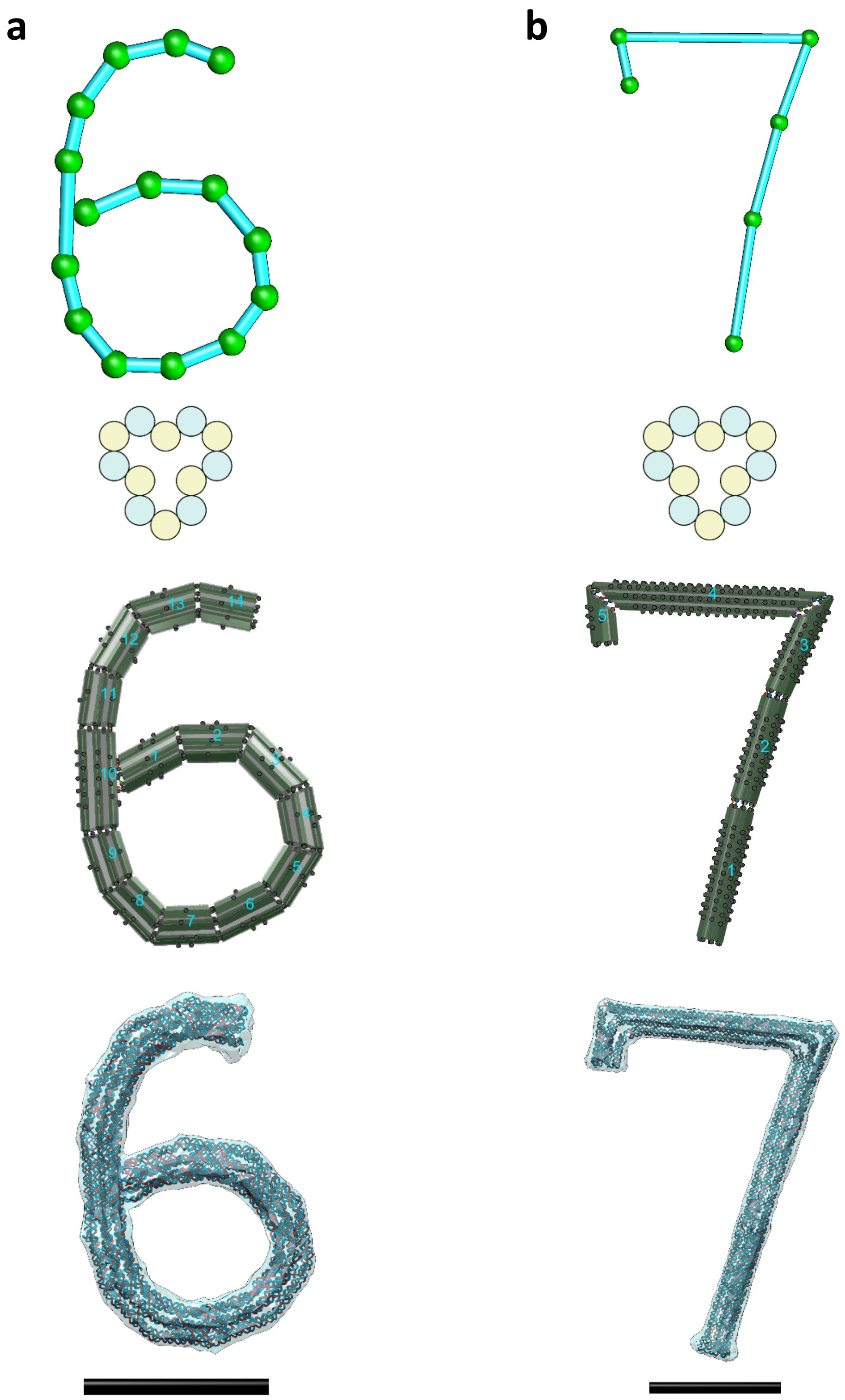


**Fig. S40.**

**Other freeform examples in the number series (6 & 7).** (a) A design of “6”. (b) A design of “7”. From Top to bottom, line sketch, cross-section, assembly model, and oxDNA simulation result. Scale bars = 50 nm.

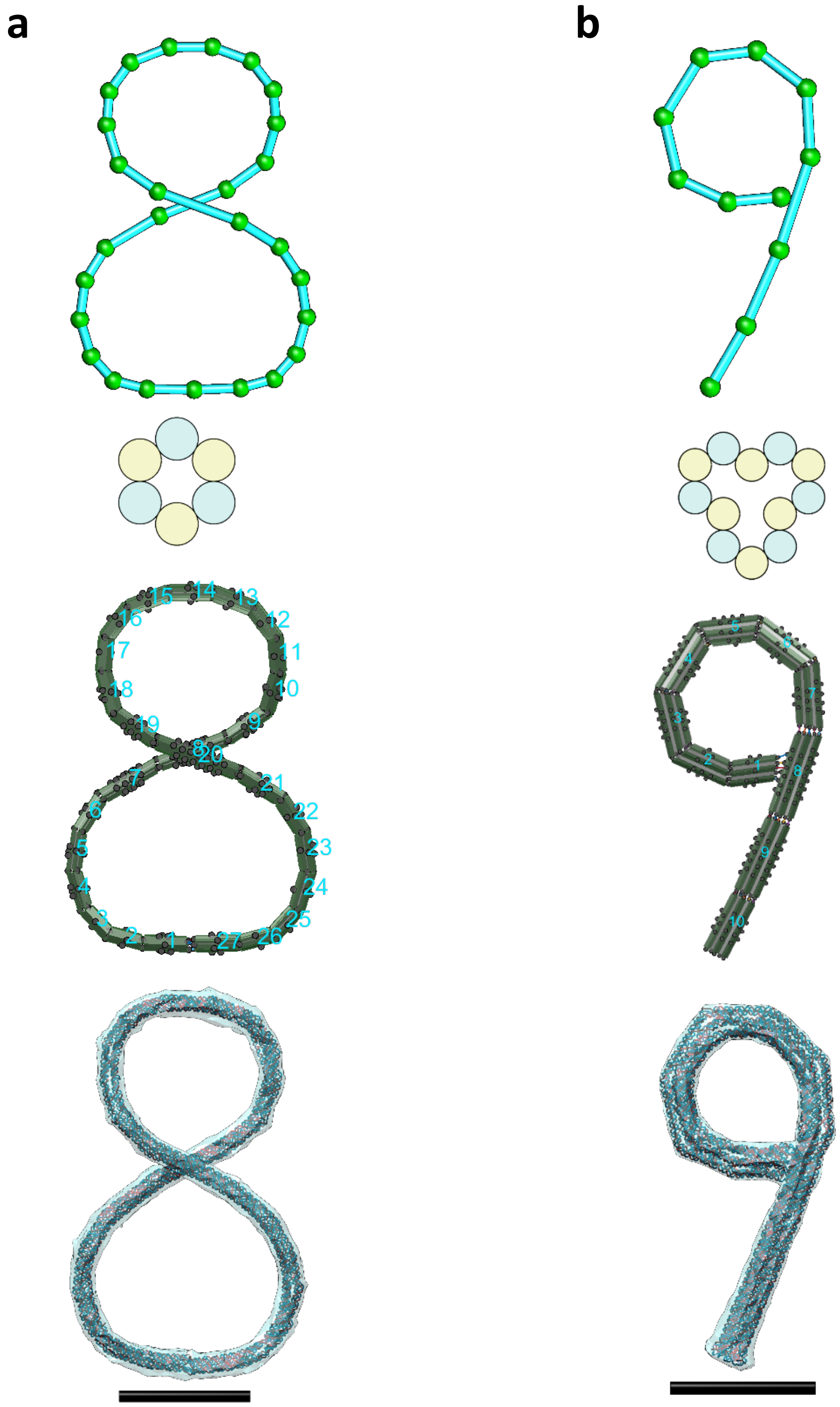


**Fig. S41.**

**Other freeform examples in the number series (8 & 9).** (a) A design of “8”. (b) A design of “9”. From Top to bottom, line sketch, cross-section, assembly model, and oxDNA simulation result. Scale bars = 50 nm.

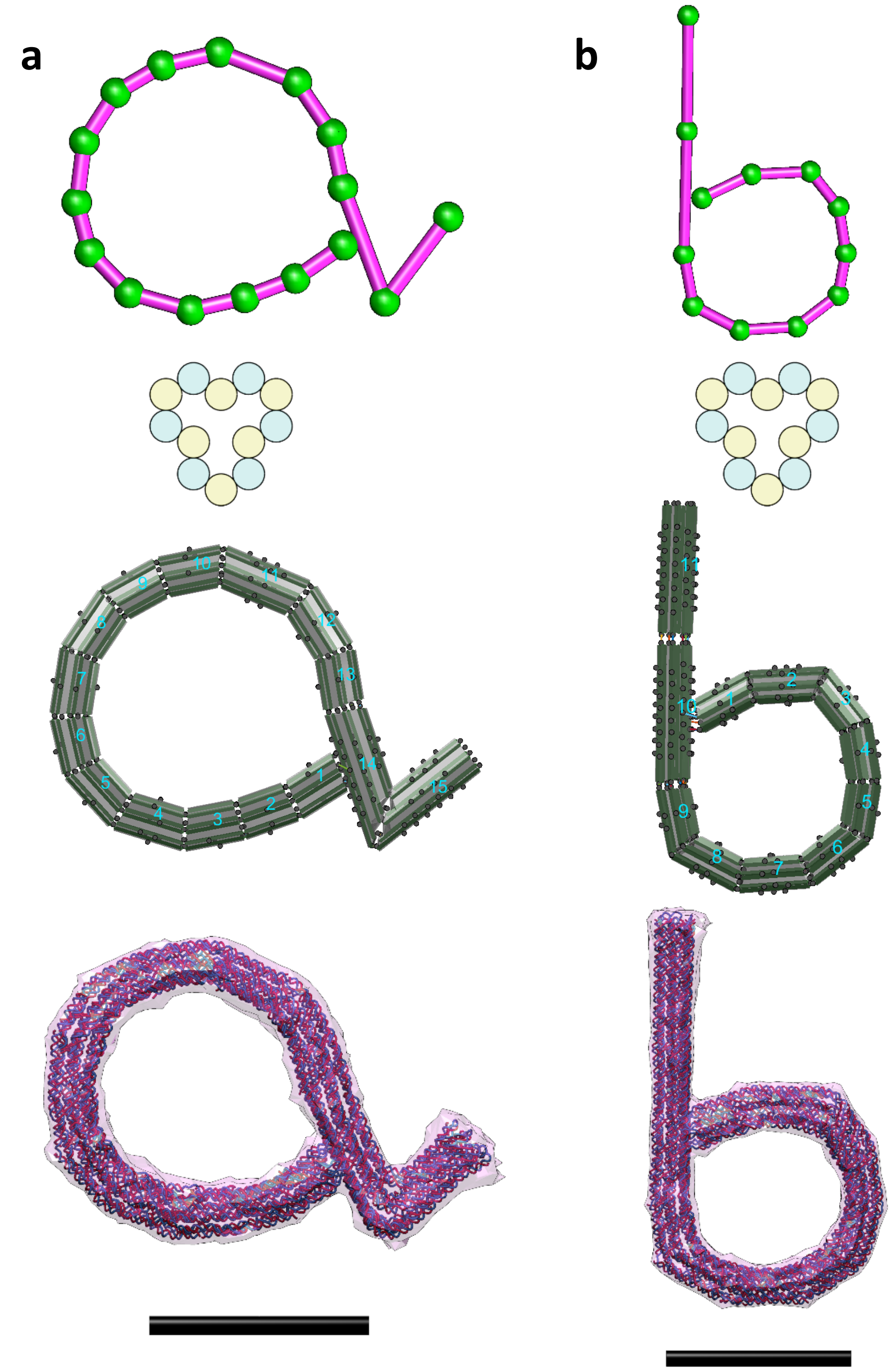


**Fig. S42.**

**Other freeform examples in the lowercase series (a & b).** (a) A design of “a”. (b) A design of “b”. From Top to bottom, line sketch, cross-section, assembly model, and oxDNA simulation result. Scale bars = 50 nm.

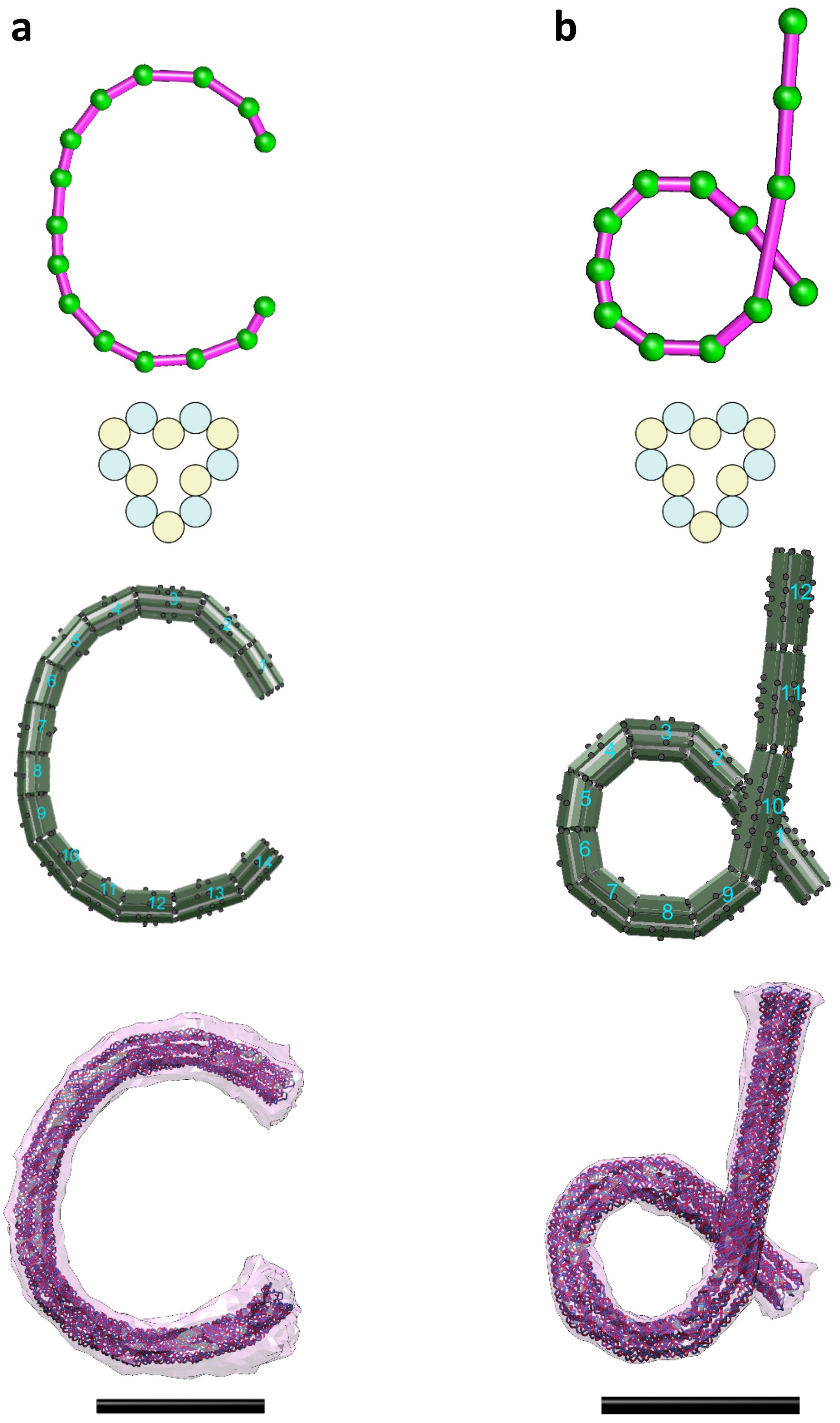


**Fig. S43.**

**Other freeform examples in the lowercase series (c & d).** (a) A design of "c". (b) A design of "d". From Top to bottom, line sketch, cross-section, assembly model, and oxDNA simulation result. Scale bars = 50 nm.

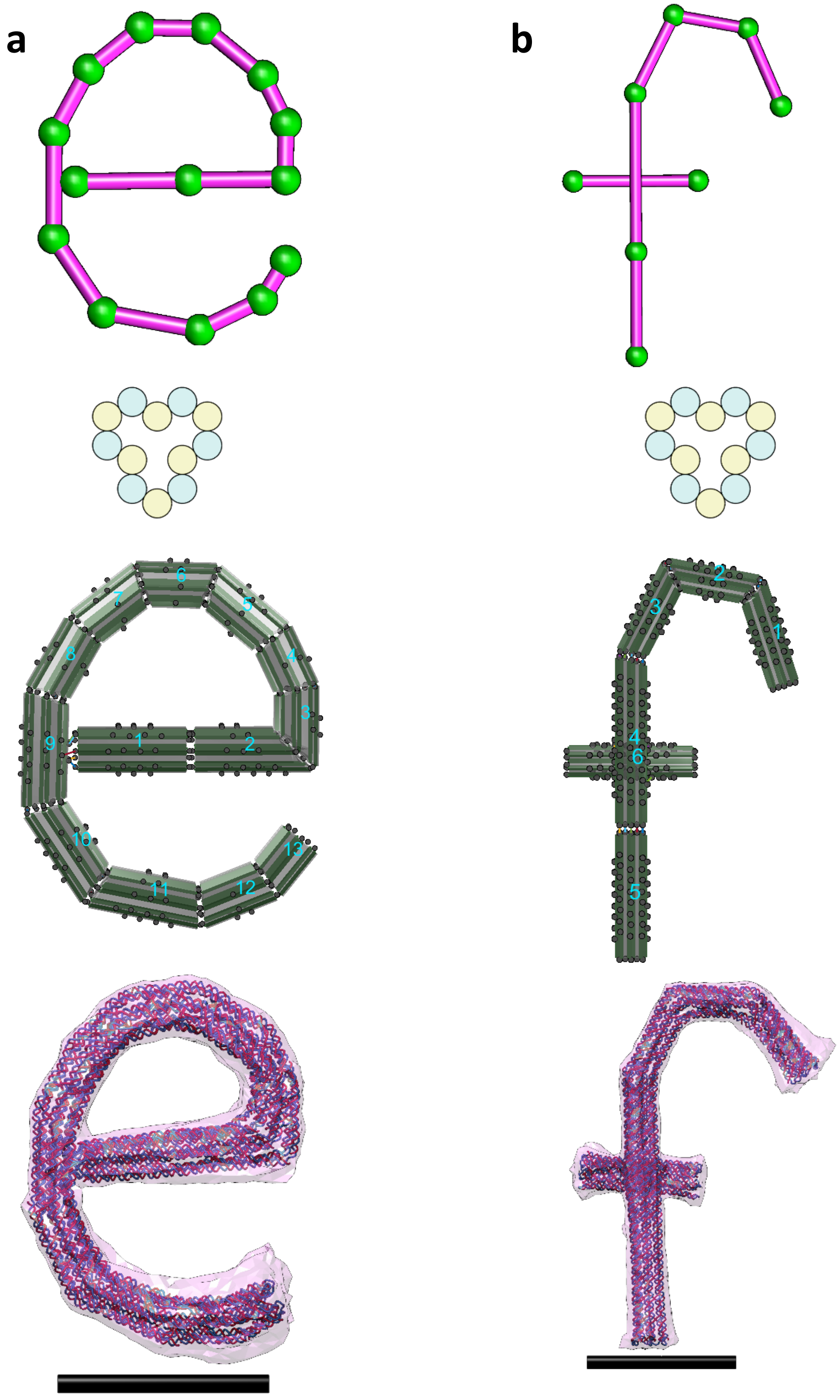


**Fig. S44.**

**Other freeform examples in the lowercase series (e & f).** (a) A design of "e". (b) A design of "f". From Top to bottom, line sketch, cross-section, assembly model, and oxDNA simulation result. Scale bars = 50 nm.

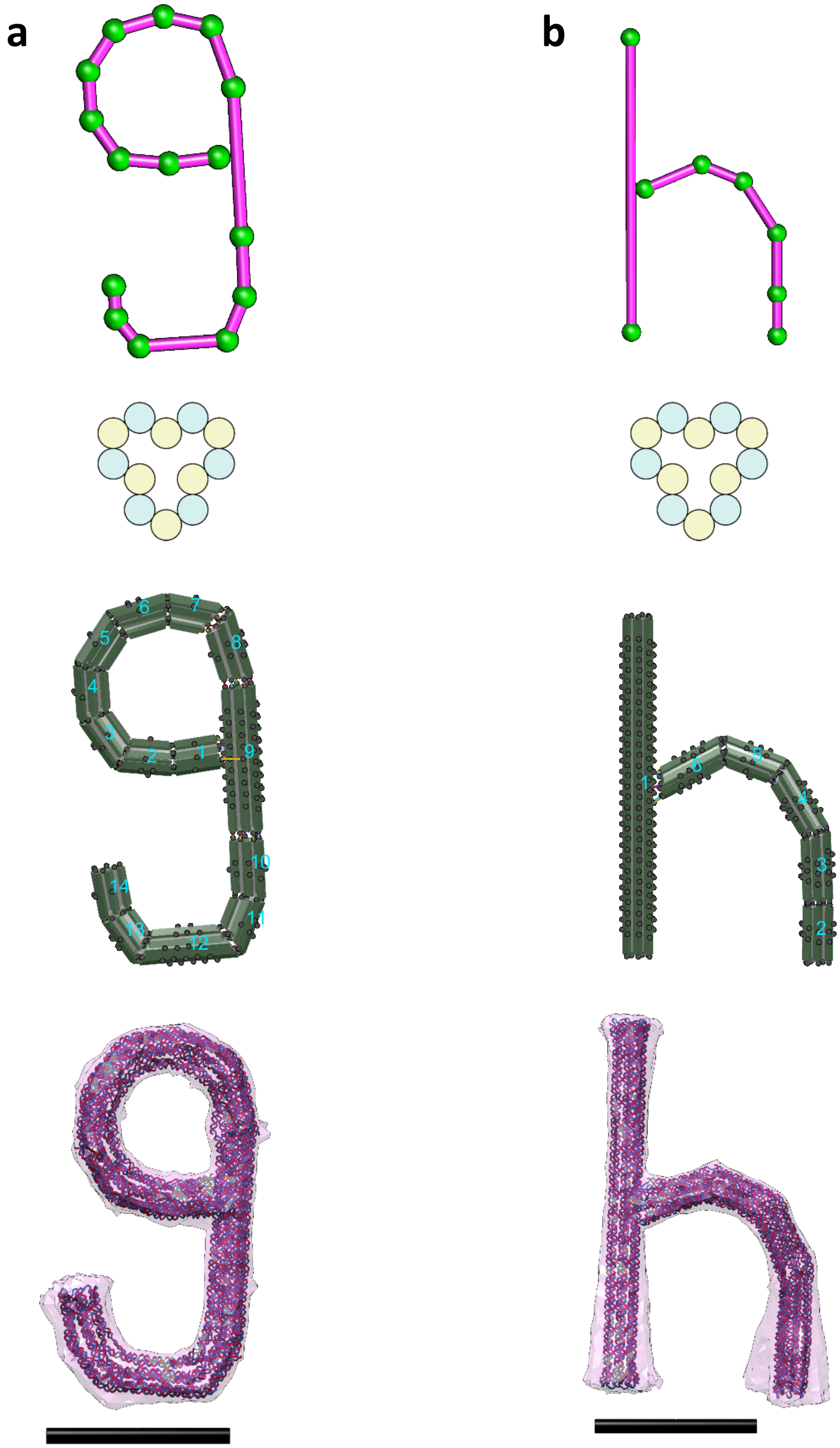


**Fig. S45.**

**Other freeform examples in the lowercase series (g & h).** (a) A design of "g". (b) A design of "h". From Top to bottom, line sketch, cross-section, assembly model, and oxDNA simulation result. Scale bars = 50 nm.

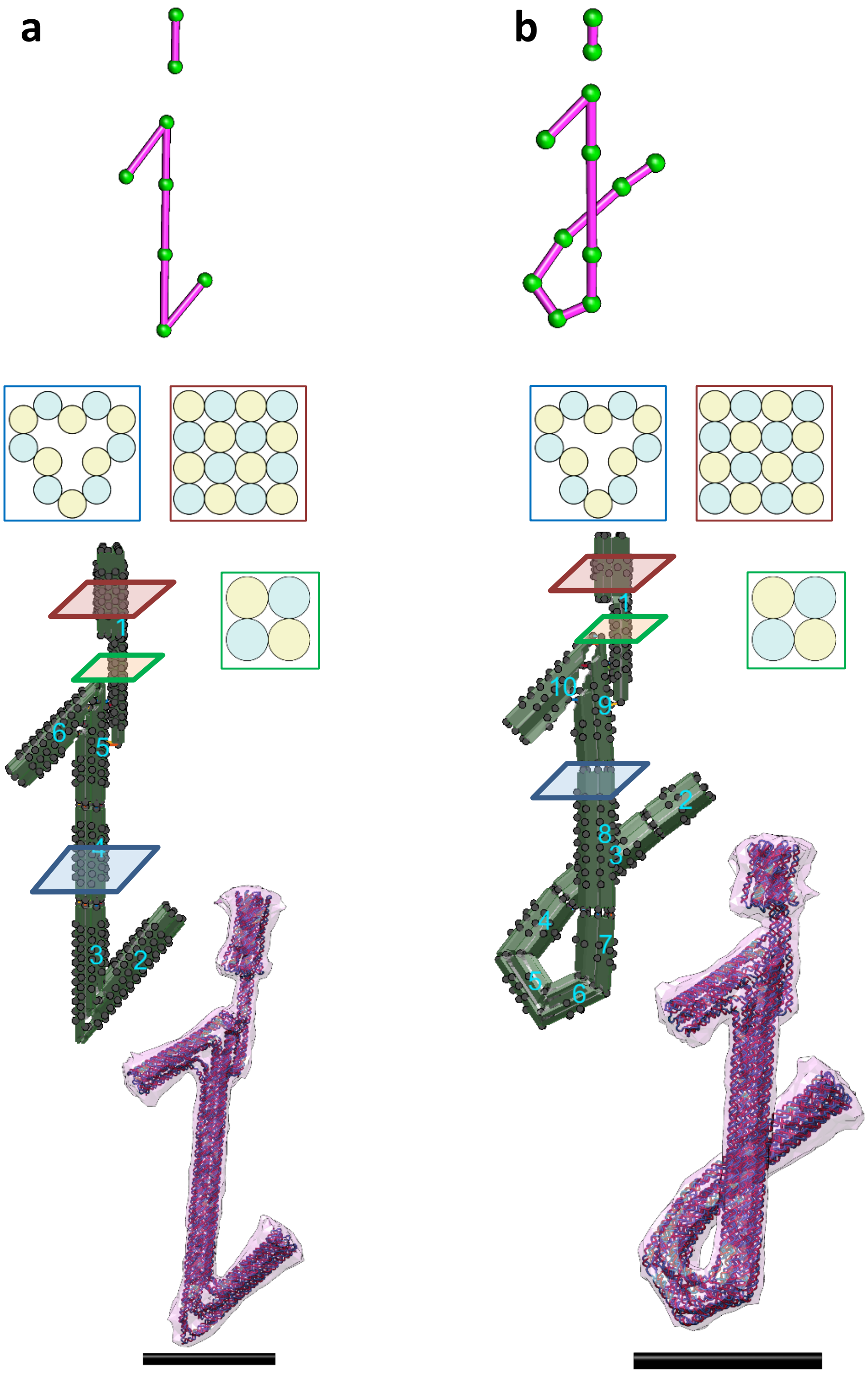


**Fig. S46.**

**Other freeform examples in the lowercase series (i & j).** (a) A design of "i". (b) A design of "j". From Top to bottom, line sketch, cross-section, assembly model, and oxDNA simulation result. Note that these two designs were involved the use of the bundle editing GUI to adjust individual cylinder lengths such that the four cylinders in the first bundle is extended to connect to the bottom bundles. Scale bars = 50 nm.

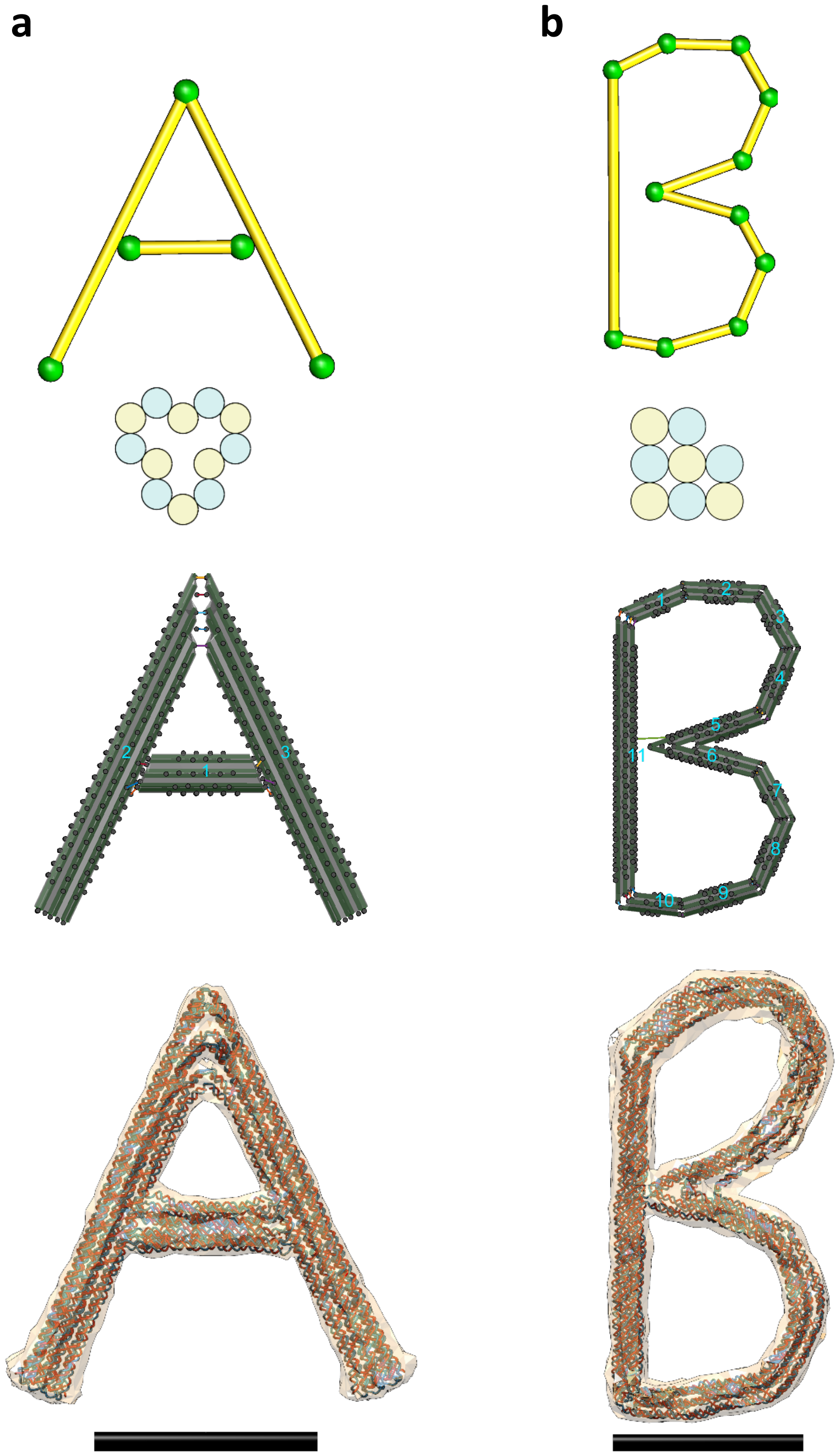


**Fig. S47.**

**Other freeform examples in the uppercase series (A & B).** (a) A design of "A". (b) A design of "B". From Top to bottom, line sketch, cross-section, assembly model, and oxDNA simulation result. Scale bars = 50 nm.

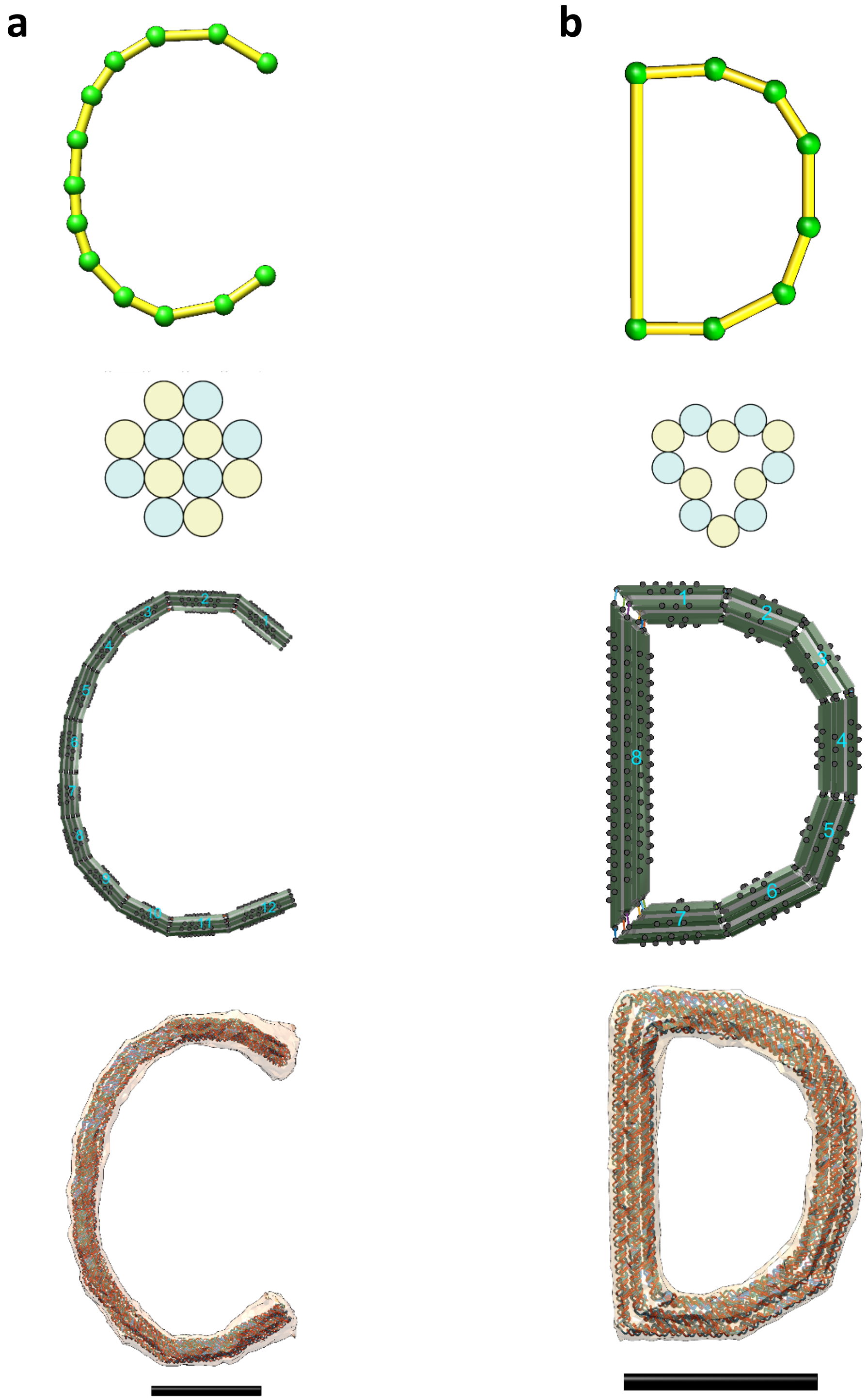


**Fig. S48.**

**Other freeform examples in the uppercase series (C & D).** (a) A design of "C". (b) A design of "D". From Top to bottom, line sketch, cross-section, assembly model, and oxDNA simulation result. Scale bars = 50 nm.

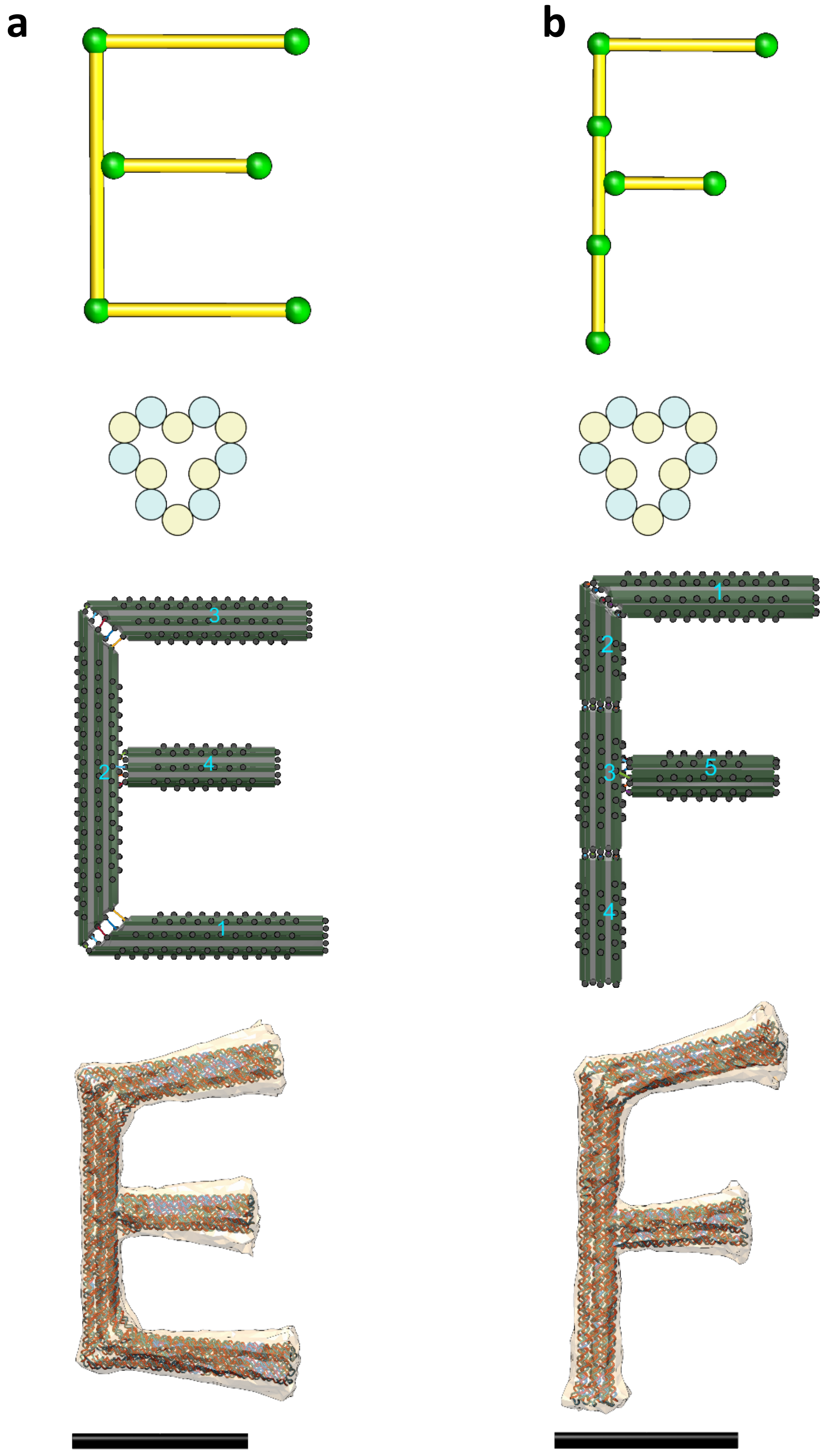


**Fig. S49.**

**Other freeform examples in the uppercase series (E & F).** (a) A design of "E". (b) A design of "F". From Top to bottom, line sketch, cross-section, assembly model, and oxDNA simulation result. Scale bars = 50 nm.

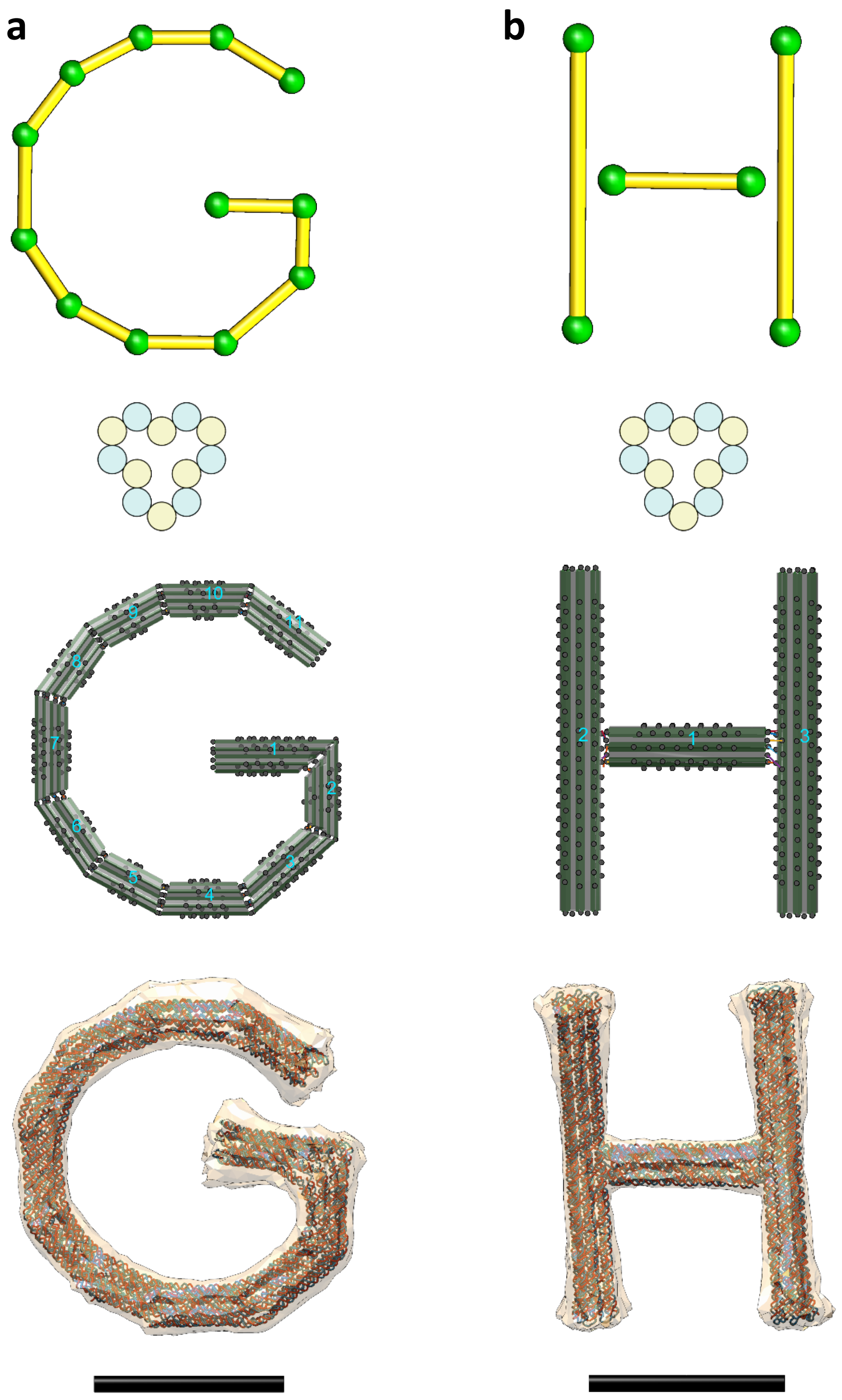


**Fig. S50.**

**Other freeform examples in the uppercase series (G & H).** (a) A design of "G". (b) A design of "H". From Top to bottom, line sketch, cross-section, assembly model, and oxDNA simulation result. Scale bars = 50 nm.

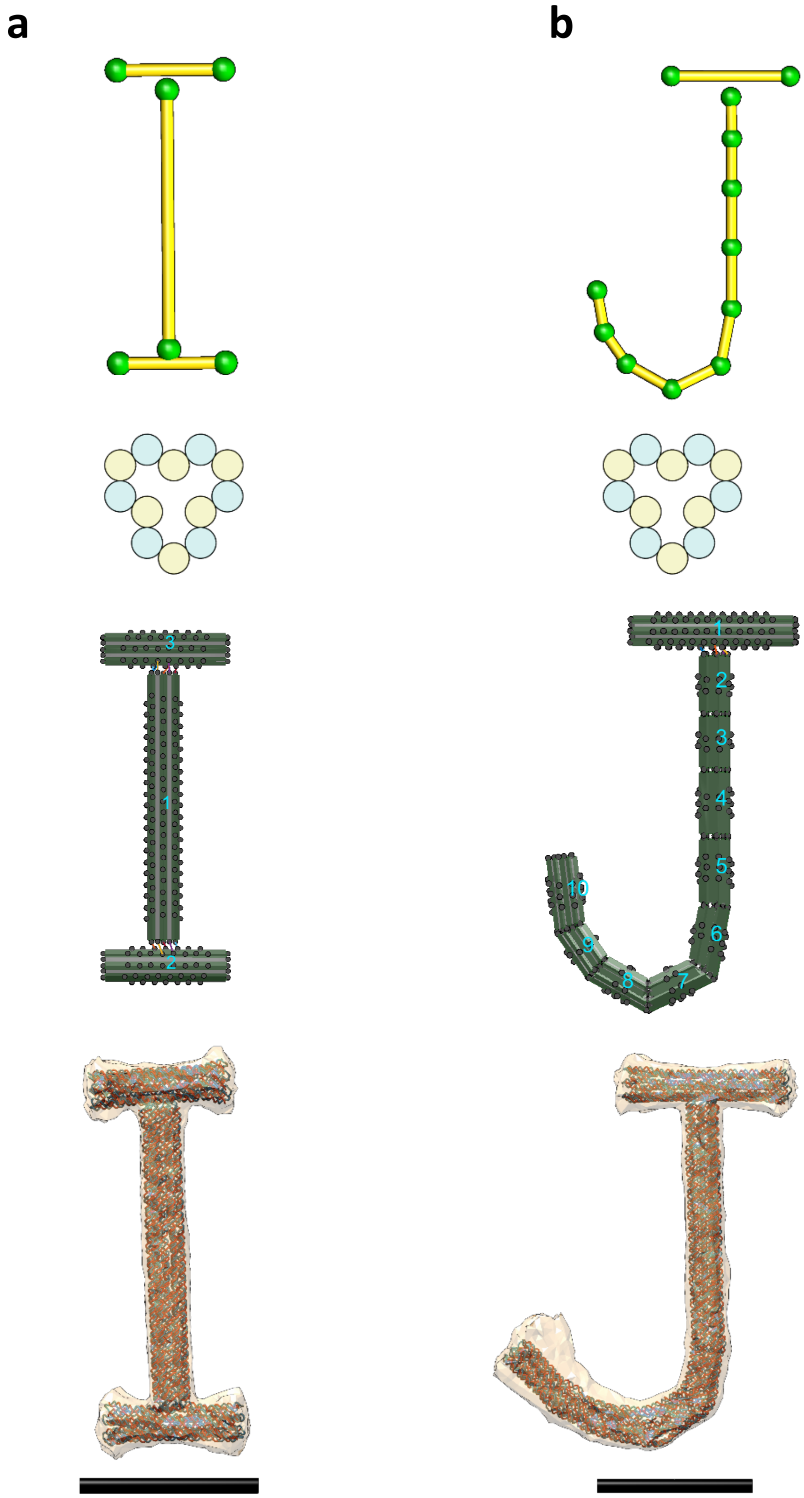


**Fig. S51.**

**Other freeform examples in the uppercase series (I & J).** (a) A design of "I". (b) A design of "J". From Top to bottom, line sketch, cross-section, assembly model, and oxDNA simulation result. Scale bars = 50 nm.

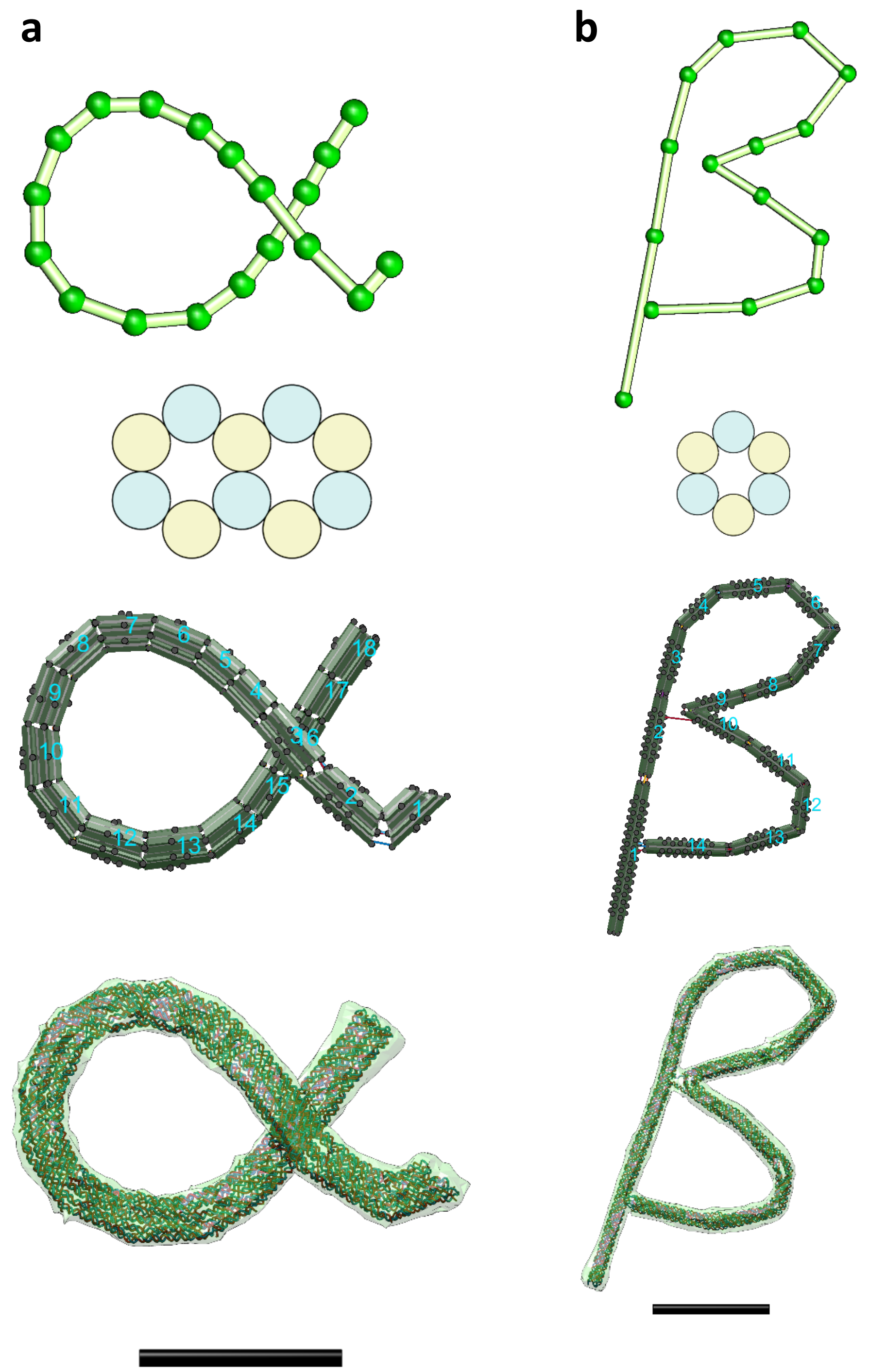


**Fig. S52.**

**Other freeform examples in the symbol series (alpha & beta).** (a) A design of “alpha”. (b) A design of “beta”. From Top to bottom, line sketch, cross-section, assembly model, and oxDNA simulation result. Scale bars = 50 nm.

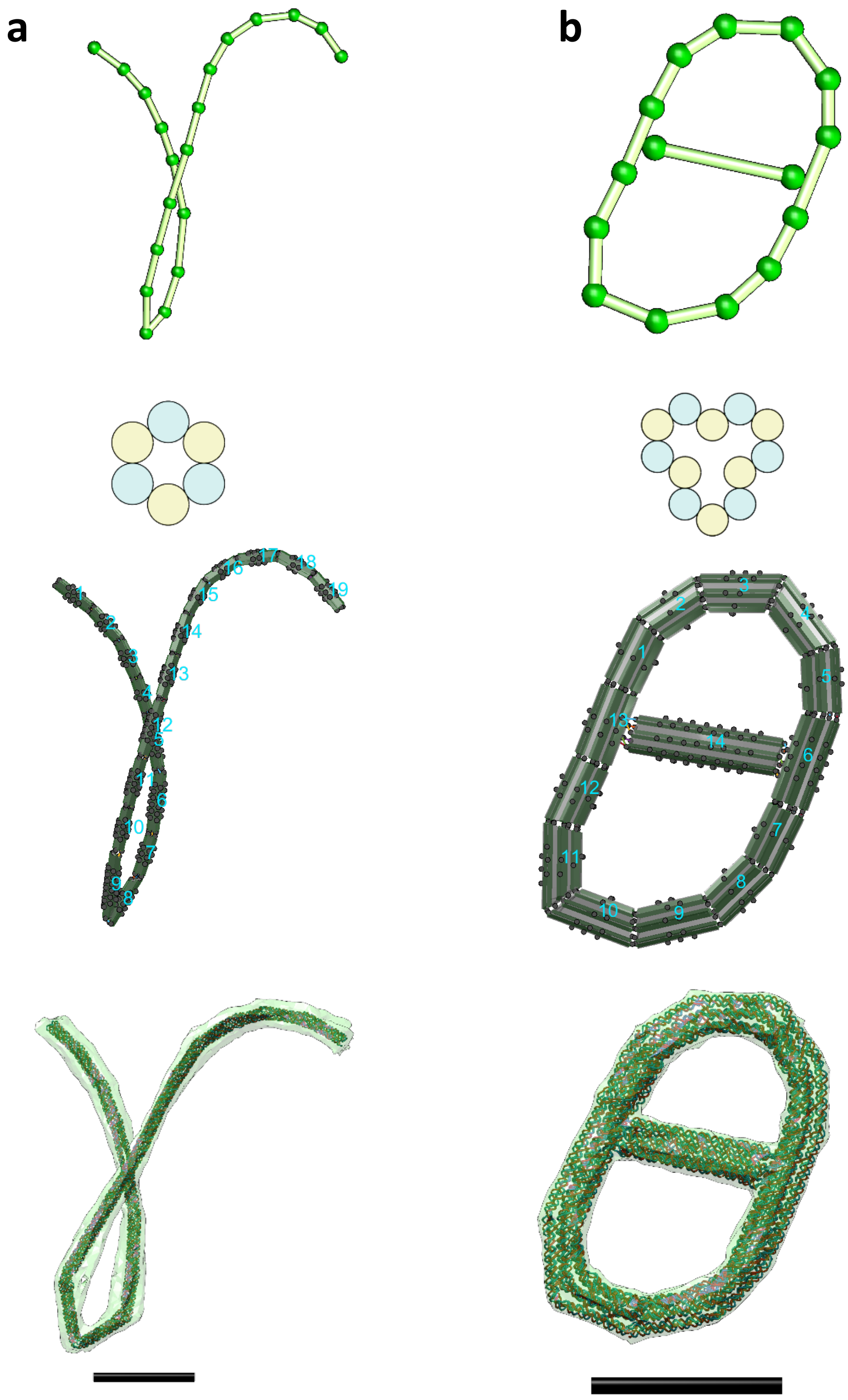


**Fig. S53.**

**Other freeform examples in the symbol series (gamma & theta).** (a) A design of "gamma". (b) A design of "theta". From Top to bottom, line sketch, cross-section, assembly model, and oxDNA simulation result. Scale bars = 50 nm.

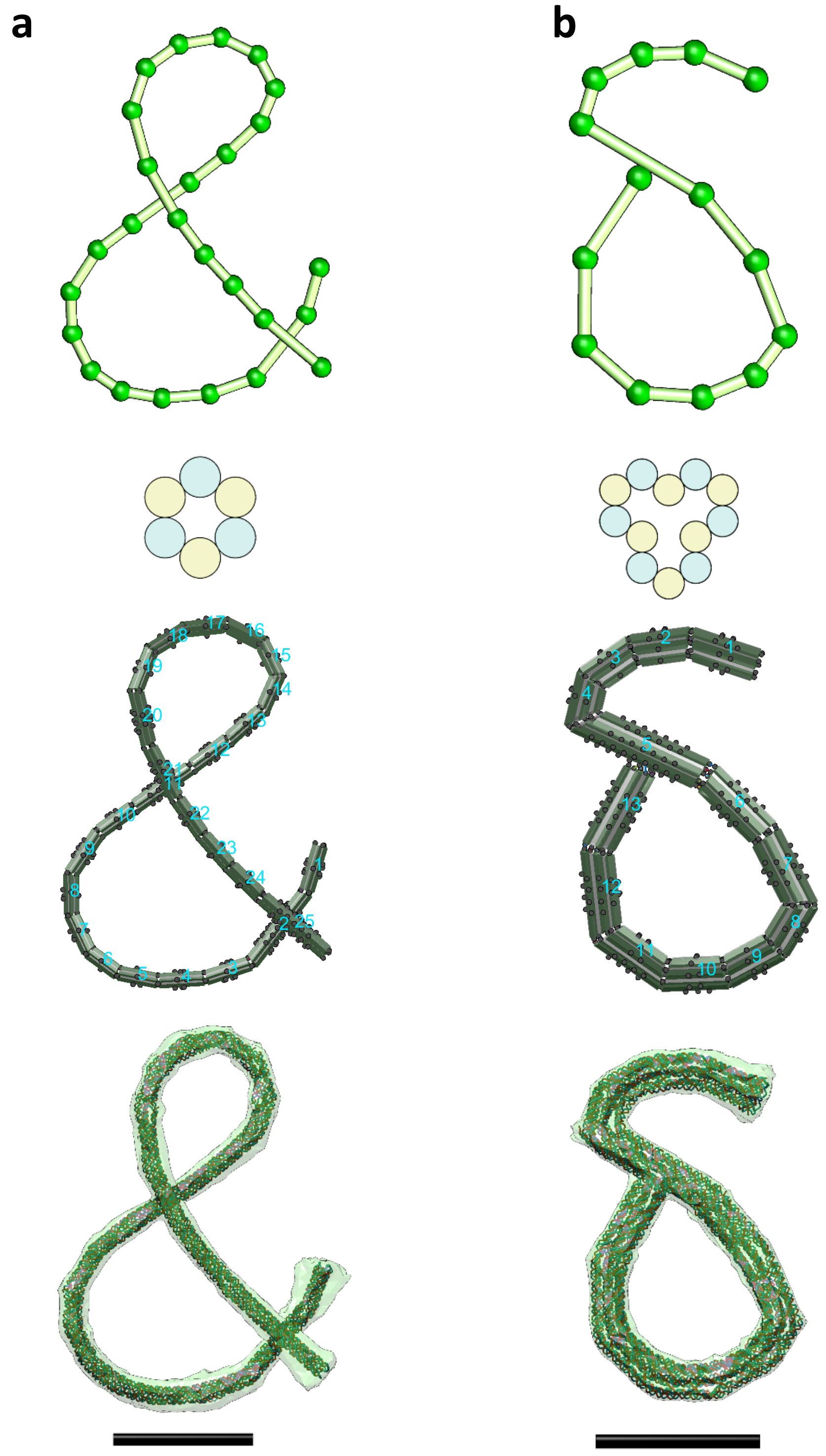


**Fig. S54.**

**Other freeform examples in the symbol series (ampersand & delta).** (a) A design of “&”. (b) A design of “delta”. From Top to bottom, line sketch, cross-section, assembly model, and oxDNA simulation result. Scale bars = 50 nm.

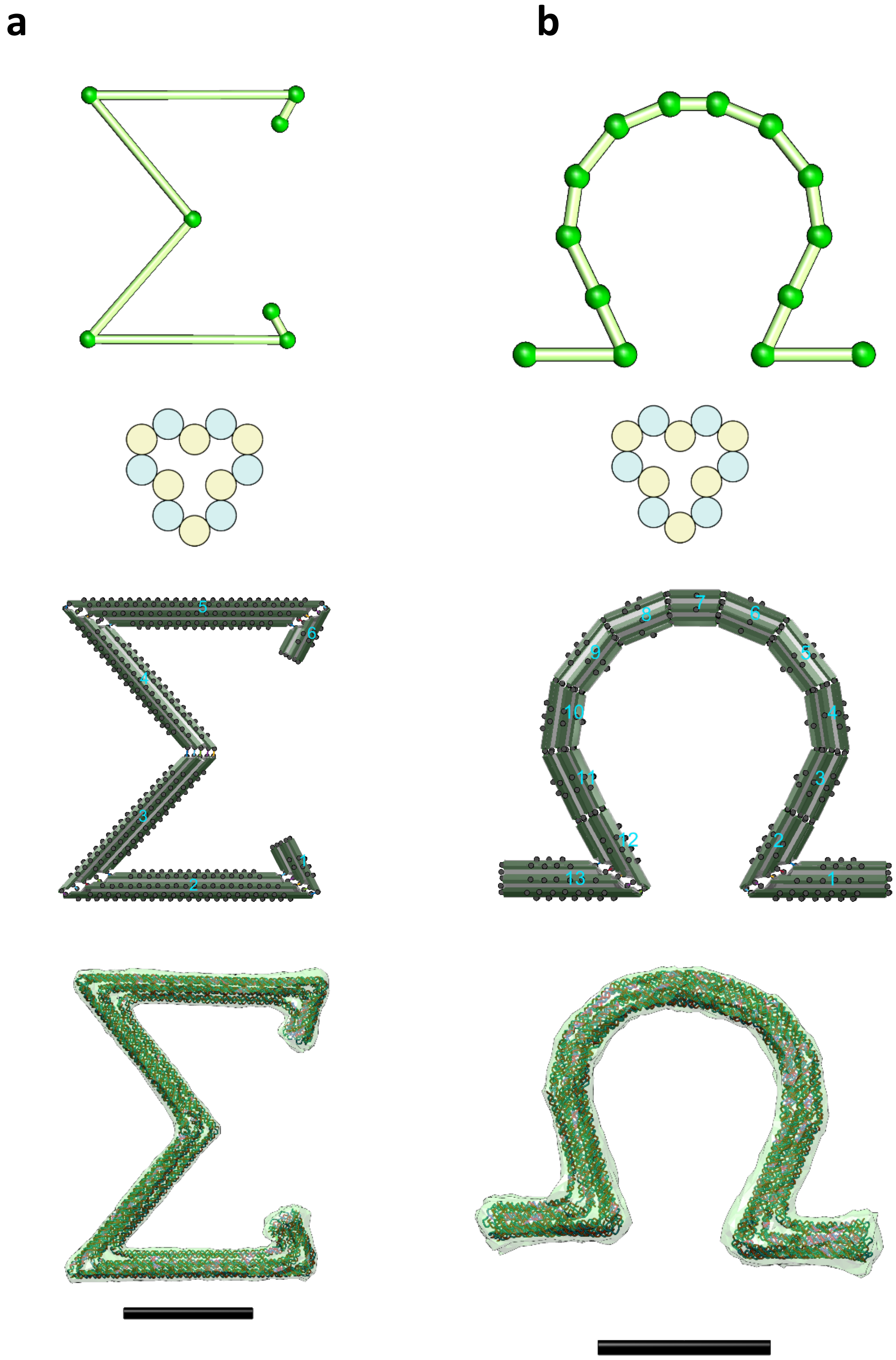


**Fig. S55.**

**Other freeform examples in the symbol series (sigma & omega).** (a) A design of "sigma". (b) A design of "omega". From Top to bottom, line sketch, cross-section, assembly model, and oxDNA simulation result. Scale bars = 50 nm.

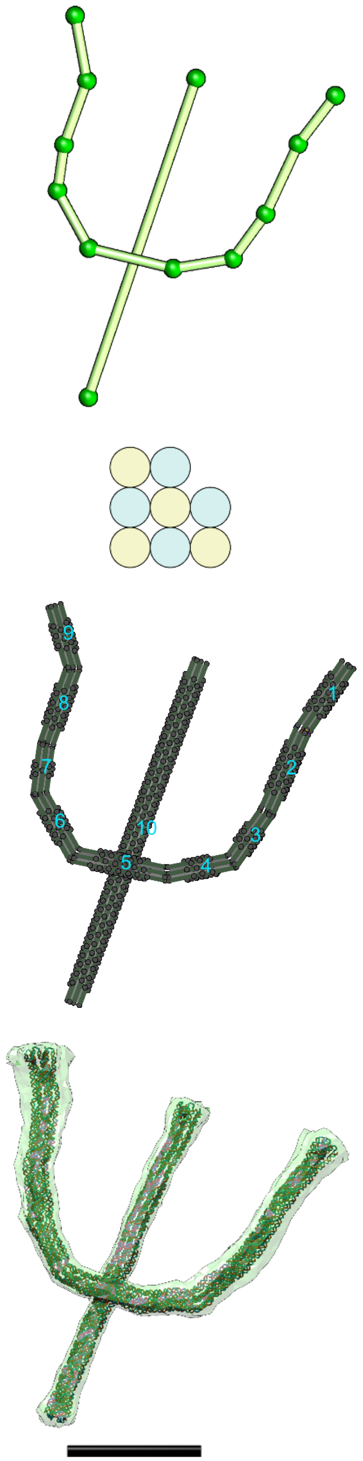


**Fig. S56.**

**Other freeform examples in the symbol series (psi).** A design of “psi”. From Top to bottom, line sketch, cross-section, assembly model, and oxDNA simulation result. Scale bars = 50 nm.

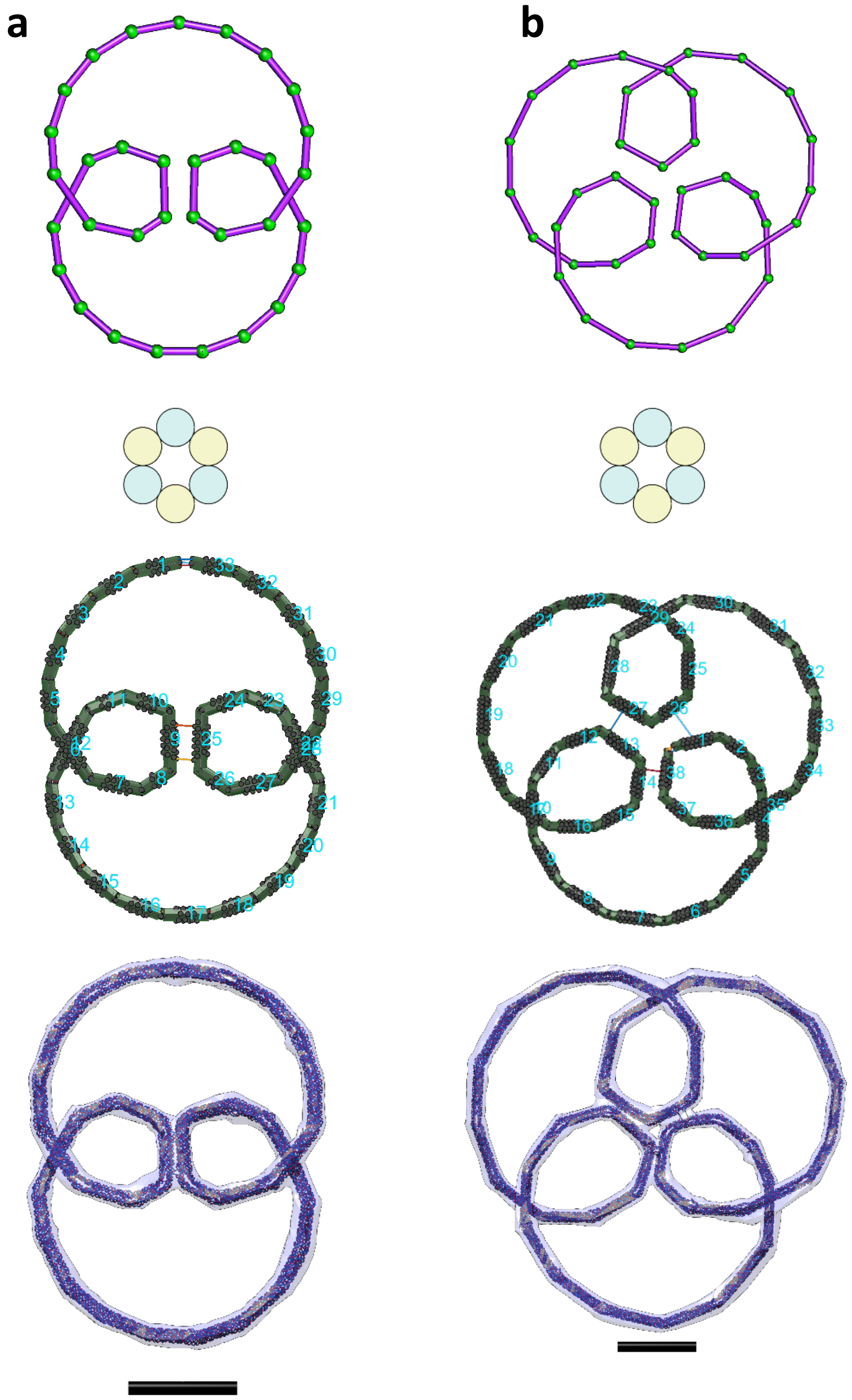


**Fig. S57.**
**Other freeform examples in the parametric curve series (I).** (a) An example design of the Hypotrochoid curve where the coefficients a=4, b=6, and c=2. (b) Another example design of the Hypotrochoid curve where the coefficients a=6, b=8, and c=2.2. From Top to bottom, line sketch, cross-section, assembly model, and oxDNA simulation result. Scale bars = 50 nm.

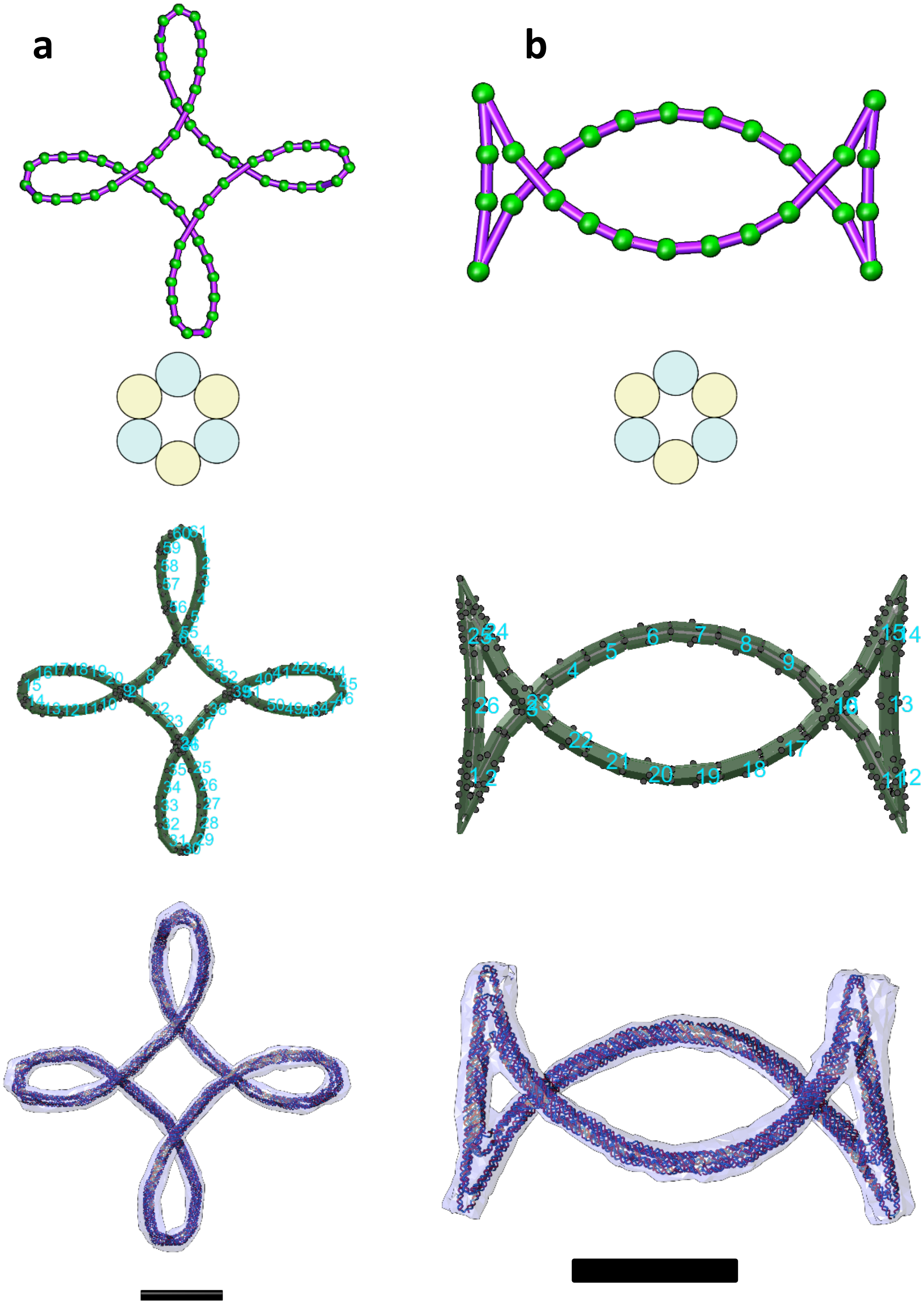


**Fig. S58.**

**Other freeform examples in the parametric curve series (II).** (a) The other example design of the Hypotrochoid curve where the coefficients a=1, b=3/4, and c=5/13. (b) An example design of Talbot's curve where the coefficients a=1.1, b=0.6, and f=1. From Top to bottom, line sketch, cross-section, assembly model, and oxDNA simulation result. Scale bars = 50 nm.

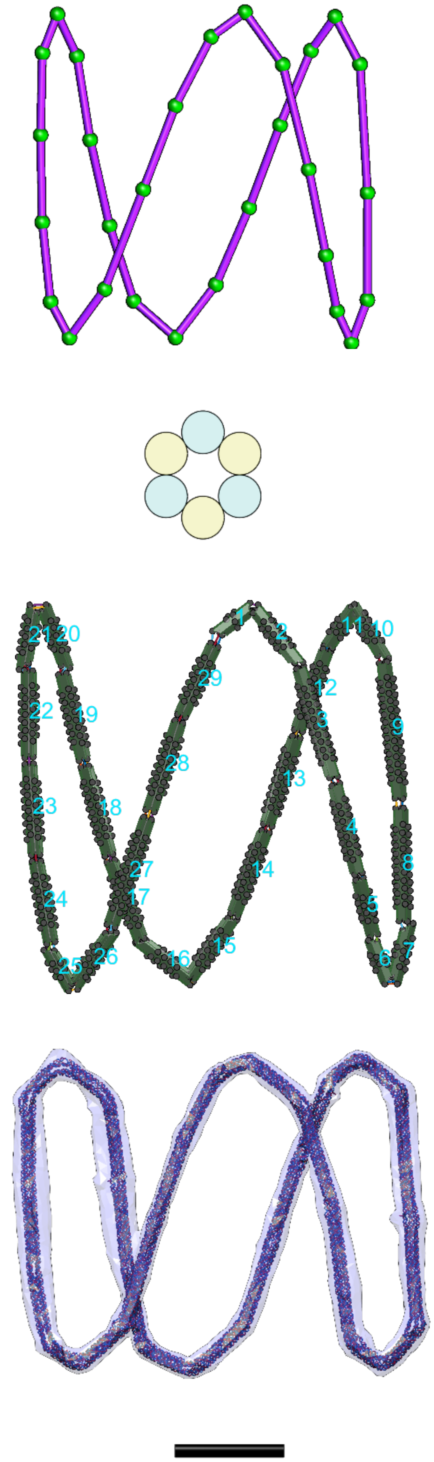

**Fig. S59.**

**Other freeform examples in the parametric curve series (III).** An example design of a Lissajous curve where the coefficients a=1, b=1, c=1, and n=3. From Top to bottom, line sketch, cross-section, assembly model, and oxDNA simulation result. Scale bar = 50 nm.

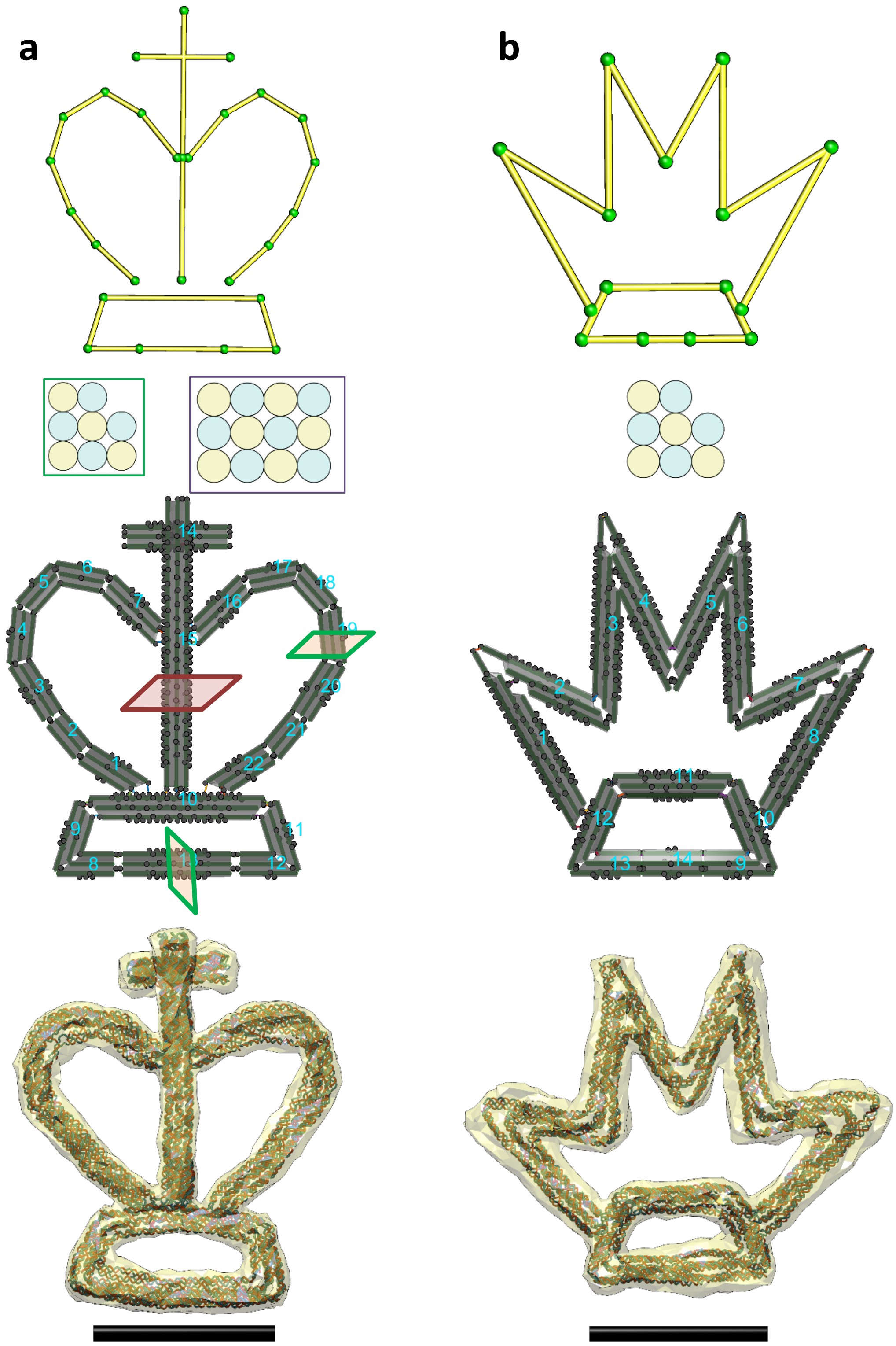


**Fig. S60.**

**Other freeform examples in the chess series (Queen & King).** (a) A design of "King". (b) A design of "Queen". From Top to bottom, line sketch, cross-section, assembly model, and oxDNA simulation result. Scale bars = 50 nm.

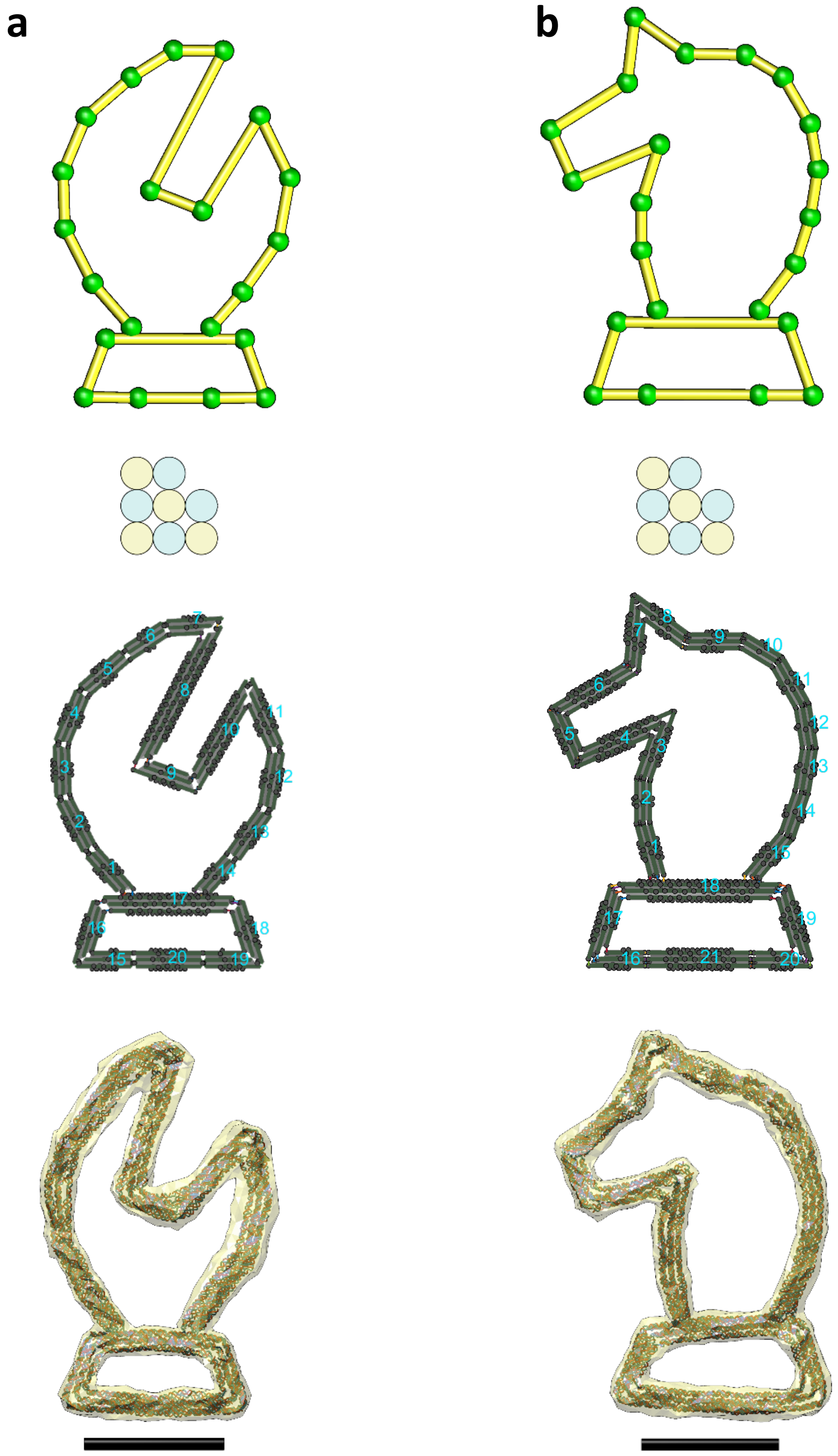


**Fig. S61.**

**Other freeform examples in the chess series (Bishop & Knight).** (a) A design of “Bishop”. (b) A design of “Knight”. From Top to bottom, line sketch, cross-section, assembly model, and oxDNA simulation result. Scale bars = 50 nm.

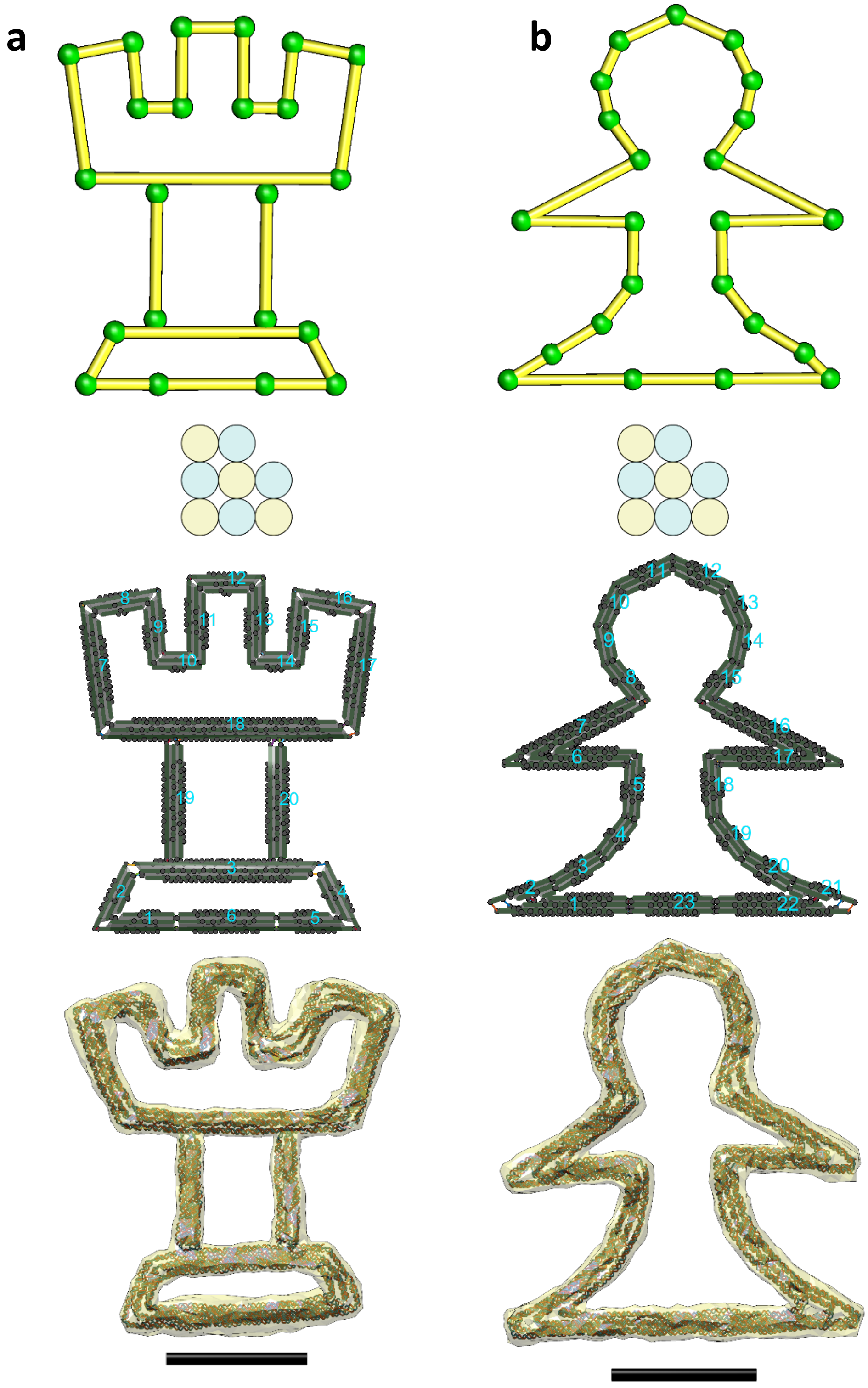


**Fig. S62.**

**Other freeform examples in the chess series (Rook & Pawn).** (a) A design of “Rook”. (b) A design of “Pawn”. From Top to bottom, line sketch, cross-section, assembly model, and oxDNA simulation result. Scale bars = 50 nm.

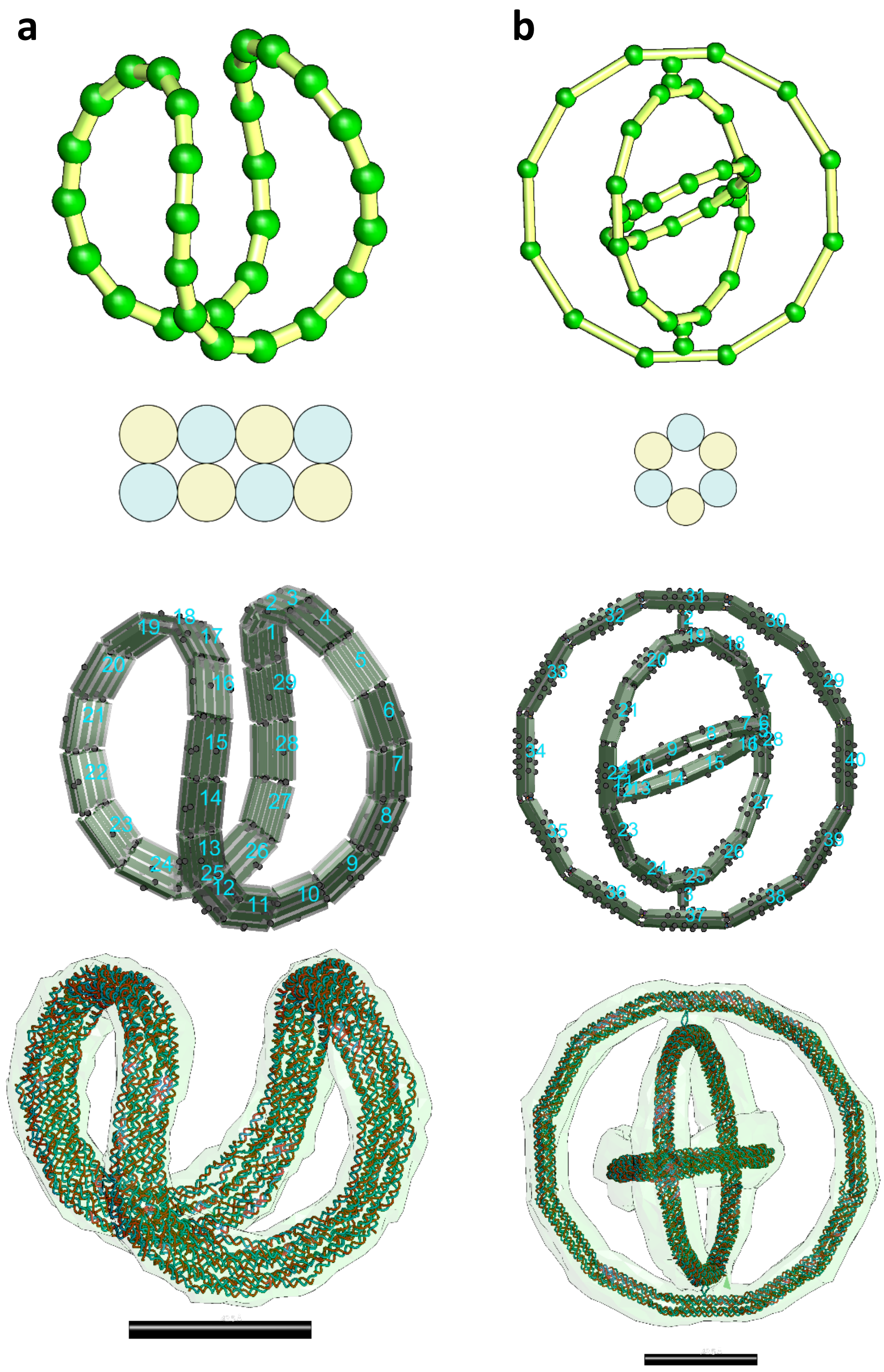


**Fig. S63.**

**Other freeform examples in the 3D series (Baseball-seam & Gyroscope).** (a) A design of the space curve which defines the seam of a baseball or tennis ball, where the coefficients a =7 and b=3. (b) A design of a three-layer Gyroscope. From Top to bottom, line sketch, cross-section, assembly model, and oxDNA simulation result. Scale bars = 50 nm.

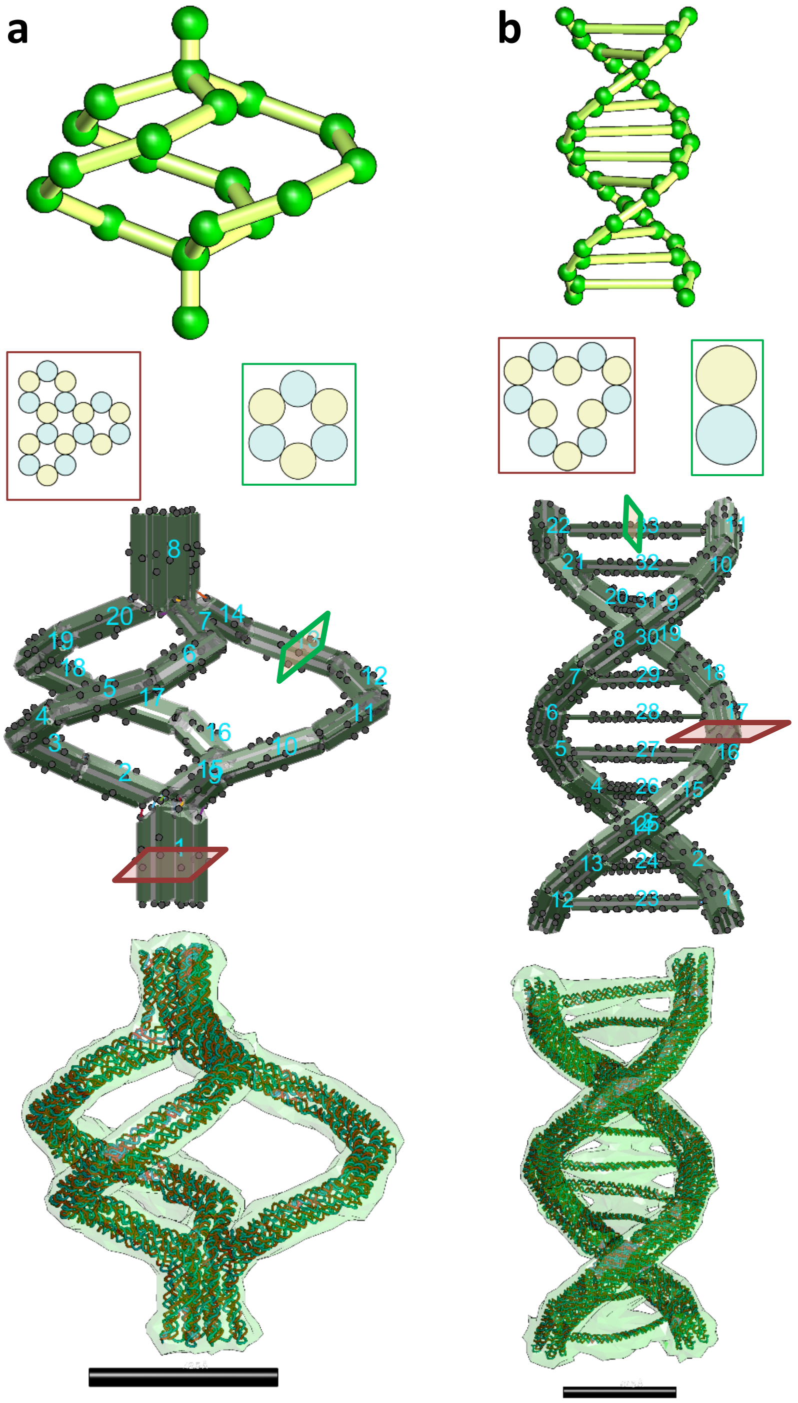


**Fig. S64.**

**Other freeform examples in the 3D series (Twist stair railing & Duplex DNA).** (a) A design of a twist stair railing. Note this design was involved the use of the bundle editing GUI to adjust the cylinder lengths in Bundles 1 and 8 (with 18HB cross-sections) to accommodate the geometries (edge gradients) of their three individual neighboring bundles (with 6HB cross-sections). (b) A design of DNA duplex. From Top to bottom, line sketch, cross-section, assembly model, and oxDNA simulation result. Scale bars = 50 nm.

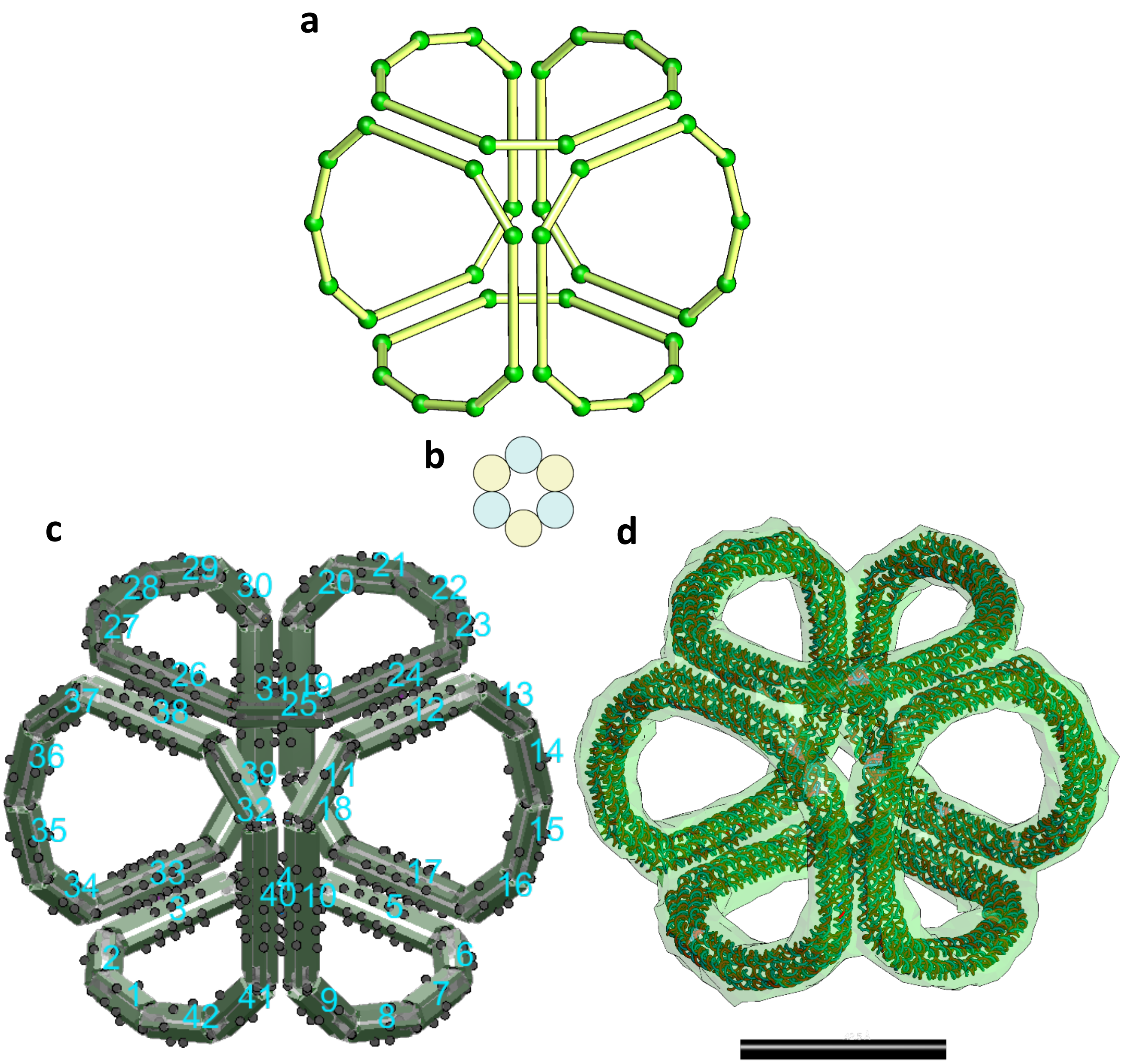


**Fig. S65.**

**Other freeform example in the 3D series (3D continuous curve).** A design of a 3D continuous curve which routes through the six faces of a cube [ref]. (a) line sketch, (b) cross-section, (c) assembly model, and (d) oxDNA simulation result. Scale bar =50 nm.
